# Mapping unsolved lipidomes accelerates lipid discovery in major bacterial pathogens

**DOI:** 10.1101/2025.11.06.685907

**Authors:** Yashodhan M. Nair, Aruna R. Menon, Zonghao Lin, Michiel R. L. Vossenberg, Vanisha Munsamy-Govender, David C. Young, Ana M. Xet-Mull, Gregory H. Babunovic, Tan-Yun Cheng, Sahadevan Raman, Kyu Y. Rhee, Jeremy M. Rock, Annemieke de Jong, Adriaan J. Minnaard, Jacob A. Mayfield, David M. Tobin, D. Branch Moody

## Abstract

Unlike gene-first approaches to understanding bacterial pathogenesis, molecule-forward discovery can uncover unexpected chemical diversity. Here, new lipidomic analytical methods and quality metrics defined the large scope of unknown lipids in the world’s deadliest pathogen, *Mycobacterium tuberculosis* (Mtb). This map allowed rapid discovery of Mtb lysyldiacylglycerol linked to the biosynthetic gene *lysX,* which controls *in vivo* infection outcomes in moth larvae, mice, guinea pigs, and here, zebrafish. A broader search for orthologous lysyltransferase domains identified the *Staphylococcus aureus* virulence gene *mprF*, where the same lipoamino acid was shown to be a previously unknown biosynthetic product. Thus, lipidomic mapping showed that the cell envelope composition of well-studied bacterial pathogens remains substantially unsolved and offers a new way to generate lists of discoverable lipids to accelerate molecular discovery.

## Main Text

Mtb’s success as the world’s deadliest human pathogen results from evasion and manipulation of human immunity (*1*). The interface between Mtb and the human host is formed by an extraordinarily complex cell envelope (Fig. 1A) (*2*). During infection, host nutrients and drugs must pass through two distinct phospholipid and neutral lipid membranes containing at least 22 known major lipid classes (*3*). Beyond barrier function, genus-specific lipids account for the unique biologic behavior of mycobacteria, including sulfoglycolipids (SGL) that induce cough (*4*), phthiocerol dimycocerosates (PDIM) that control drug entry (*5*) and immune evasion (*6*, *7*), mycobactins which scavenge host iron (*8*), ‘cord factor’ immunogens (*9*), as well as terpene nucleosides that block phagolysosomal acidification (*10*).

**Fig. 1.**
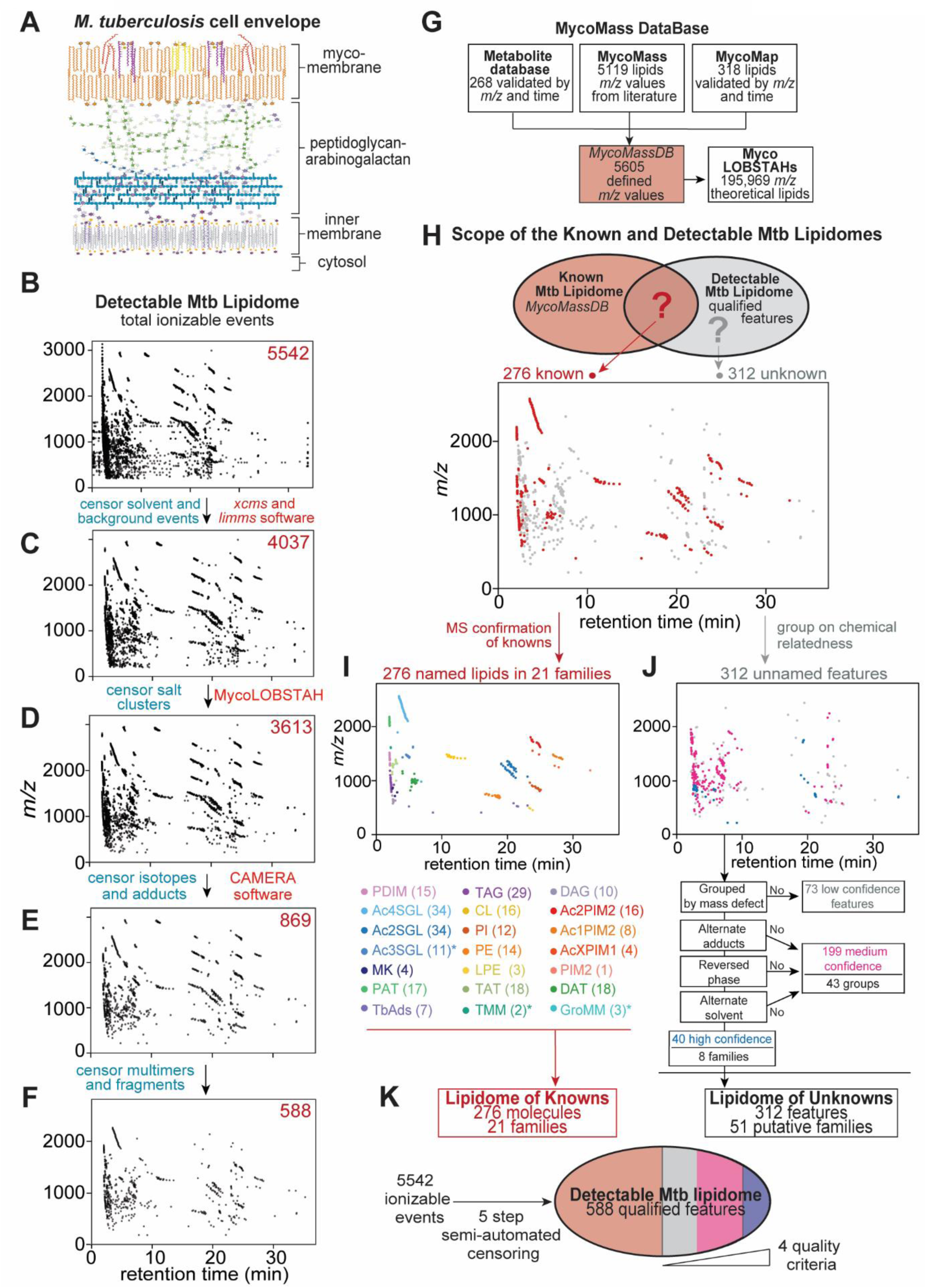
Mapping a lipidome of unknowns in *M. tuberculosis* (**A**) The Mtb cell envelope comprises two lipid membranes. (**B**) The 5542 features of the positive mode lipidome of Mtb H37Rv in biological quadruplicate underwent sequential, semi-automated censoring to remove (**C**) solvent ions, (**D**) salt clusters (**E**) isotopes and alternate adducts, and (**F**) in source multimers and fragments, yielding 588 credentialed features. (**G**) Source databases were updated and combined to create a metabolite and lipid database of defined *m/z* values (MycoMassDB), and a propagated database using LOBSTAHs (*21*) to generate theoretical lipid variants. (**H**) The 588 qualified features of the ‘detectable lipidome’ were classified as (**I**) 276 features of the ‘lipidome of knowns’, with lead compounds identified by collisional MS and (**J**) 312 features of the ‘lipidome of unknowns’ grouped into lipid families and ranked as high, medium, or low confidence ions based on four criteria: detectable acylforms, the presence of alternate adducts, detection in reversed phase chromatography, and alternate solvent extraction. (**K**) The lipidomes of detectable, credentialed known and unknown lipids were summarized. PDIM, phthiocerol dimycocerosates; Ac4SGL, tetra-acyl sulfoglycolipid; Ac3SGL, triacyl sulfoglycolipid; Ac2SGL, diacyl sulfoglycolipid; MK, menaquinone; PAT, polyacyl trehalose; TAT, triacyl trehalose; DAT, diacyl trehalose; DAG, diacylglycerol; TAG, triacylglycerol; CL, cardiolipin; PI, phosphatidylinositol; PE, phosphatidylethanolamine; LPE, lyso phosphatidylethanolamine; TMM, trehalose monomycolate; GroMM, glycerol monomycolate; Ac2PIM2, diacyl phosphatidylinositol dimannoside; Ac1PIM2 monoacyl phosphatidylinositol dimannoside; AcXPIM1, acyl phosphatidylinositol monomannoside; PIM2, phosphatidylinositol dimannoside. *Lipids with asterisks mass matched to MycoMassDB and were not studied by collisional MS.

Recognizing both high lipid complexity in the cell envelope and the incomplete functional annotation of the Mtb genome (*11*) we hypothesized that uncharacterized lipid virulence factors might exist in large numbers and their discovery could improve our understanding of tuberculosis (TB) disease. Indeed, recent studies have shown that abundant Mtb lipid families escaped discovery despite decades of intensive study of this major pathogen (*12*, *13*). Advances in mass spectrometry, combined with new software and databases for pathogenic bacteria (*3*, *14*) and viruses (*15*) now allow for rapid and comprehensive detection of a lipidome (*16*). This technology, which detects mass values for both known and unknown compounds, affords a new opportunity to exclude ions corresponding to known lipids and generate a ‘lipidome of unknowns.’ Such a lipid map might quantitatively circumscribe the scope of undiscovered molecules and accelerate their discovery by focusing on a defined list of targets.

### Qualifying the Mtb lipidome

We extracted total membrane lipids from matched quadruplicate cultures of Mtb strain H37Rv using chloroform and methanol, excluding glycans, proteins, and nucleic acids. These complex lipid mixtures entered a broadly separating, normal phase high performance liquid chromatography-electrospray ionization time of flight mass spectrometry system (HPLC-ESI-TOF-MS) optimized for high sensitivity and dynamic range (*3*). We first detected 5542 distinct ion traces known as molecular features, which are linked mass-to-charge ratio (*m/z*), retention time, and intensity values (Fig. 1B).

Detectable but unannotated features in any metabolome may represent valid undiscovered lipids, or MS artifacts from contaminants or redundant detection, where a single valid molecule may be represented as several ions due to modifications that shift the *m/z* (*3*). These modifications may be biological, such as natural ^13^C-isotopes, or non-biological, such as alternate charged adducts, multimers or fragments formed during electrospray ionization. Chemical investigation of the many observed ions without validation can derail untargeted discovery through considerable wasted effort (*17–20*). Therefore, we developed a software-based whole organism credentialing pipeline that rapidly appraises data quality to censor artifacts and redundancies, followed by ranking of targets to positively identify features with characteristics of genuine bacterial lipids for further investigation.

Contaminants and idiosyncratic signals were censored by comparison to solvent blanks (Fig. 1C). Salt clusters with low CH content were recognized and censored based on low mass defects using LOBSTAHs (*21*) (Fig. 1D). Alternate adducts of H^+^, NH_4_^+^, Na^+^ or ^13^C isotopes were censored using CAMERA (*22*) (Fig. 1E). *In source* lipid aggregates, with high confounding potential given their natural isotopes, and potential acylforms, required manual recognition of multimers of diacyltrehalose (DAT), phosphatidylinositol (PI), phosphatidylethanolamine (PE), phosphatidylinositol mannosides (PIMs) (fig. S1 A to C), recognizable glycerolipid fragments (fig. S1, D to G), and solvent-induced carbamate esters (*23*) (fig. S1, H to M), yielding 588 unique features.

This conservatively drawn map likely underestimated the true scope of the Mtb ‘pan-lipidome’, as it lacked molecules selectively made by other Mtb strains, or in different growth conditions, trace or unionizable molecules, valid lipids censored by these filters, and high mass lipids above the detection range (∼*m/z* 3200). Yet, we intended to determine a focused list as a starting point for molecular discovery with the potential for expansion. Notably, our yield of 588 chemically qualified features from 5542 total features (Fig. 1F) is consistent with previous observations of the high redundancy of detection in MS-based systems biology (*17*). These ∼600 well vetted targets begin to quantitatively circumscribe the Mtb lipidome, and might galvanize discovery approaches to Mtb, depending on what proportion is previously known.

### The unknown lipidome

Further analysis relied on clustering patterns seen in the retention time versus *m/z* plots (Fig. 1H). Considering lipidomics as a distinct sub-discipline of metabolomics, metabolite ions are distributed individually (*18*, *19*), but lipids cluster in groups, where each feature has the same underlying structure and displays the biological signatures of acylforms. Features within a lipid group display characteristic mass differences reflecting their natural biosynthesis in varying alkane (CH_2_, 14.0156 amu) or isoprenoid (C_5_H_8_, 68.0626 amu) series, unsaturation (H_2_, 2.0156 amu), and oxidation (O, 15.9949 amu) patterns. The emergence of clusters and the loss of isolated datapoints during sequential ion censoring (Fig. 1, B to F) served as a general validation of the credentialing pipeline, where artifacts are expected to appear as single ions but true lipids naturally cluster. Further leveraging grouping to visualize and analyze large datasets, we implemented a variant of mass defect filtering (*24*) whereby the Kendrick mass defect (*25*), defined as a mass co-efficient transformed by a methylene unit, classifies related acylforms on a horizontal axis (fig. S2, A to B). This made unsaturations, oxygenation, and other modifications apparent on inspection (fig. S2, B to M), and excluded inorganic molecules with low mass defect (fig. S2B).

Next we compared all qualified features to the largest available mycobacterial lipid database, MycoMass (*3*), which we further augmented with recently reported lipids and 268 validated mycobacterial metabolites to create the MycoMassDB as 5605 known molecules connected to *m/z* values that encompassed both lipidomic and metabolomic space (Fig. 1G, and Dataset S2). After initial analysis connected grouped *m/z* values to named compounds in MycoMassDB, we sought to directly validate matches with collisional MS. To simplify the large scope of effort needed to collide more than 200 matched compounds, we leveraged the phenomenon of lipid family clustering in mass-retention time space. We directly solved 18 lead compounds that defined 18 families, including PDIM, SGLs, acyltrehaloses (AcT), diacylglycerol (DAG), triacylglycerol (TAG), menaquinones (MK), cardiolipins (CL), phospholipids (PI, PE, PIMs) and terpene nucleosides (TbAds) (Fig. 1I and fig. S3, A to P). Overall, using the methylene transform tool for grouped clusters, we defined the known lipidome of Mtb H37Rv to 276 alkane or unsaturation variants (fig. S2).

Strikingly, 312 qualified features did not match known molecules from MycoMassDB (Fig. 1J), suggesting that unknown lipids might dominate in the Mtb lipidome. Recognizing that even a credentialed lipid list might contain residual artifacts or redundancies (*17*, *23*) we developed rigorous data quality metrics to rank unannotated signals (Fig. 1J). Unknown features were designated as high confidence based on 4 criteria, where ions showed 1.) methylene transformed mass defect grouping patterns indicating acylforms with natural carbon composition and variation; 2.) one molecule [M] as multiple adducts, suggesting a chemical functional group and lack of ionization-dependent fragments; 3.) identification in both reversed and normal phase chromatography, mitigating heteromultimers formed by chromatographic co-elution and shared ionization; and 4.) re-identification with a separate extraction in both chloroform/methanol, and substituted with methylene chloride or deuterated methanol respectively, excluding solvent-induced lipid modification (Fig. 1J, high confidence, *blue*).

154 features in 31 groups met between 1 and 3 and criteria and displayed some features of bacterial lipids, so were considered intermediate confidence and not prioritized for investigation (Fig. 1J, *magenta*). 73 ungrouped low confidence features were not pursued further due to the absence of a lipid alkane series, although they may still be genuine lipids without natural biosynthetic variants (Fig. 1J, *grey*). Overall, conservatively excluding these low and intermediate confidence features, our credentialing pipeline identified a core set of 8 previously unknown families consisting of 40 features that have the highest potential to be undiscovered lipids in this globally important pathogen (Fig. 1K).

### Discovery of lysyldiacylglycerol in Mtb

Next we investigated a cluster of seven high quality unnamed features eluting near 21 minutes (Fig. 2A), where monoisotopic *m/z* values differed by 14.015 and 2.015 *m/z*, indicating acylforms of a single underlying core structure. Colliding the C32:0 acylform (*m/z* 697.610) released lysine and diacylglycerol fragments (Fig. 2B and fig. S3Q) (*26*). Thus, we proposed a structure of lysyldiacylglycerol (lysylDAG) with *sn3* ester-linked lysine appearing as 7 acylforms in Mtb H37Rv (Fig. 2C). These lysine lipoamino acids were distinct from lysine-containing glycerophospholipids (*27*, *28*) based on the absence of phosphate. While these molecules corroborated descriptions in *Mycolicibacterium phlei* (*29*) and more recently in *Corynebacterium pseudotuberculosis* (*30*), they were previously unknown in Mtb or the genus *Mycobacterium*.

**Fig. 2.**
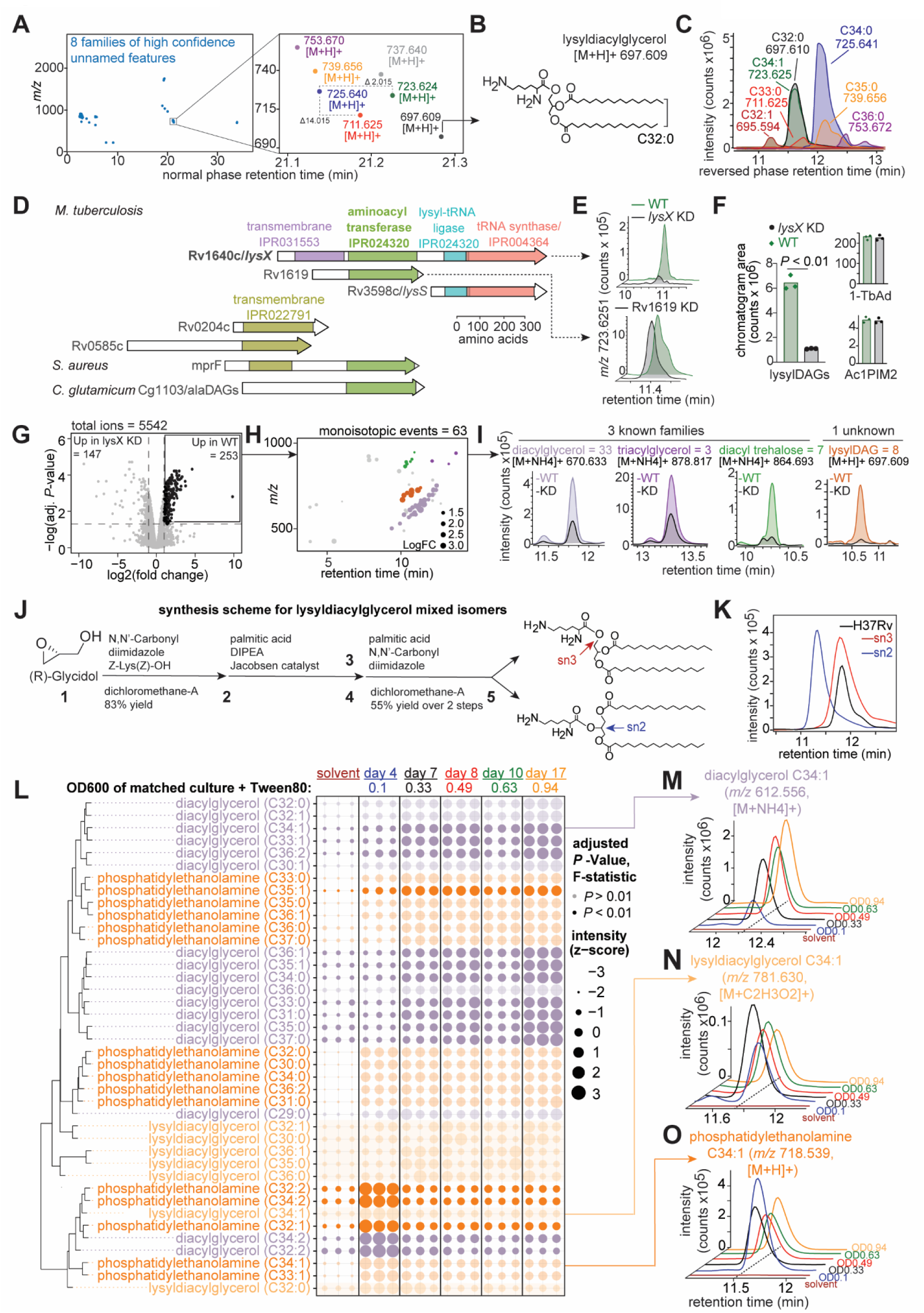
Identification of *lysX*-dependent lysyldiacylglycerol. (**A**) A cluster of high confidence (*blue*) features was shown to have 7 acylforms. (**B**) Collisional MS identified lysyldiacylglycerol (lysylDAG), with (**C**) representative mass chromatograms of acylforms in Mtb H37Rv. (**D**) A search for Mtb genes with homology to lysyltransferases by Interpro (*49*) subdomain structure identified two candidates, *lysX* and Rv1619. (**E**) Representative single ion chromatograms for the mass of lysylDAG (C34:1, [M+H]+) in total Mtb lipids from CRISPRi knockdowns of *lysX* and Rv1619 show *lysX*-dependence. (**F**) Chromatogram areas for summed lysylDAG acylforms, tuberculosinyl adenosine (*m/z* 540.355 [M+H]+, TbAd), and monoacyl phosphatidylinositol dimannoside (*m/z* 1432.942 [M+NH4]+, Ac1PIM2) analyzed in biological triplicate in the *lysX* KD background. *P*-value determined by pairwise *t*-test. (**G**) Comparative lipidome of *lysX* knockdown against the untreated control in biological triplicate, representative of two independent experiments. A significance threshold using the Benjamini-Hochberg adjusted *P*-value < 0.05 and two-fold change was used to identify *lysX*-dependent features. (**H**) 63 monoisotopic features were classified into three known and one unknown lipid families by grouping acylforms, *m/z* matching, and collisional MS. (**I**) Representative mass chromatograms of *lysX*-dependent families show the relative difference in intensity with *lysX* KD. (**J**) Both lysylDAG with lysine in the sn2 (*blue*) and sn3 (*red*) positions were synthesized in 5 steps. (**K**) Mass chromatograms of the C32:0 (*m/z* 697.609 [M+H]+) acylform of Mtb H37Rv lysylDAG (*black*) co-elutes in the reversed phase with the *sn3* synthetic lysylDAG (*red*) and not the *sn2* synthetic isomer (*blue*). (**L**) The z-score of the intensities of 7 alkylforms of lysylDAG, 17 diacylglycerol (DAG), and 16 phosphatidylethanolamine (PE) in Mtb H37Rv were measured across growth timepoints and plotted as scaled circles with a solvent only negative control. Distributions with a non-significant *P*-value of the F-statistic across all non-solvent pairwise timepoint contrasts, *P* > 0.01, are shown with 25% opacity. Representative of two independent experiments. Representative mass chromatograms show the distribution of (**M**) DAG, (**N**) lysylDAG and (**O**) PE across logarithmic to stationary growth timepoints sampled.

### Identification of *lysX* as the biosynthetic gene

Amino acyl phosphatidylglycerol synthases (aaPGs) canonically modify phospholipids with cationic amino acids (*31*), but might have a previously unrecognized function in modifying neutral lipids. For example, *Staphylococcus aureus* MprF catalyzes lysinylation of phosphatidylglycerol (PG) (*32*), but neutral lipid substrates have recently been identified (*33*, *34*). Mtb contains two paralogous aaPGs: Rv1640c (*lysX*) (*35*) and Rv1619 (*36*) (Fig. 2D). CRISPRi gene silencing (*37*) coupled to HPLC-MS demonstrated that all lysylDAG signals were significantly decreased by the silencing of *lysX* and not Rv1619 (Fig. 2, E and F). By contrast, unrelated lipids 1-tuberculosinyl adenosine (1-TbAd) and monoacyl phosphatidylinositol dimannoside (Ac1PIM2) were unaffected by silencing (Fig. 2F).

Although prior data linked *lysX* to lysyl phosphatidylglycerol (lysylPG), the known MprF product (*35*), the assignment was not validated by collisional MS or synthetic standards. Further, we could not detect the previously reported phospholipid ion (681.1 *m/z*). Also, another group found that lysylPG persists in *C. pseudotuberculosis* after *lysX* deletion (*30*). Thus, most data support the assignment of Mtb LysX as an amino acyl diacylglycerol synthase (aaDAGS) (*38*), rather than a glycerophospholipid modifying enzyme. LysX may be a lysine specific transferase that functionally resembles the corynebacterial enzyme AlaDAGS which catalyzes alanylation of DAG (*38*).

### Untargeted analysis of *lysX* mutants

Whereas targeted analysis measures a predetermined enzyme product, untargeted lipidomic analysis of *lysX*-silenced bacteria allowed hypothesis-free interrogation of all ionizable Mtb lipids as potential products. Indeed, silencing *lysX* significantly altered 253 of 5542 total features, representing a large lipidomic footprint beyond lysylDAG (Fig. 2G). Censoring isotopes, adducts and multimers, yielded 63 unique events, from which we identified 51 *lysX-*dependent lipids in 4 families (Fig. 2I). Three families were known compounds in MycoMassDB: 33 DAG, 7 DAT, and 3 TAG acylforms. These known lipids showed incomplete depletion (Fig. 2*I*, *left*), and their structures did not match expected products of an amino acyltransferase, suggesting their signals decreased due to indirect effects of precursor shunting or cell envelope remodeling in response to *lysX* depletion.

In contrast, a single family of eight lipids comprising lysylDAG showed nearly complete loss of MS signal (Fig. 2I, *right*), consistent with being products of the LysX enzyme. To enable structural confirmation by chromatographic co-elution and collisional patterns, we synthesized dipalmitic-lysylglycerol initially in two isomeric forms (Fig 2J, and fig. S4A). Mtb lysylDAG matched the *sn3* isomer in chromatographic retention time (Fig. 2K), mass (fig. S4B), and collisional MS spectra (fig. S4, C and D), demonstrating the complete chemical structure of the natural product. Thereafter, we developed a synthesis method for isomerically pure sn3-lysylDAG (fig. S4E).

To test whether lysylDAGs were consistent cell envelope components, we extracted lipids at 5 timepoints across Mtb H37Rv logarithmic to stationary growth phases in media without supplemental oleate or dextrose (fig. S5, A and B). We measured 7 acylforms of lysylDAG, 17 DAG, and 16 PE and analyzed their distributions (Fig. 2L). Next we used all non-solvent pairwise comparisons (fig. S5C) to identify significant (adjusted *P*-value < 0.01 of the F-statistic) lipid variation across growth phases. We observed similar levels of lysylDAG across all timepoints (Fig 2L). This distribution was similar to acylforms of the unrelated glycerolipid PE, and in contrast to the precursor neutral lipid DAG which accumulated in the Mtb cell envelope in liquid culture across growth phases (Fig. 2, M to O). Similarly, in Sauton’s media, where glycerol is the sole carbon source, with and without supplemental lysine, we detected consistent lysylDAG features across 4 growth timepoints, like distributions of PE and DAG (fig. S5, D to I). Thus, lysylDAG biosynthesis was a stable feature of the Mtb cell envelope produced constitutively throughout growth stages, independent of typical precursor flux, media composition, and exogenous lysine.

Lysinylated lipids have a putative mechanistic function in controlling the charge state of the envelope (*31*). Unlike lysinylated phospholipids that carry an anionic phosphate, lysylDAG has stronger cationic potential that could account for the role of Mtb *lysX* in maintaining Mtb resting membrane potential, resistance to cationic antibiotics vancomycin and polymyxin-B (*35*), and broader resistance to treatment-relevant antibiotics such as rifampicin and isoniazid (*39*). Overall, an untargeted ‘lipid-first’ search of the unknown lipidome here identified the downstream metabolite that likely mediates the role of the *lysX* in Mtb virulence and antimicrobial resistance, and expands MycoMassDB by 1 new major class and 30 possible acylforms for future interpretation of altered cell envelope composition by untargeted lipidomics.

### Mapping mycobacterial virulence lipids

To determine the species distribution of lysylDAG we reconstructed the phylogeny of major aminoacyl transferases, demonstrating the presence of *lysX* across both disease-causing and environmental mycobacteria (Fig. 3A), consistent with *lysX* as a core mycobacterial gene. We selected representative species across the Mtb complex of highly genetically similar tuberculosis-causing bacteria with different host tropisms, non-tuberculous opportunistic pathogenic mycobacteria, and non-pathogenic environmental mycobacteria, and extracted lipids with equivalent growth conditions in biological quadruplicate. Targeted HPLC-MS lipid analysis of 3 Mtb strains and 12 mycobacterial species identified major C34 acylforms of lysylDAG in every mycobacterium tested with species-specific saturation patterns (Fig. 3B).

**Fig. 3.**
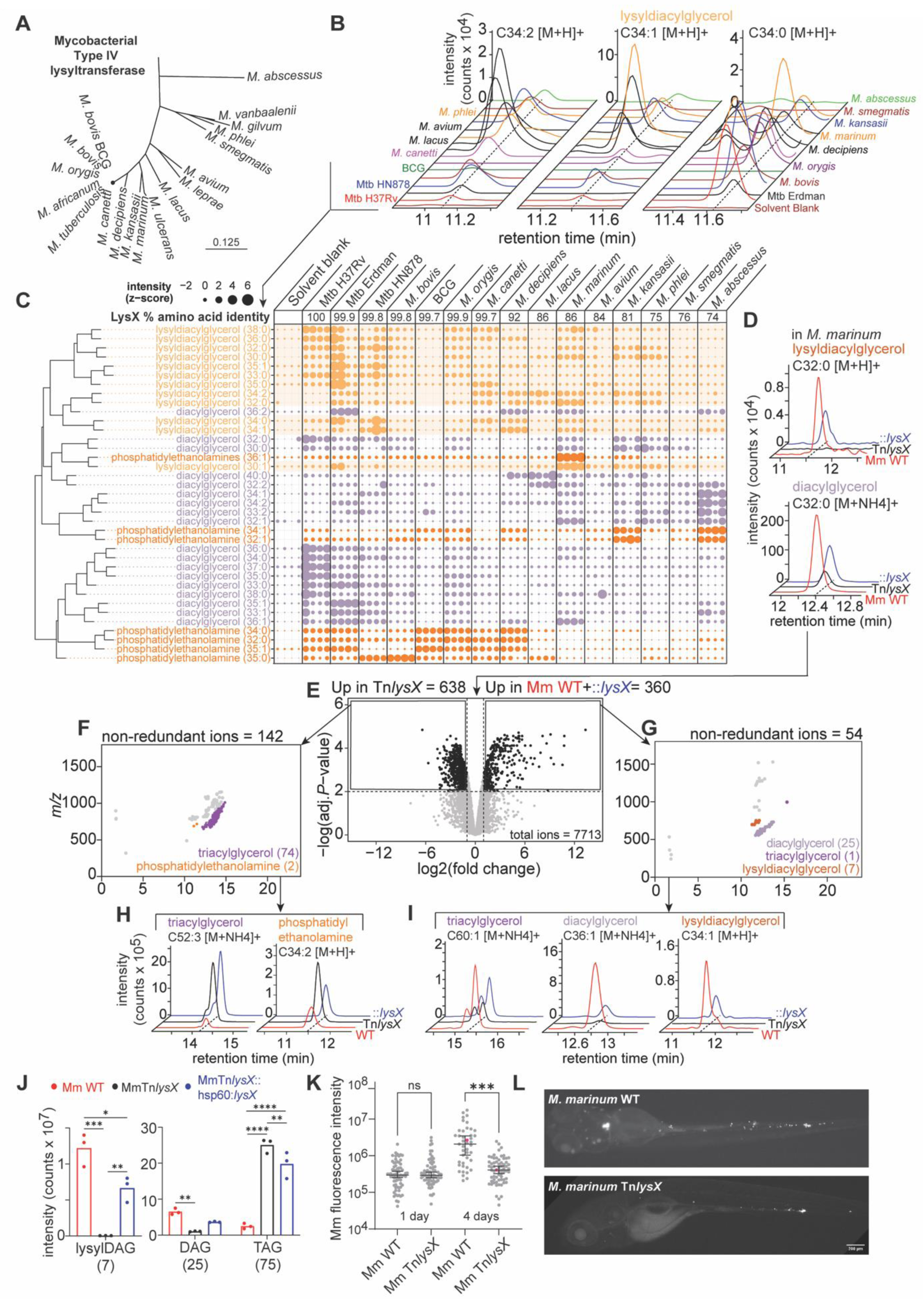
Lysyldiacylglycerol-*lysX*: a conserved mycobacterial virulence pair. (**A**) Subset of a phylogenetic tree of LysX amino acid sequence shows the conservation of *lysX* across mycobacteria, with MprF-domain enzyme classification described previously (*38*). (**B**) Mass chromatograms show the distribution of representative lysyldiacylglycerol (lysylDAG) acylforms across mycobacteria profiled. (**C**) The z-score intensities of acylforms of lysylDAG, diacylglycerol (DAG), and phosphatidylethanolamine (PE) were plotted as scaled circles in biological quadruplicate lipid extracts from 3 Mtb strains and 12 mycobacteria grown in parallel. (**D**) Mass chromatograms of lysylDAG and DAG in *M. marinum* (Mm) WT, *lysX* transposon mutant (Tn*lysX*) and *lysX* complement (Tn*lysX*::hsp60::*lysX*) show *lysX* dependence. (**E**) A threshold of two-fold change and *P*-value < 0.01 in a compound lipidomic contrast of Mm WT and *lysX* complement against Tn*lysX* identified lipids significantly changed by *lysX* disruption. Strains were grown in biological triplicate, representative of two independent experiments. (**F**) 142 monoisotopic features were enriched in Mm Tn*lysX*. Grouping and lead compound collisional MS identified 74 acylforms of triacylglycerol (TAG) and 2 acylforms of phosphatidylethanolamine (PE). (**G**) 72 monoisotopic features were enriched in Mm WT and ::*lysX*. Grouping and lead compound collisional-MS identified 25 acylforms of DAG, 1 of TAG, and 7 of lysylDAG. Representative single ion chromatograms of (**H**) PE and TAG show enrichment in both Tn*lysX* (*black*) and ::*lysX* (*blue*), whereas (**I**) DAG, lysylDAG, and TAG show enrichment in Mm WT (*red*) and ::*lysX* (*blue*). lysylDAG was absent in the *lysX* mutant. (**J**) Chromatographic areas in the Mm *lysX* lipidome were summed and significant differences were evaluated by two-way ANOVA with Tukey’s post-test (*: *P* < 0.05; **: *P* < 0.01; ***: *P* < 0.001; ****: *P* < 0.0001). (**K**) *M. marinum* burden in zebrafish infection measured using bacterial mCerulean fluorescence. Data shows one representative experiment of 3 biological replicates with 30-60 independent infections per replicate. Median and 95% confidence interval are displayed. Statistical analyses were performed using one-way Welch’s ANOVA followed by a Dunnett’s T3 multiple comparison test of each group to the WT strain (ns: *P* > 0.05; **: *P* ≤ 0.01; ***: *P* ≤ 0.001; ****: *P* < 0.0001). (**L**) Representative images from zebrafish, depicted in (K) in pink, infected with an initial dose of 150-200 fluorescence units of either WT or Tn*lysX* at 4 days post infection.

While an individual lipid family may be conserved, acylform preferences (*40*) may hamper attempts to catalogue lipids across species by a single *m/z*. So, we analyzed the acylform distributions of glycerolipids by assessing their relative intensities, using a custom function *mzrtMeta* to align peaks across lipidomic experiments (Materials and Methods). All mycobacteria produced lysylDAG with species-specific acylform patterns. Consistent with PE and DAG, the Mtb complex preferred fully saturated variants of lysylDAG with predominance of the C34:0, C35:0 and C36:0 acylforms. By contrast, environmental and non-tuberculous mycobacteria preferred unsaturated acylforms such as C34:1 and C34:2, that may be consistent with temperature adaptation of cell envelope components to the endothermic host by the Mtb complex common ancestor or the *cfa*-dependent production of tuberculostearic acid from oleic acid by some mycobacteria (*41*). Therefore, supporting a significant biological role, both *lysX* and lysyldiacylglycerol are broadly conserved across the family Mycobacteriaceae, which includes Mtb and other clinically important non-tuberculous mycobacteria such as *Mycobacterioides abscessus*. In contrast, we also identified reduced but detectable lysyldiacylglycerol levels in the live attenuated vaccine strain Bacille-Calmette Guerin (BCG), the lowest producer of lysyldiacylglycerol across all mycobacteria tested. Moreover, deletion of *lysX*, which is shown here to disrupt lysyldiacylglycerol, previously attenuated Mtb growth *in vivo* or host survival in mice (*35*), guinea pigs (*42*) and *C. pseudotuberculosis* outcomes in wax moth larvae (*30*) in experiments performed with genetic complementation. Thus, the *lysX*-lysylDAG pathway identified here has a non-redundant role in promoting pathogen survival or virulence across evolutionarily diverse actinobacteria-host pairs.

### A *lysX*-lysyldiacylglycerol pathway in *M. marinum*

Beyond demonstrating the presence of lysylDAG in tuberculous and non-tuberculous mycobacteria, phylogenetic analysis showed relatively high production of lysylDAG in *M. marinum* (Mm) (Fig. 3C), a natural pathogen of ectotherms. Mm infection shares key conserved features of tuberculosis pathogenesis that are directly observable in a transparent zebrafish host (*43*). After generating a *lysX* mutant (Mm:Tn*lysX*) in the candidate orthologue (MMAR_2247) with 86% sequence identity to Mtb LysX (Fig. 3C), we observed loss of lysylDAG, which was restored with complementation (Mm:Tn*lysX*::hsp60::*lysX*) (Fig. 3D). These data established the existence of a lysylDAG pathway in a tractable fish pathogenesis model, setting the stage for whole organism lipidomics and *in vivo* infection experiments.

Untargeted comparative lipidomics of Mm:Tn*<u>lysX</u>* against the wild type (WT) and complemented strains identified ∼13% of the lipidome significantly altered by *lysX* disruption (two-fold change, *P* < 0.01) (Fig. 3E). Among 142 non-redundant features enriched in the mutant (Fig 3E), we identified 74 TAG and 2 PE alkylforms, noting a pattern of enrichment in unsaturated acylforms (Fig 3G and fig. S6, A and B). Among 54 non-redundant features reduced in the deletion mutant, 7 lysylDAG acylforms showed complete loss and restoration with complementation. Polar or second site effects were further excluded as disruption of the potential co-operonic downstream gene Rv1639c/MMAR_2446 had no significant effect on lysylDAG levels (fig. S6F). We also noted partial loss of 25 acylforms of DAG and 1 of TAG, which were restored with complementation (Figure 3H and fig. S6C to E).

Despite the untargeted nature and broad scale of the two analyses in Mtb and Mm, the lipidomic patterns of change after *lysX* disruption were strikingly similar, leading to general conclusions and new questions. First, the complete loss of all acylforms of lysylDAG and its restoration by genetic complementation were most consistent with *lysX* acting as the sole biosynthetic enzyme to catalyze these pathways in both Mtb and Mm. Second, the emergence of changes in DAG and TAG in similar numbers and fold-change parameters in both pathogens represents a highly penetrant and reproducible effect of *lysX* inactivation on neutral lipids other than lysylDAG. While loss of DAG lysinylation through *lysX* deletion predicts accumulation of DAG as a substrate, we observed the opposite. Further, DAG and TAG changed in opposite directions, which, given their usual relationship, predicts DAG to TAG conversion by an unknown mechanism as a means of energy storage (*44*). *lysX*’s conserved and unexpectedly broad role in governing mycobacterial cell envelope neutral lipid composition suggests an indirect or secondary function beyond that of lysinylation that may contribute to its role in virulence, here uncovered by an untargeted analysis of changes in all detectable downstream lipids following *lysX* disruption independently in pathogenic mycobacteria.

### *lysX* effects on *in vivo* infection of zebrafish

Taking advantage of the tractable and optically accessible nature of zebrafish, we infected larval zebrafish with either wildtype or *lysX* mutant Mm constitutively expressing cerulean fluorescent protein (Fig. 3J). Bacteria from both strains were phagocytosed normally by macrophages within the first hours of infection and resided intracellularly throughout the course of the infection. By 4 days, we observed significant attenuation for the *lysX* mutants deficient in lysylDAG (Fig. 3, J and K), suggesting a role for *lysX* in promoting Mm infection *in vivo*. Along with studies of guinea pig (*42*), mouse (*35*) and moth larvae (*30*), these data provide clear evidence for both the presence and functional importance of *lysX* and its products during infection of evolutionarily diverse hosts.

### Identifying *S. aureus* lysyldiacylglycerol

LysX is here shown to lysinylate neutral lipids. Such type IV aminoacyl transferases are found throughout Actinobacteria (Fig. 3*A*) (*38*). In contrast, two types of related enzymes catalyze other aminoacyl transferase reactions. The corynebacterial Type VII AlaDAGS transfers alanine to neutral lipids. The firmicute Type II enzyme MprF is a lysyltransferase classically producing the abundant firmicute virulence lipid, lysylPG, a phospholipid that mediates broad staphylococcal resistance to cationic antibiotics and antimicrobial peptides (*31*). To assess possible shared orthology of *lysX*, *mprF,* and *alaDAGS* genes, we constructed a phylogeny by amino acid sequence (*38*) (Fig. 4A). We mapped *lysX* orthologues beyond mycobacteria as in Figure 3A, defined by a unique transmembrane domain IPR031553 (Fig. 2D), to other actinobacteria including *Rhodococcus*, *Gordonia*, and pathogenic corynebacteria including *Corynebacterium diphtheriae*.

**Fig. 4.**
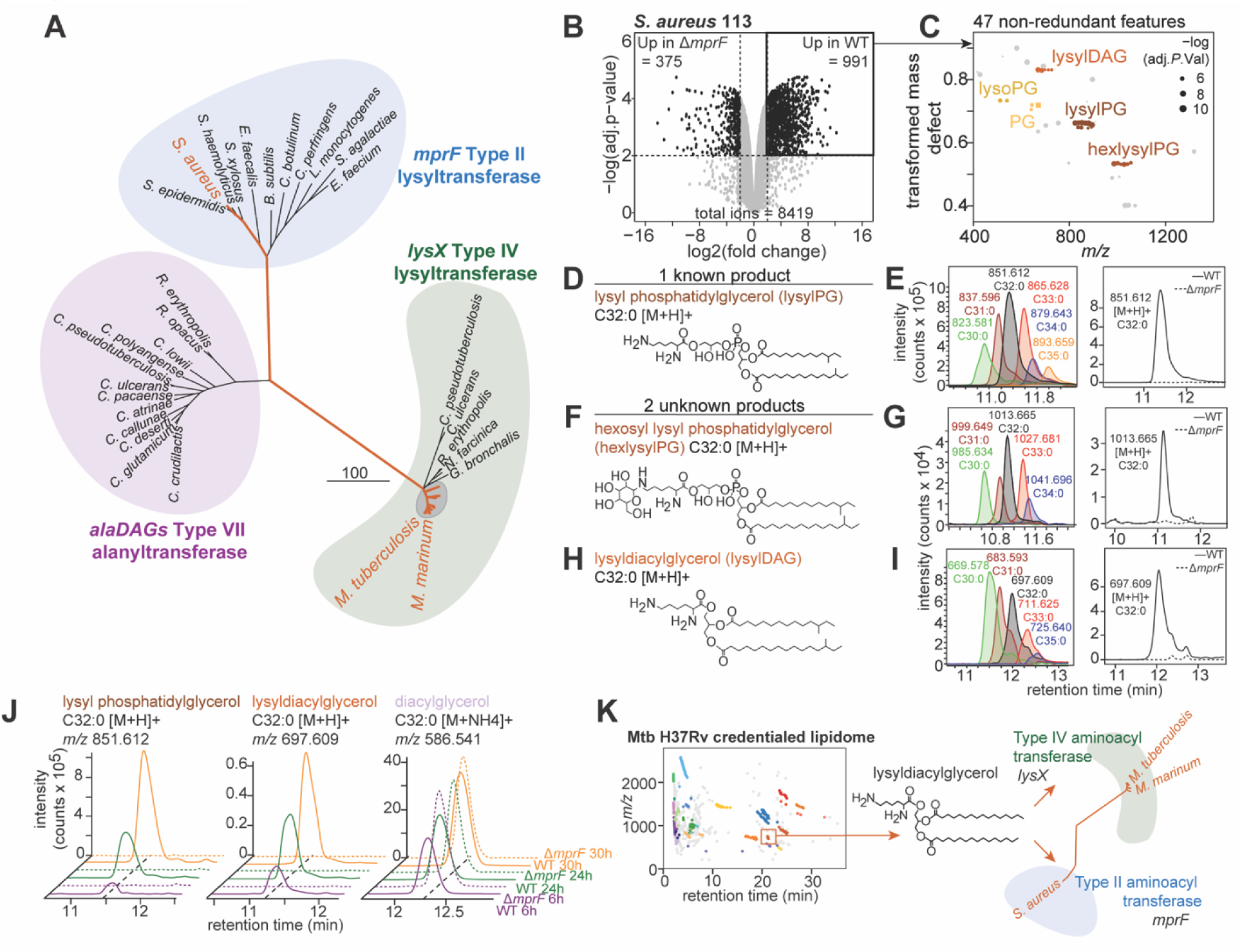
Lysyldiacylglycerol in firmicutes. (**A**) Amino acid sequence phylogeny of major MprF-domain containing proteins in actinobacteria and firmicutes show three clades. (**B**) A comparative lipidome of *S. aureus* 113 WT against the Δ*mprF* mutant was used to identify *mprF*-dependent features in biological triplicate (**C**) Grouping by transformed mass defect, 5 families with > 2-fold change were enriched in the S. aureus WT strain, Benjamini-Hochberg adjusted *P* value < 0.01. (**D**) One family consistent with a known product was identified as lysyl phosphatidylglycerol (lysylPG). (**E**) Representative chromatograms show the lysylPG acylform distribution *S. aureus* WT, *left*, and mass chromatogram showing absence in the *mprF* mutant, *right*. (**F** and **H**) Two previously unknown families were identified by collisional MS as an (*F*) hexosyl-modified lysylPG (hexlysylPG) and (**H**) lysyldiacylglycerol (lysylDAG). (**G** and **I**) Representative mass chromatograms show the acylform distribution of hexlysylPG and lysylDAG in *S. aureus* WT, *left*, and mass chromatogram showing absence in the *mprF* mutant, *right,* consistent with MprF products, and (**J**) show detection of lysylPG, lysylDAG, and DAG across growth phases. (**K**) A summary figure maps the identification of lysylDAG in the ‘lipidome of unknowns’ of Mtb H37Rv, extending this discovery to representative mycobacteria dependent on *lysX* and to *S. aureus* dependent on *mprF*.

Recognizing the shared lysyltransferase activity of LysX and MprF, and because recent lipid discovery efforts have identified additional MprF products of neutral lipids such as lysyl glucosyl diacylglycerol in *Streptococcus agalactiae* (*34*) and lysyl diglucosyl diacylglycerol in *Enterococci* (*33*), we hypothesized that substrate diversity of the MprF enzyme family might be broader than currently known, where undiscovered *mprF*-dependent products might be discoverable by lipidomics tools. We focused on *Staphylococcus aureus*, which is major Gram-positive pathogen that remains a leading cause of death from bloodstream infections and a major source of drug-resistant bacteria in hospital settings (*45*). Hypothesizing a broader distribution of lysine-modified neutral glycerolipids, we investigated the unsupervised *mprF*-dependent lipidome of *S. aureus*.

Contrasting the *S. aureus* SA113 lipidome against the Δ*mprF* mutant (Fig. 4B), we identified 47 features significantly enriched in the WT strain (Fig. 4C). We used the methylene transformed mass defect tool to identify five lipid families, of which 3 displayed complete signal dependence on *mprF.* First, one of these families was identified as the known *mprF*-product lysylPG both in accurate mass, collisional spectra, and a previously reported acylform preference for odd-chain lipids in staphylococci (*46*) (Fig. 4, D and E and fig. S7A). Notably, we identified two additional families of lipids with complete *mprF*-dependence and transformed mass defects characteristic of differing oxygenation states (Fig. 4C). Collisional MS identified one of these new families as lysylPG with a hexosyl modification, where neutral loss of 162.050 *m/z* was present in both positive and negative modes (Fig 4F, and fig. S7B). The acylform distribution of this glycerophospholipid in *S. aureus* mirrored that of lysylPG consistent with the derivation therefrom (Fig. 4G). Fulfilling the original hypothesis, we identified a third *mprF*-dependent family as staphylococcal lysylDAG with diagnostic fragments for lysine and neutral losses consistent with odd chain fatty acids (Fig. 4H and fig. S7C). The acylform diversity was distinct from both lysylPG and hexosyl lysylPG characterized here (Fig. 4*I*), consistent with synthesis from an alternate substrate pool.

To test whether lysylDAG was a consistent feature of the *S. aureus* cell envelope we investigated an *S. aureus* time course dependent lipidome by extracting lipids from both WT and Δ*mprF* mutant at three timepoints and plotting relative lipid intensity by z-score for eight acylforms of lysylPG, 8 PG, 3 lysylDAG and 6 DAG (fig. S7, D to F). Both lysylPG and lysylDAG were consistently detected across all 3 timepoints exclusively in the WT strain in an *mprF*-dependent manner (Fig. 4J). In contrast the neutral precursor, DAG, showed a similar distribution across timepoints and strains (Fig. 4J and fig. S7F). Thus, using both chemical homologies along with gene orthology to drive discovery, this untargeted discovery process extended identification of lysylDAG from Mtb and Actinobacteria to the well-characterized Gram-positive staphylococcal virulence factor *mprF*. Overall, the taxonomic distribution of lysylDAG here identifies a previously unrecognized, broadly distributed component of pathogenic bacterial cell envelopes, dependent on two virulence genes *lysX* and *mprF*, implicated in resistance to antibiotics and human innate immunity.

### Leveraging untargeted lipid discovery

A major limitation of MS-based discovery is that metabolites generate artifactual or redundant signals (*17*, *18*, *47*). We quantitatively validate this perspective by showing that in raw Mtb datasets such errors exceed the number of biologically relevant molecules tenfold. Yet, a step-wise qualification process defined specific sources of error and estimated their frequency to pinpoint ∼300 unnamed mass-defined features that constitute a ‘lipidome of unknowns.’ A caveat is that we substantially leveraged grouped acylforms as a biosignature of natural molecules that is detectable in high throughput. Thus, while this new approach is applicable to lipids from any source, it may not apply to non-lipid metabolites without repeating mass intervals. Overall, the process of mapping unsolved lipids supports several conclusions.

The search for new molecules is motivated by the expectation that they exist. Arguing against the perspective that decades-long research has already mined the biologically meaningful lipids in Mtb, by extending molecular credentialing efforts (*20*, *48*) we demonstrated that the Mtb lipidome is substantially unsolved. Molecular mapping identified a core set of 40 undiscovered lipids in 8 families that now represent specific high value targets. The first discovery outcome of this process identified lysine-modified neutral lipids produced by both mycobacterial *lysX* and staphylococcal *mprF*, which have broad virulence effects across two bacterial phyla responsible for major global diseases (Figure 4K). Beyond lipoamino acids, by combining genetic manipulation with organism-wide lipidomics we allow unknown downstream chemical mediators of disease to enter studies of bacterial pathogenesis, addressing a mechanistic blind-spot in gene-first analysis. In modern genomics, the complete catalogue of genes is defined, and investigation prioritizes functional phenotyping. By contrast, the scope of unknown metabolites in any system is fundamentally undefined. Here, we argue that sensitive detection of mass-identified unknowns, followed by systematic credentialing and triage, can focus and advance discovery efforts to biologically meaningful outcomes. Finally, by developing an approach to map the ‘dark matter’ of the lipidome, we chart a course to discover further unknown bacterial lipids that mediate diseases which continue to bedevil mankind.

## Acknowledgements

We thank Marcel Behr for providing *M. orygis* and *M. bovis*, Mark Sullivan and Eric Rubin for *M. abscessus*, Hafid Soualhine and Christine Turenne for *M. lacus*, Barbara Brown-Elliott for *M. decipiens*, Andreas Peschel for *S. aureus* and the Δ*mprF* mutant, Catherine Vilchèze and William R. Jacobs Jr. for the Mtb auxotrophic strain mc^2^7902, and Christine Cosma and Lalita Ramakrishnan for use of the *M. marinum* transposon library.

## Author contributions

Conceptualization: YMN, DBM, DCY, TYC

Methodology: YMN, ARM, ZL, MRLV, VMG, DCY, AMXM, GHB, TYC, SR, JMR, ADJ, AJM, JAM Investigation: YMN, AM, ZL, MRLV, VMG, AMXM, DCY, TYC

Visualization: YMN, DCY, ARM Funding acquisition: DBM, DMT, ADJ

Writing – original draft: YMN, DBM

Writing – review & editing: YMN, JAM, JMR, DBM, DMT, ARM, GHB, SR, TYC

## Data and materials availability

All study data, replication markdown, custom functions, and source *xcms* and phenotype objects to replicate all R-based analyses and figures are detailed in the supplementary materials.

## Supplementary Materials for

### Bacterial Strains

Strains used for the feature credentialing pipeline included both *Mycobacterium tuberculosis* H37Rv and the triple auxotrophic *Mycobacterium tuberculosis* H37RvΔ*panCD*Δ*leuCD*Δ*argB* strain mc^2^7902 was described previously (*50*). Reference bacterial strains used for the intermycobacterial lipid comparative analysis include *Mycobacterium tuberculosis* H37Rv, *Mycobacterium tuberculosis* Erdman, *Mycobacterium tuberculosis* HN878, *Mycobacterium bovis* Ravenel ATCC 35720; *Mycobacterium orygis* 51145; *Mycobacterium canetti* NR-49248/NLA00017120, BEI Resources; *Mycobacterium lacus* sp.Nov NRCM 00-25, BEI Resources; *Mycobacterium decipiens* ATCC TSD-117; *Mycobacterium kansasii* ATCC 12478; *Mycobacterium marinum* ATCC BAA-535, *Mycobacterioides abscessus* sp. *abscessus* Clinical Isolate Taiwan-35; *Mycolicibacterium smegmatis* mc^2^155; and *Mycolicibacterium phlei* ATCC 11758. *Staphylococcus aureus* SA113 and the Δ*mprF* mutant were previously described (*46*).

### Bacterial culture

For lipid extraction related to feature credentialling, Mtb H37Rv was cultured in Difco 7H9 media (BD) supplemented with 10% ADN (albumin 0.5%, dextrose 0.2%, NaCl 0.085% [% w/v], final concentration), 0.2% glycerol [% w/v], at 100 rpm agitation and 37°C in 30 mL PETG Media inkwells (Nalgene). Auxotrophic Mtb H37Rv mc^2^7902 was grown in 7H9 + 10% ADN + glycerol as above with supplemented PLAM (L-pantothenate 24 mg/L, L-leucine 5 mg/L, L-arginine 200 mg/L and L-methionine 50 mg/L).

For intermycobacterial comparisons, strains were cultured in batches in parallel in either Biosafety Level 3 or 2 laboratories using identical media conditions as above. For *M. marinum-*specific experiments, *M. marinum* was cultured as above but at 120 rpm and 32°C. Cultures were grown in media containing Tween-80 (0.05%, [v/v]) to mid-log phase (OD 0.6-0.8), then subcultured in detergent-free media. Biological replicates of 10 mL cultures were inoculated therefrom in parallel and grown to turbidity in the absence of detergent to a final pelleted cell volume of between 50 - 100 μL/10mL.

To generate CRISPR interference mutants Mtb H37Rv was grown at 37°C in Difco Middlebrook 7H9 broth or on 7H10 agar supplemented with 0.2% glycerol (7H9) or 0.5% glycerol (7H10), 0.05% Tween-80, and 1x oleic acid-albumin-dextrose-catalase (OADC) and 20 μg/mL kanamycin. For lipidomics, Mtb CRISPR interference gene knockdown with anhydrous tetracycline (ATc) 100 ng/mL every 3 days was performed in parallel with uninduced cultures.

For time course lipidomics 7H9 media supplemented with 10% AN was prepared as above without 0.2% dextrose, or Complete Sauton’s media (*51*) was prepared as described previously with supplemented 0.02% [w/v] dextrose or 0.02% [w/v] lysine as indicated. For growth phase experiments OD600 was measured in a matched culture containing 0.05% [v/v] Tween-80. Tween free cultures of 50 mL cultures in 125 mL PETG Media inkwells (Nalgene) were extracted in 10 mL at each time point.

*S. aureus* SA113 and SA113Δ*mprF* were cultured in Tryptic Soy Broth as described previously (*46*). For time course lipidomics *S. aureus* SA113 was cultured in biological triplicate in 5mL Luria Bertani (LB; Bacto Tryptone 10g/L, Gibco BD; Bacto Yeast Extract 5 g/L, Gibco BD; NaCl 10 g/L, Fisher) media in 14 mL vented snap-cap tubes (Corning) at 200 rpm and 37°C.

### Lipid extraction

All organic solvents were HPLC-grade or Optima® (Fisher). Mycobacterial cultures of 10 mL were pelleted at 4000 rpm (Thermo Sorvall Legend RT centrifuge; Rotor, Sorvall Heraeus 75006445), washed and resuspended in phosphate buffered saline (PBS; pH 7.4, Gibco) twice and resuspended in 2:1 [v/v] methanol:chloroform in 15 mL conical borosilicate glass tubes (Kimble) with phenolic caps (Kimble) for inactivation. After removal from the BSL-3 or BSL-2 by approved inactivation protocols, bacteria were agitated for 1 h at 40 rpm on an orbital shaker. The cells were pelleted at 3000 rpm and the supernatant transferred to clean round-bottomed 15 mL glass tubes (Pyrex). A second extraction with 1:1 [v/v] methanol:chloroform was performed and the resulting supernatant after agitation and centrifugation as above was pooled. Lipids were dried under N_2_-gas (N-EVAP 111, Organomotion) or by GeneVac (Low BP Mixture setting, 30°C; EZ-2, SP Scientific), resuspended in 1:1 [v/v] methanol:chloroform and sonicated for 3 min at 60 sonics/min (Branson 5510 Sonicator). The undissolved particles were pelleted at 3000 rpm. The supernatant was transferred to pre-weighed 4 mL amber vials (Supelco) with teflon-lined caps (Supelco), dried under N_2_-gas and weighed (Mettler Toledo, XP205). Lipid yield ranged between 1-2 mg/10 mL culture. Lipid extracts were resuspended in 1:1 [v/v] methanol:chloroform, normalized to 1 mg/mL and stored at −20°C.

For lipid extraction using alternate solvents deuterated methanol (d4-methanol) and methylene chloride we performed two parallel extracts of Mtb H37Rv auxotroph mc^2^7902 as above, replacing methanol with d4-methanol [v/v], and chloroform with methylene chloride [v/v], respectively.

For lipid extraction from *S. aureus*, cells were washed twice with 20 mM acetate buffer, pH 4.5 as described previously (*46*), thereafter inactivation with 2:1 [v/v] methanol:chloroform, and subsequent lipid extraction proceeded as above for mycobacterial lipids.

### Liquid Chromatography and Mass Spectrometry

An optimized method for mass spectrometry of mycobacterial lipids was described previously (*3*). Briefly, an Agilent 6530 Accurate-Mass QTOF coupled to a 1220 Infinity series HPLC system was used for normal phase chromatography with an Inertsil diol column (3 μm, 2.1 x 150 mm) and a 3 μm cartridge guard column (3 x 10 mm x 2). Lipids were resuspended at 1 mg/mL in solvent B (hexanes:isopropanol, 70:30 [v/v], 0.02% [m/v] formic acid, 0.01% [m/v] ammonium hydroxide) in 2 mL amber screw vials (Agilent) with 250 μL pulled point conical glass insert (Agilent) and PTFE/RS blue screw caps (9 mm, Agilent). 20 μg of total lipid per sample were injected and eluted by binary gradient of 0.15 mL/min from 0% to 100% solvent A (isopropanol:methanol, 70:30 [v/v], 0.02% [m/v] formic acid, 0.01% [m/v] ammonium hydroxide): 0–10 min, 0% A; 17–22 min, 50% A; 30–35 min, 100% A; 40–44 min, 0% A, followed by additional 5 min 0% A postrun.

For reversed phase chromatography we used an InfinityLab Poroshell 120 EC-C18 (3.0 x 50 mm, 1.9 μm, Agilent) with a 3 mm guard column (2.7 μm, Agilent). Lipids were resuspended at 1 mg/mL in solvent A (9:1 methanol:water [v/v], and 2 or 10 mM NH_4_HCOO), and eluted by binary gradient of 0.15 mL/mL from 0% to 100% solvent B (9:1 1-propanol:cyclohexane [v/v], 3 mM NH_4_HCOO): 0-10 min, 0% B; 10-15 min, 100% B; 15-23 min, 100% A. After chromatographic separation, samples were ionized by electrospray with a gas temperature of 325°C, 5 L/min drying gas, 15 psi nebulizer pressure, and 5500 V with time of flight detection within a range of 200 to 3200 *m/z*. For the *lysX* and Rv1619 CRISPRi knockdown lipidomics we used an InfinityLab Poroshell HPH-C18 column (2.1 x 50 mm, 2.7 μm, Agilent) and a 4.6 x 5 mm guard column (2.7 μm, Agilent) with 0.1% [v/v] ammonium hydroxide added to both reversed phase mobile phases.

Collision-induced MS was performed with an optimized voltage for each lipid family ranging from 15-175 V in positive and/or negative modes as indicated in, fig. S3. A narrow window for parent ion isolation (1.3 *m/z*) was used with parent ion identification by chromatographic retention time and *m/z* within 10 ppm of the initial detected *m/z*.

### Processing of MS data

Chromatograms and mass spectra were generated from raw data (.d) using Mass Hunter Qualitative Analysis software (Agilent) and images were formatted for publication in Adobe Illustrator. Centroided data as .mzXML files were obtained from raw data using *msconvert* (*52*) Peak picking was performed using *xcms* (*53*). Standard parameters included a minimum filter of 200 *m/z,* signal-to-noise threshold of 10, peakwidth of 20-120, and *m/z* filter of 200 to 3200 *m/z*. Arguments such as bandwidth (5–30) and ppm (5-10) were optimized to the experiment and chromatography phase. See Data S1 R-Markdown script for a detailed breakdown of *xcms* parameters used for each experiment, including the detailed sequential credentialing pipeline for unknown lipids, the methylene transform analysis and plots, as well as *ggplot2* scripts used to generate each R-based main and supplemental panel. Included is an additional R-markdown of the custom functions used to access *xcms* and CAMERA that were not included in the *limms* package (*16*). Statistical analysis not using *limms* used Graphpad Prism (v10.3).

Briefly, we used *limms* for differential abundance analysis and imputing local intensity minima to enable pairwise or compound statistical contrasts, as described previously (*16*). Thereafter, linear models were tested to generate Benjamini-Hochberg adjusted *P*-values reported for the contrast. For the credentialing pipeline, contaminants and idiosyncratic signals were identified by statistical comparison to solvent-only HPLC-MS control runs. Ions that did not meet a significance threshold of *P* < 0.05, fold change > 2 were censored. The core lipid structures in MycoMassDB were propagated to ∼200,000 theoretical variants using LOBSTAHs (Data S2) (*21*). The MycoLOBSTAHs database worked best as an indicator that an observed masses contained C, H, O, N, P, or S atoms in a ratio typical of a lipid rather than an annotation tool, and was used to identify inorganic ions and salt clusters due to their low CH content. Automated mass matching was performed using the dbMatch function within *limms* and MycoMassDB or MycoLOBSTAHs databases to 10 ppm and no retention time matching. Adducts of H^+^, NH_4_ ^+^, Na^+^ or ^13^C isotopes were identified using the R-package CAMERA using default parameters in the positive mode polarity (*22*). Identifying *in source* lipid aggregation required manual recognition of multimers of diacyl trehalose (DAT), phosphatidylinositol (PI), phosphatidylethanolamine (PE), phosphatidylinositol mannosides (PIMs) and glycerolipid fragments, as seen in, Fig. S1. To generate the methylene transformed co-efficient, or the CH_2_ Kendrick mass defect, the detected *m/z* of each lipid was multiplied by 14/14.01565 and subtracted by the floor integer of the transformed mass to give the transformed mass defect (Equation 1) (*25*). Lipids were grouped based on their retention time (± 3 min) and transformed mass defect coefficient (±0.025).

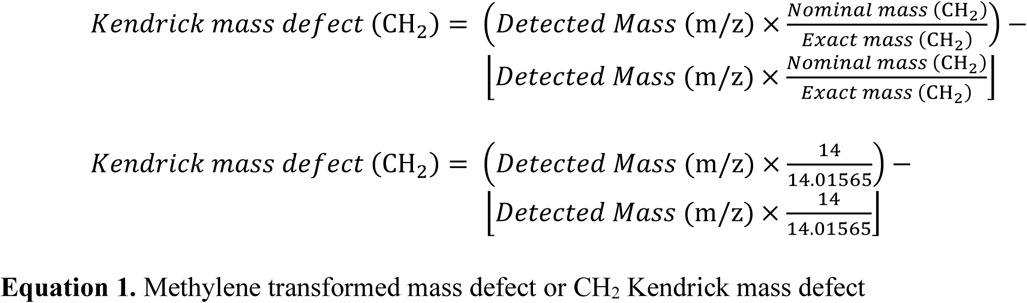

### Domain architecture of lysyltransferases in Mtb

Using the Uniprot entries for Mtb *lysX*/Rv1640c (P9WFUZ), *S. aureus mprF* (Q2G2M2) and *C. glutamicum alaDAGS*/cg1103 (P35867) genes with shared Interpro classified domains were identified in Mtb (*49*). The data on gene and domain length were obtained from Interpro. Visualization used the R-package *gggenes* function *geom_gene_arrows*.

### Generation of Rv1619 and *lysX* CRISPRi knockdown strains

Individual CRISPRi plasmids were generated following the protocol outlined in (*54*) using Addgene plasmid #163631. Briefly, the CRISPRi plasmid backbone was digested with BsmBI-v2 (NEB #R0739L) and subsequently gel purified. sgRNAs were designed to target the non-template strand of the target gene ORF. For each sgRNA, two complementary oligonucleotides with compatible sticky-end overhangs were annealed and ligated into the BsmBI-digested plasmid backbone using T4 DNA ligase (NEB #M0202M). For lysX/Rv1640c the top oligonucleotide was 5’-GGGAGCCCAGAACTCCCGATAGCCCAGCA-3’ and bottom oligonucleotide was 5’-AAACTGCTGGGCTATCGGGAGTTCTGGGC-3’, predicted strength 0.91 (PAM 5’-CCAGAAC-3’). For Rv1619 the top oligonucelotide was 5’-GGGAGCCGCGAAGTCGGCGACCAGCTG-3’ and the bottom oligonucleotide was 5’-AAACCAGCTGGTCGCCGACTTCGCGGC-3’, predicted strength 0.96 (PAM 5’-GGGGAAT-3’).

Ligation products were transformed into NEB 5-alpha chemically competent cells and selected on LB + kanamycin (50 μg/mL). Individual colonies were picked and the correct sgRNA sequence was confirmed by Sanger sequencing before subsequent transformation into *M. tuberculosis* H37Rv.

For transformations of *M. tuberculosis* H37Rv, cultures of at least 20 mL were grown to late logarithmic phase (OD600 of 0.8–1.0) and cells were pelleted at 4,000 × g (room temperature) for 10 min. The cell pellet was washed three times in sterile 10% glycerol. The washed cells were then resuspended in 10% glycerol in a final volume of 5% of the original culture volume. For each transformation, 100 ng plasmid DNA and 100 μL of electrocompetent mycobacteria were mixed and transferred to a 2 mm electroporation cuvette (Bio-Rad #1652082). Electroporation was performed using the Gene Pulser X cell electroporation system (Bio-Rad #1652660) set at 2500 V, 700 Ω, and 25 μF. Bacteria were recovered in 7H9 for 24 hr. After the recovery incubation, cells were plated on 7H10 with 20 μg/mL kanamycin. Plates were subsequently incubated at 37°C for 14-21 days. Individual colonies were picked and the correct sgRNA sequence was confirmed by Sanger sequencing.

### Synthesis of sn3-lysyldiacylglycerol

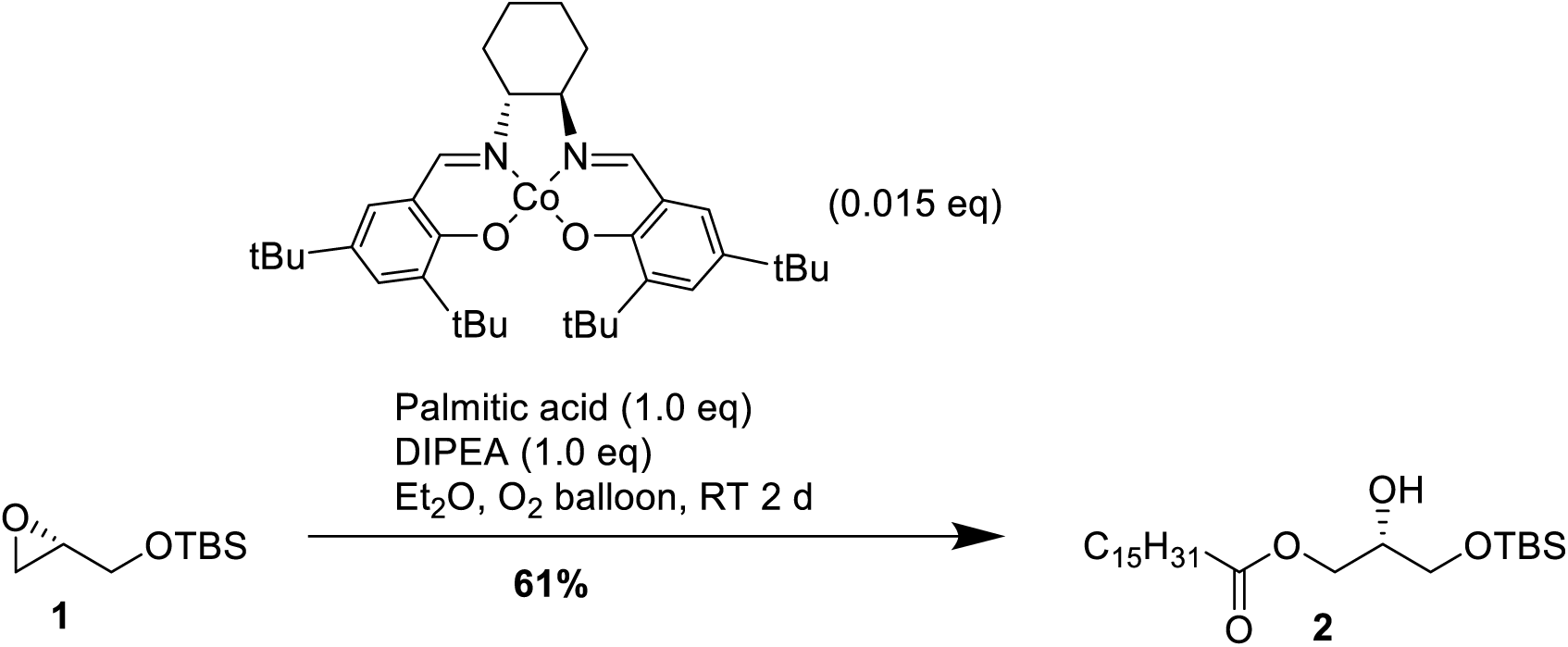

#### (*S*)-3-((tert-Butyldimethylsilyl)oxy)-2-hydroxypropyl palmitate **2**

An oven-dried flask was charged with palmitic acid (6.9 g, 26.9 mmol, 1.0 eq), (*R,R*)-Co[salen] (240 mg, 0.398 mmol, 1.5 mol%) and Et_2_O (30 mL). The flask was equipped with an oxygen balloon that provided a flow of O_2_ directly into the resulting solution. The resulting mixture was stirred for 30 min, during which the reaction turned dark brown. Afterwards, DIPEA (4.6 mL, 26.5 mmol, 1.0 eq) was added, and after 5 min of stirring, **1** (5.6 mL, 26.5 mmol). The resulting reaction was stirred at RT, until after 2 d, TLC (95 : 5 pentane : Et_2_O) indicated near-complete consumption of **1**. The reaction was subsequently concentrated, dispersed onto celite and purified using flash column chromatography on silica (pentane : Et_2_O 95 : 5 to 90 : 10) to obtain **2** as a white wax (7.2 g, 16.3 mmol, 61%).

^1^H NMR (400 MHz, Chloroform-*d*) δ 3.85 (dd, *J* = 11.9, 3.2 Hz, 1H), 3.66 (dd, *J* = 12.0, 4.8 Hz, 1H), 3.09 (tt, *J* = 4.0, 2.9 Hz, 1H), 2.77 (dd, *J* = 5.1, 4.0 Hz, 1H), 2.64 (dd, *J* = 5.2, 2.7 Hz, 1H), 2.34 (t, *J* = 7.5 Hz, 2H), 1.63 (p, *J* = 7.4 Hz, 2H), 1.25 (d, *J* = 2.1 Hz, 24H), 0.90 (s, 9H), 0.89 – 0.83 (m, 2H), 0.08 (d, *J* = 3.6 Hz, 6H). ^13^C NMR (101 MHz, cdcl_3_) δ 179.22, 63.71, 52.44, 44.49, 33.90, 31.92, 29.69, 29.68, 29.66, 29.65, 29.64, 29.59, 29.43, 29.36, 29.24, 29.06, 25.86, 24.68, 22.69, 14.12, −5.32, −5.36.

#### (*S*)-3-((tert-Butyldimethylsilyl)oxy)-2-(palmitoyloxy)propyl palmitate **3**

An oven-dried flask under N_2_-atmosphere was charged with **2** (8.06 g, 18.1 mmol), palmitic acid (5.21 g, 20.3 mmol, 1.1 eq), anhydrous DCM (120 mL) and DMAP (221 mg, 1.81 mmol, 10 mol%). The resulting solution was cooled to 0 °C over 5 min before addition of DCC (4.20 g, 20.3 mmol, 1.1 eq). The reaction was subsequently brought to RT and stirred for 16 h before TLC (98 : 2 pentane : Et_2_O) indicated near-complete conversion of the starting material. The reaction was subsequently concentrated *in vacuo*, dispersed onto celite and purified using flash column chromatography on silica (pentane : Et_2_O 99 : 1 to 90 : 10) to provide **3** as a white wax (10.2 g, 15.0 mmol, 83%).

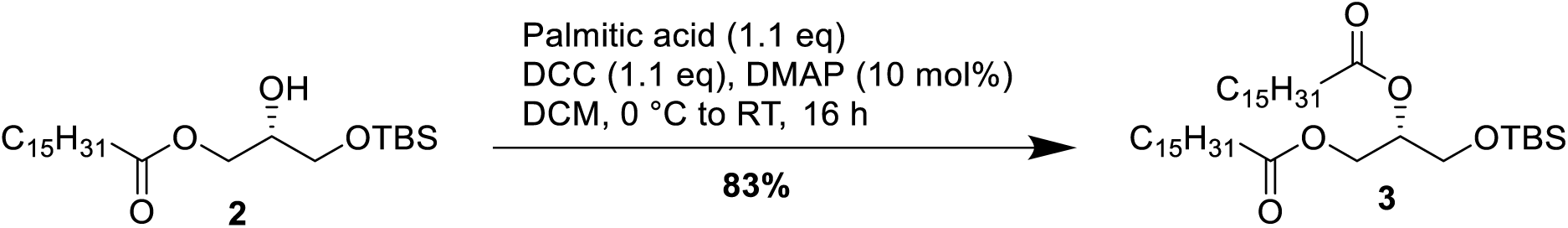

^1^H NMR (400 MHz, cdcl_3_) δ 5.05 (dqt, *J* = 9.2, 5.7, 2.8 Hz, 1H), 4.32 (ddt, *J* = 11.9, 4.4, 2.2 Hz, 1H), 4.13 (ddt, *J* = 13.1, 6.8, 3.4 Hz, 1H), 3.69 (ddd, *J* = 5.3, 3.7, 1.6 Hz, 2H), 2.27 (tdt, *J* = 7.1, 5.1, 2.4 Hz, 4H), 1.59 (dtq, *J* = 10.6, 7.5, 4.8 Hz, 4H), 1.23 (d, *J* = 4.9 Hz, 48H), 0.92 – 0.80 (m, 15H), 0.05 – 0.00 (m, 6H). ^13^C NMR (101 MHz, cdcl_3_) δ 173.36, 71.64, 62.40, 61.43, 34.30, 34.12, 31.90, 29.67, 29.64, 29.60, 29.45, 29.34, 29.26, 29.10, 29.07, 25.71, 25.69, 24.92, 24.89, 22.66, 18.15, 14.06, −5.57. HRMS (ESI+) m/z calcd for [M+Na]^+^ C_41_H_82_O_5_SiNa = 705.5824; found 705.5821

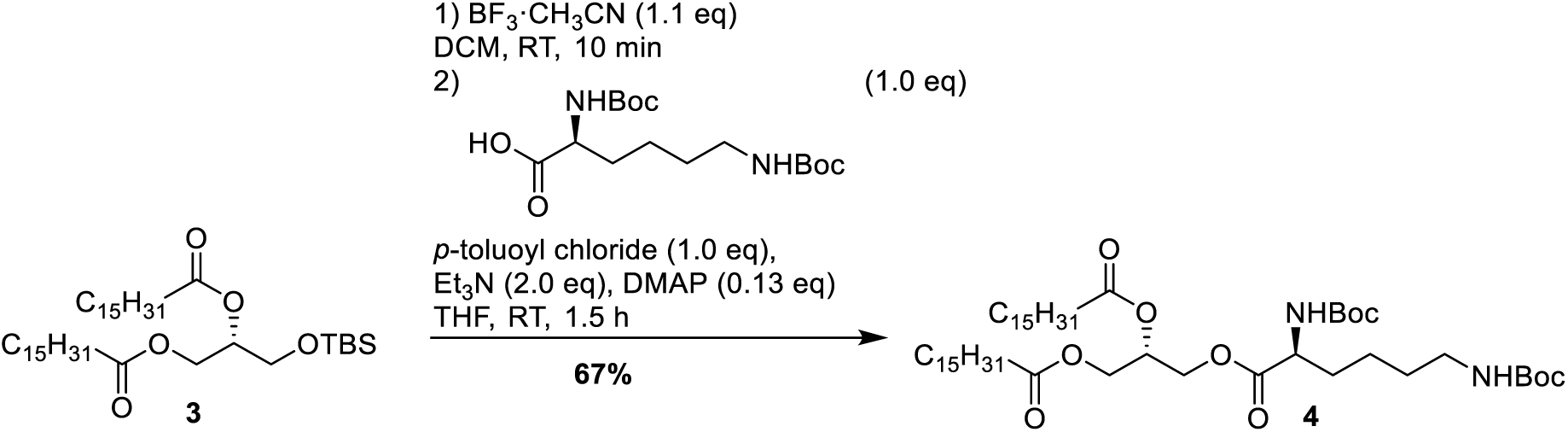

#### (*S*)-3-((*N*^2^,*N*^6^-bis(*tert*-butoxycarbonyl)-*L*-lysyl)oxy)propane-1,2-diyl-dipalmitate **4**

To an oven-dried flask, charged with TBS-protected diacylglyceride (1.04 g, 1.52 mmol) and DCM (15.2 mL) under nitrogen atmosphere, was added a 16 wt% solution of BF_3_·CH_3_CN complex in CH_3_CN (1.14 g, 1.67 mmol, 1.1 eq). The resulting yellow reaction was stirred at RT until completion after 10 min (TLC monitoring, pentane : Et_2_O 98 : 2). The reaction was subsequently quenched by addition of chilled phosphate buffer (20 mL, 1 M, pH = 7) and diluted with Et_2_O (20 mL). The resulting organic phase was washed with brine (20 mL), dried over MgSO_4_ and concentrated *in vacuo* to yield the deprotected alcohol as a white solid (820 mg, 1.44 mmol, 95%).

This intermediate product was immediately redissolved in THF (15.2 mL), followed by addition of *N*^2^,*N*^6^-Bis-boc-*L*-lysine (52.8 mg, 1.52 mmol, 1.0 eq) and *p*-toluoyl chloride (200 μL, 1.51 mmol, 1.0 eq). After complete dissolution of the starting materials, Et_3_N (0.42 mL, 3.02 mmol, 2.0 eq) and DMAP (23 mg, 0.19 mmol, 0.13 eq) were added, and an atmosphere of N_2_ was established. The resulting reaction was stirred at RT until complete consumption of the amino-acid starting material (1.5 h, TLC monitoring, pentane : EtOAc : EtOH 6 : 3 : 1, ninhydrin stain). The reaction was acidified to pH 6 using 1 M aqueous HCl solution and partitioned between water (20 mL) and EtOAc (20 mL). The aqueous phase was extracted with EtOAc (2 x 20 mL). The combined organic phases were washed with sat. aqueous NaHCO_3_ solution (30 mL), water (30 mL) and brine (30 mL), dried over MgSO_4_ and concentrated *in vacuo*. The resulting red oil was dispersed on silica and purified using flash column chromatography on silica (pentane : Et_2_O 80 : 20 to 70 : 30) to obtain **4** as a white wax (910 mg, 1.01 mmol, 67%).

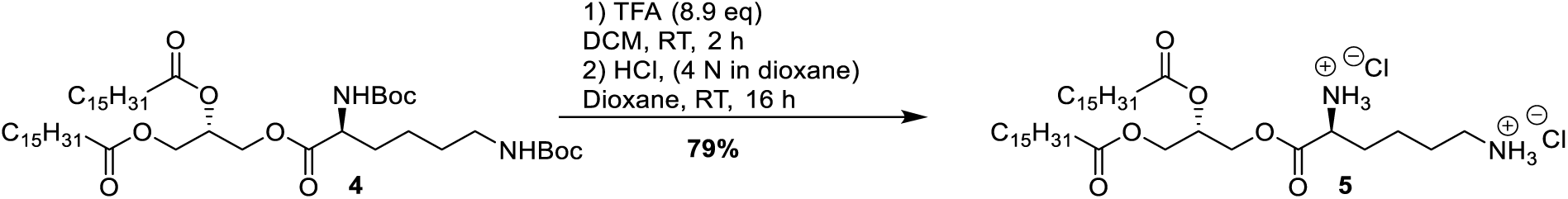

^1^H NMR (400 MHz, CDCl_3_) δ 5.26 (p, *J* = 5.2 Hz, 1H), 5.07 (d, *J* = 8.3 Hz, 1H, (N-H)), 4.64 (s, 1H, (N-H)), 4.37 – 4.19 (m, 4H), 4.13 (dd, *J* = 11.9, 5.9 Hz, 1H), 3.11 (q, *J* = 6.5 Hz, 2H), 2.31 (td, *J* = 7.5, 2.1 Hz, 4H), 1.88 – 1.73 (m, 1H), 1.61 (dt, *J* = 13.2, 6.7 Hz, 7H), 1.55 – 1.46 (m, 1H), 1.46 – 1.40 (m, 18H), 1.40 – 1.34 (m, 2H), 1.25 (s, 48H), 0.92 – 0.84 (m, 6H). ^13^C NMR (101 MHz, cdcl_3_) δ 173.28, 172.89, 172.33, 68.67, 62.99, 61.89, 53.23, 39.96, 34.14, 34.03, 32.17, 31.92, 29.70, 29.65, 29.63, 29.50, 29.48, 29.36, 29.29, 29.27, 29.12, 29.09, 28.42, 28.31. HRMS (ESI+) m/z calcd for [M+Na]^+^ C_51_H_96_N_2_O_10_Na = 919.6957; found 919.6944

#### (*S*)-3-((*L*-lysyl)oxy)propane-1,2-diyl-dipalmitate bis hydrochloride salt **5** (Lys-DAG double chloride salt)

An oven-dried flask was charged with **4** (80 mg, 0.089 mmol), DCM (0.8 mL) and trifluoroacetic acid (0.2 mL, 0.80 mmol, 8.9 eq) under N_2_ atmosphere. The resulting reaction was stirred at RT for 2 h (TLC monitoring, EtOAc : EtOH 4 : 1, ninhydrin stain). The reaction was then concentrated *in vacuo*, before redissolution in 1,4-dioxane (1.5 mL). To the resulting solution was added HCl (4 M solution in 1,4-dioxane, 89 μL). The reaction was stirred for 16 h at which point a white precipitate evidenced that the reaction had progressed. The reaction was concentrated *in vacuo*, redissolved in water and lyophilized to obtain **5** as a fine white powder (49 mg, 0.070 mmol, 79%).

^1^H NMR (600 MHz, CD_3_OD_SPE) δ 5.36 (tt, *J* = 6.4, 3.8 Hz, 1H), 4.48 (dd, *J* = 11.9, 4.2 Hz, 1H), 4.45 – 4.39 (m, 2H), 4.20 (dd, *J* = 12.1, 6.8 Hz, 1H), 4.12 (t, *J* = 6.5 Hz, 1H), 3.02 – 2.96 (m, 2H), 2.35 (dt, *J* = 10.1, 7.4 Hz, 4H), 2.06 – 1.89 (m, 2H), 1.75 (dq, *J* = 9.7, 7.7 Hz, 2H), 1.67 – 1.57 (m, 5H), 1.57 – 1.49 (m, 1H), 1.30 (d, *J* = 5.6 Hz, 48H), 0.91 (t, *J* = 7.0 Hz, 6H).

^13^C NMR (151 MHz, MeOD) δ 174.81, 174.44, 170.17, 70.32, 65.40, 64.14, 63.25, 53.68, 49.57, 40.26, 35.03, 34.88, 33.09, 31.02, 30.83, 30.78, 30.69, 30.66, 30.50, 30.47, 30.23, 30.20, 28.07, 26.02, 23.75, 23.23, 14.45. HRMS (ESI+) m/z calcd for [M+H]^+^ C_41_H_80_N_2_O_6_H = 697.6089; found 697.6084

### Spectra

#### ^1^H-NMR of 2

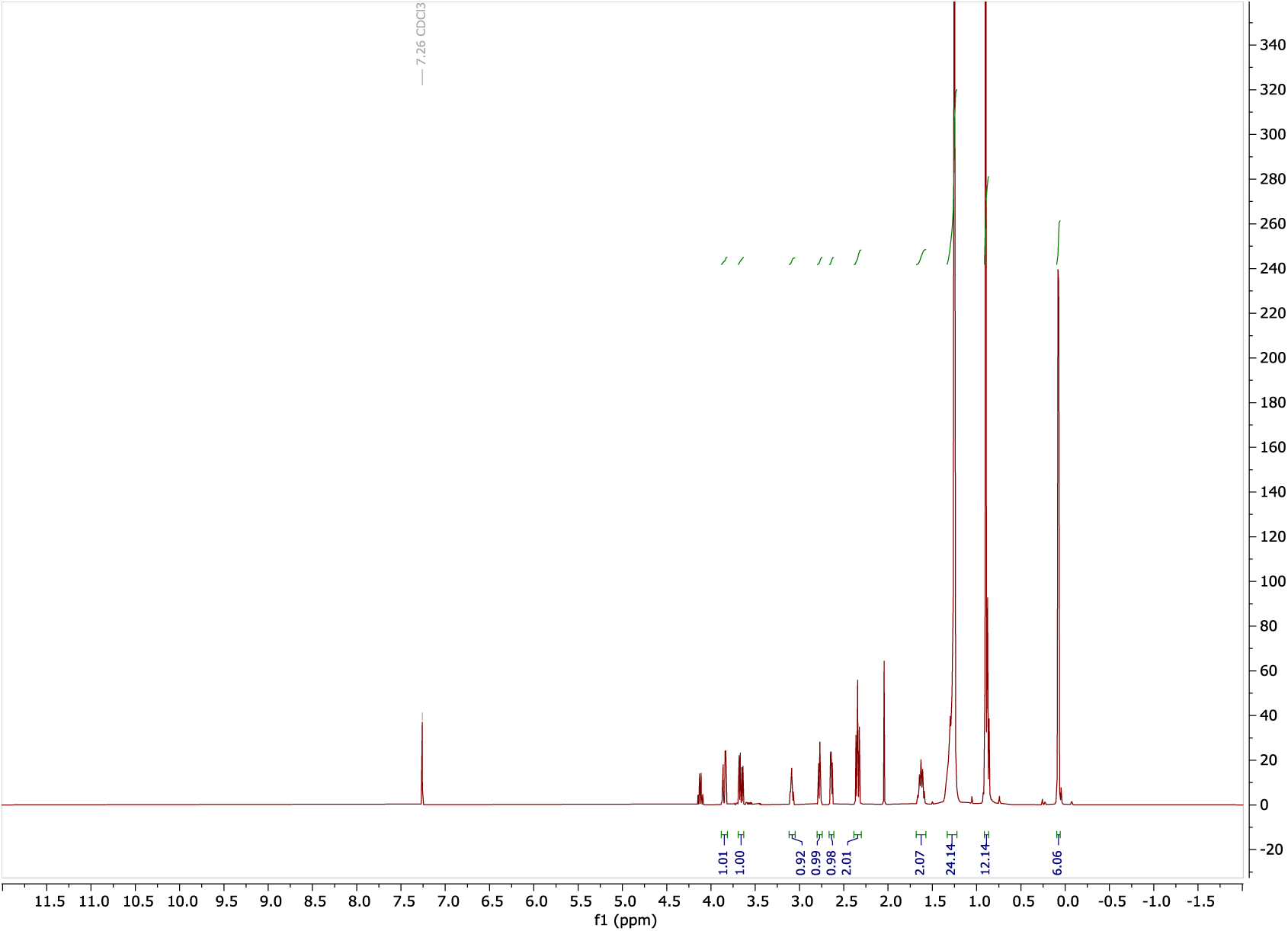

#### ^13^C-NMR of 2

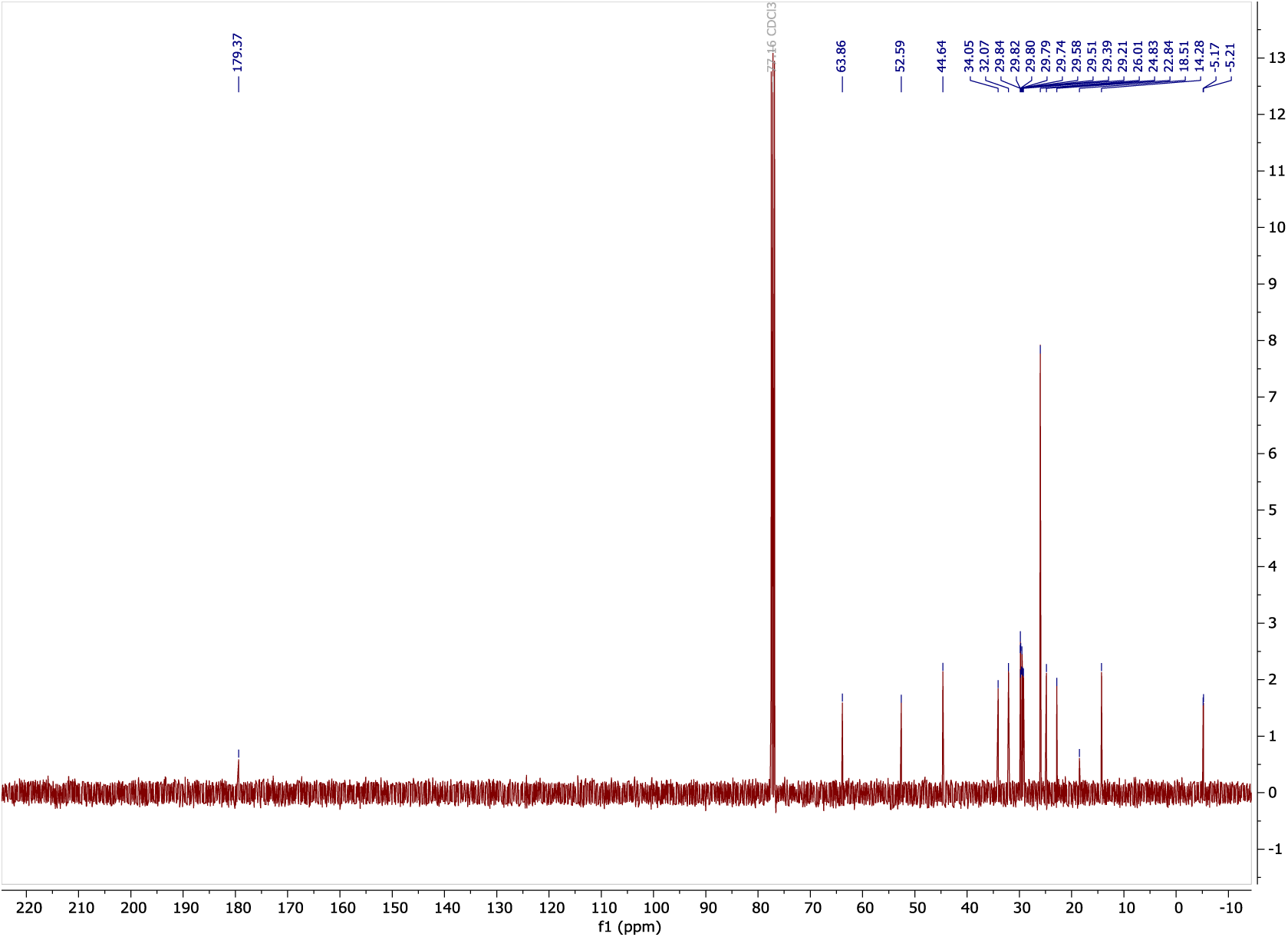

#### COSY of 2

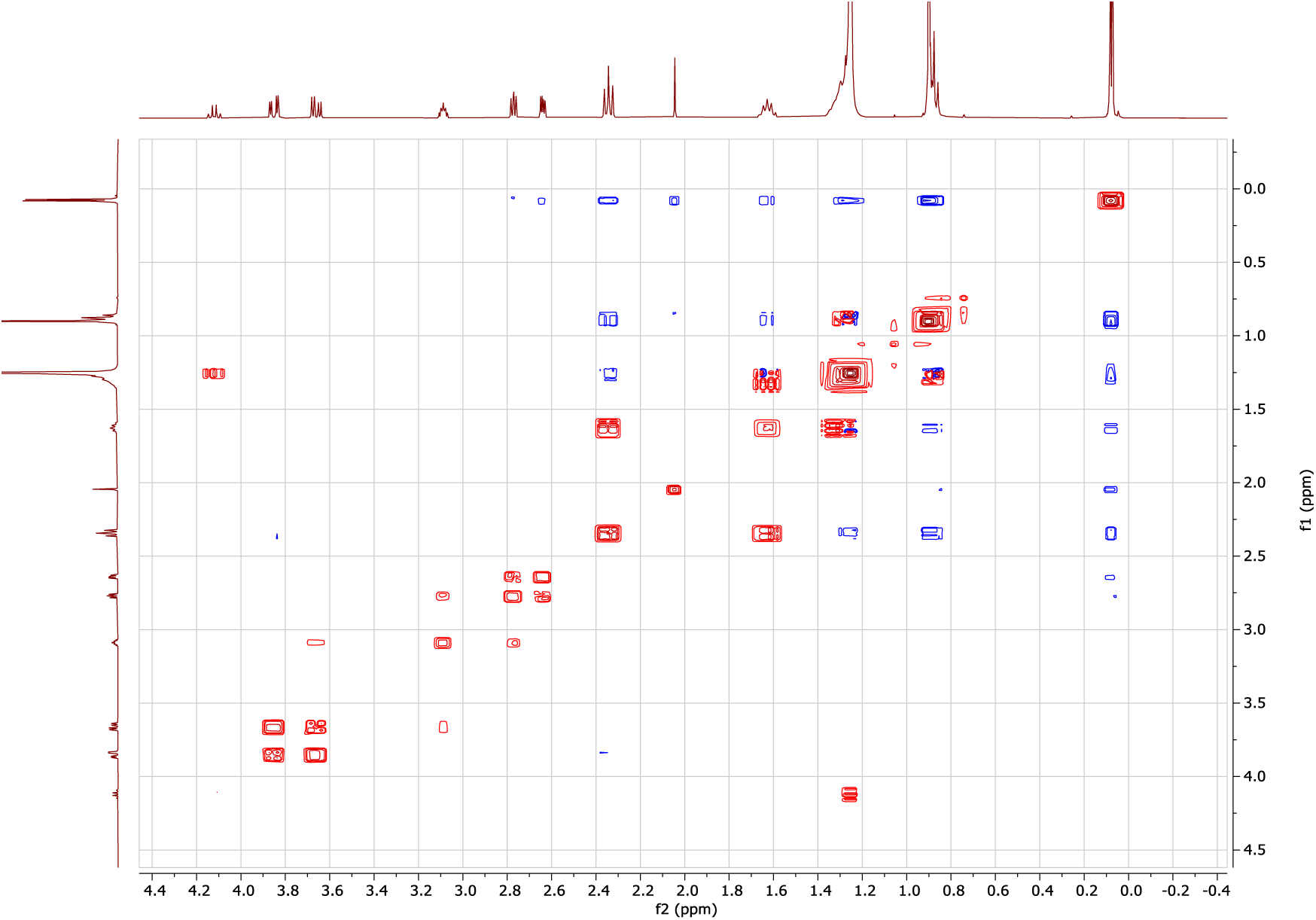

#### ^1^H-NMR of 3

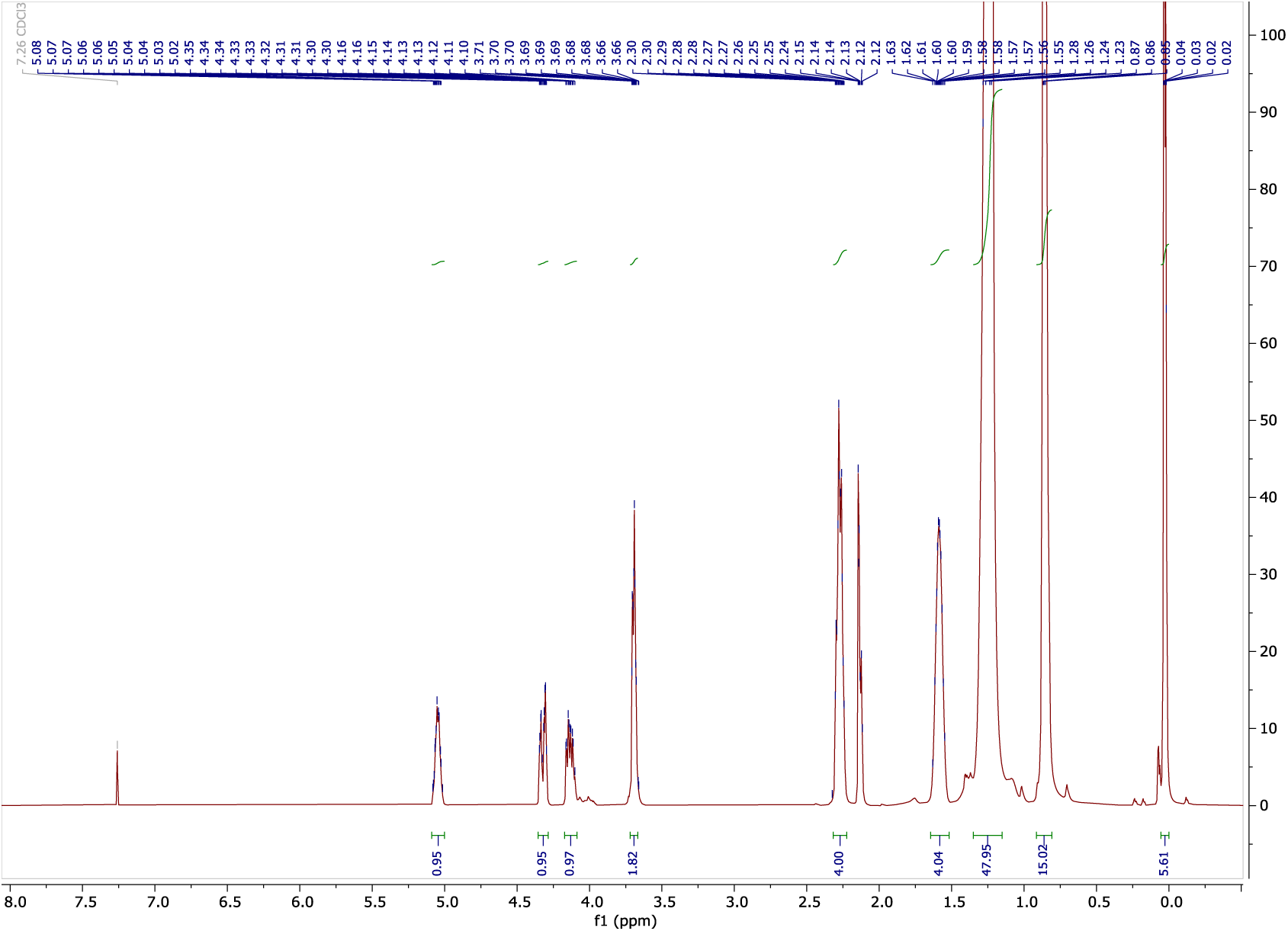

#### ^13^C-NMR of 3

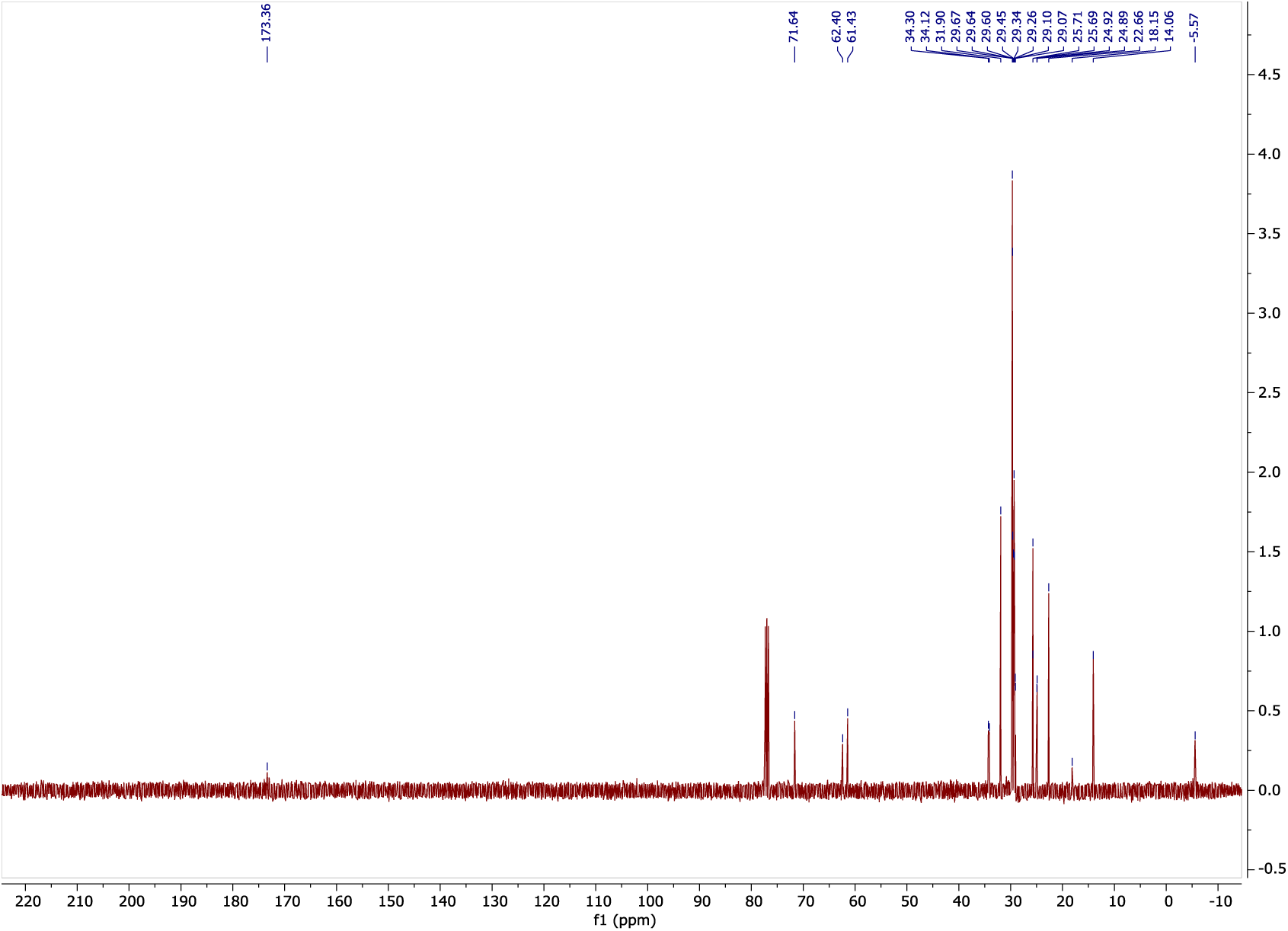

#### COSY of 3

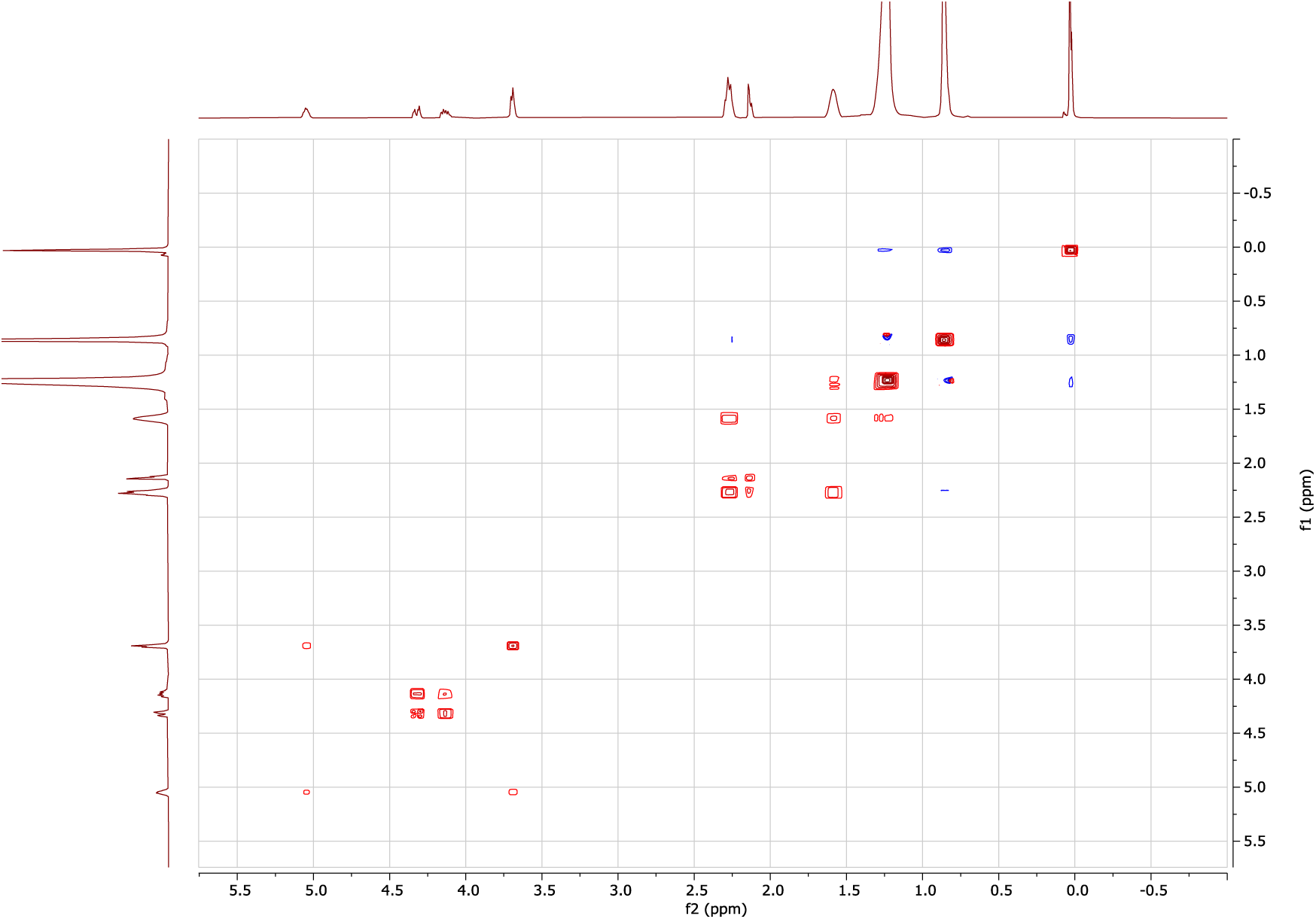

#### HSQC of 3

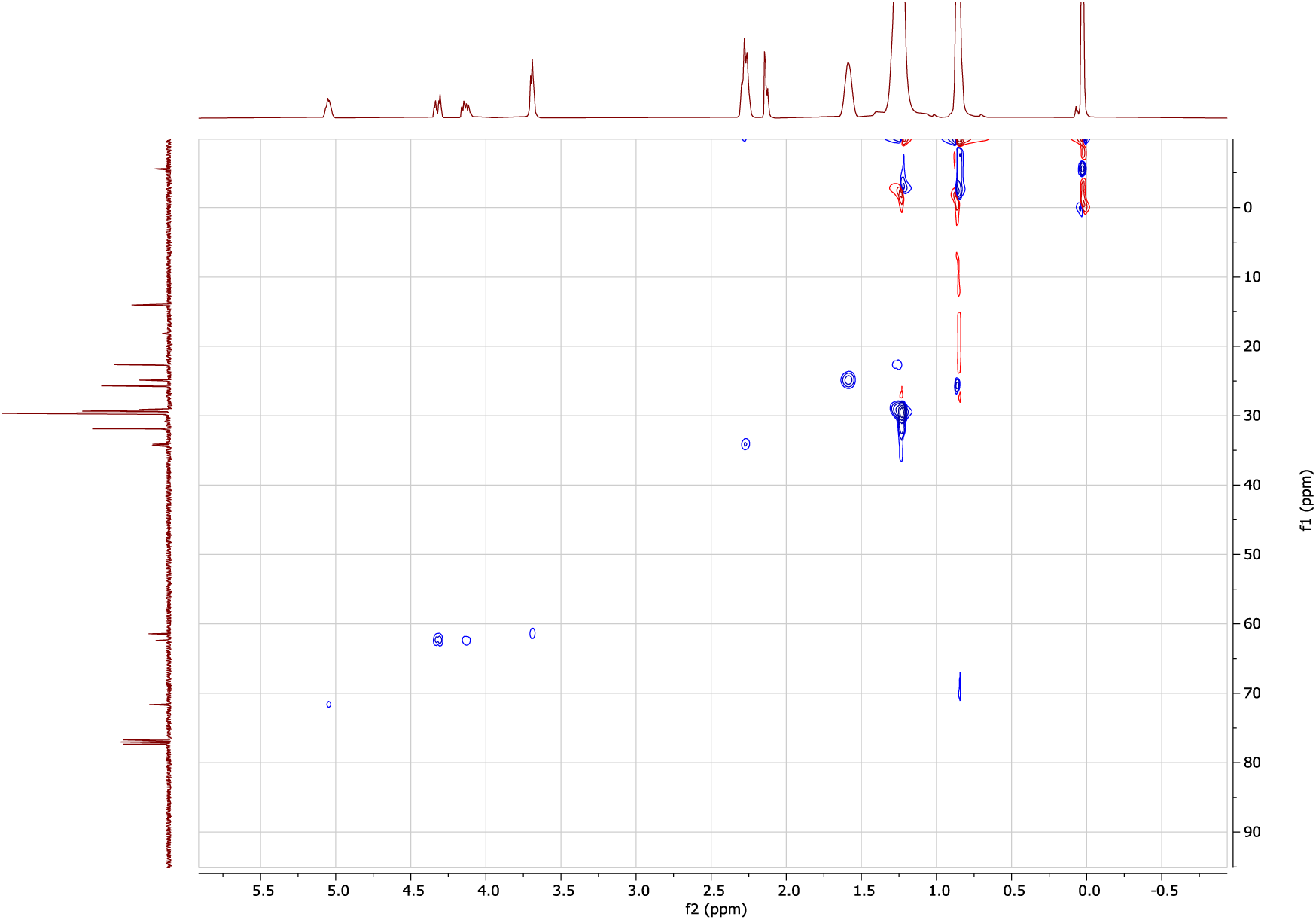

#### HMBC of 3

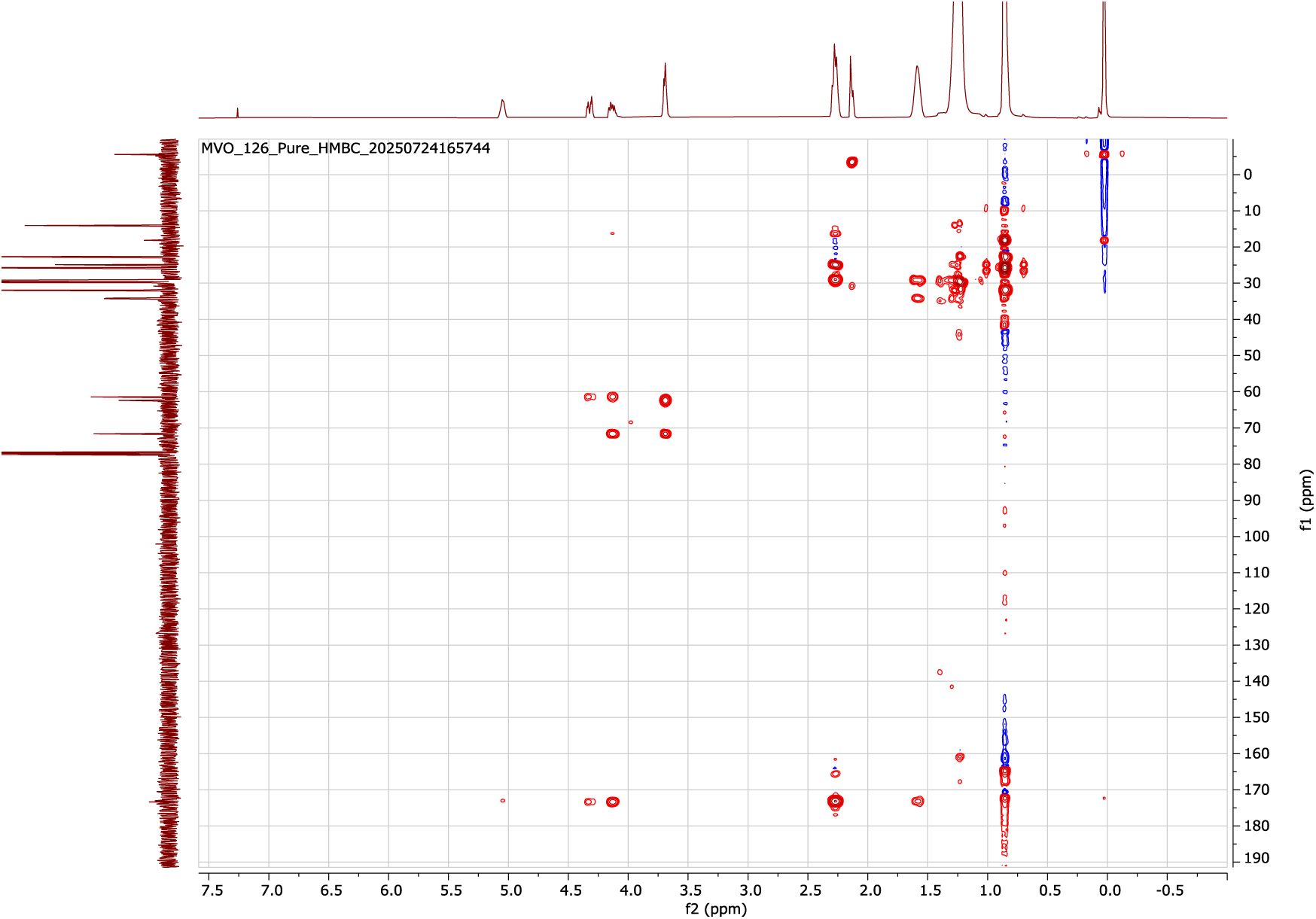

#### ESI-HRMS of 3

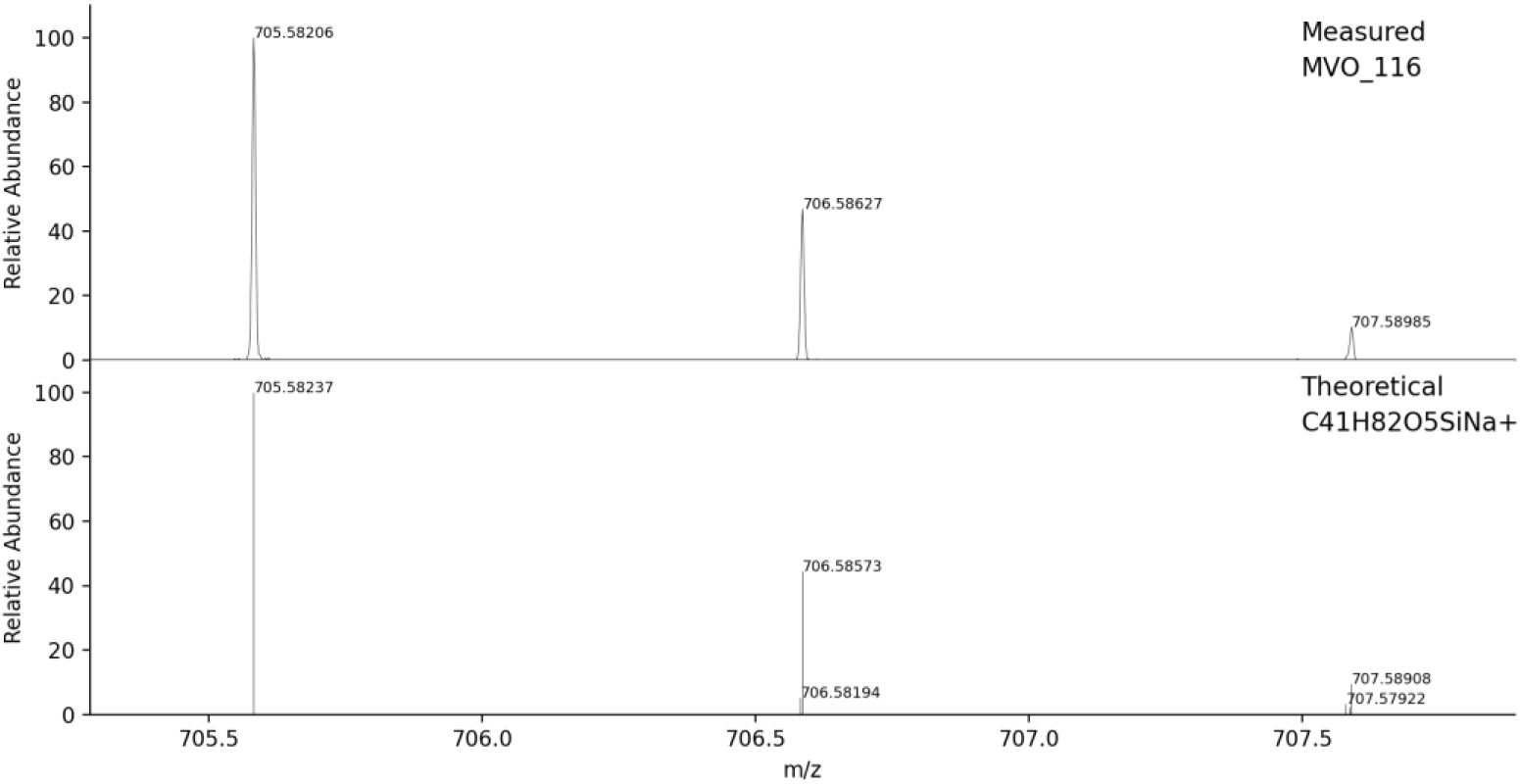

#### ^1^H-NMR of 4

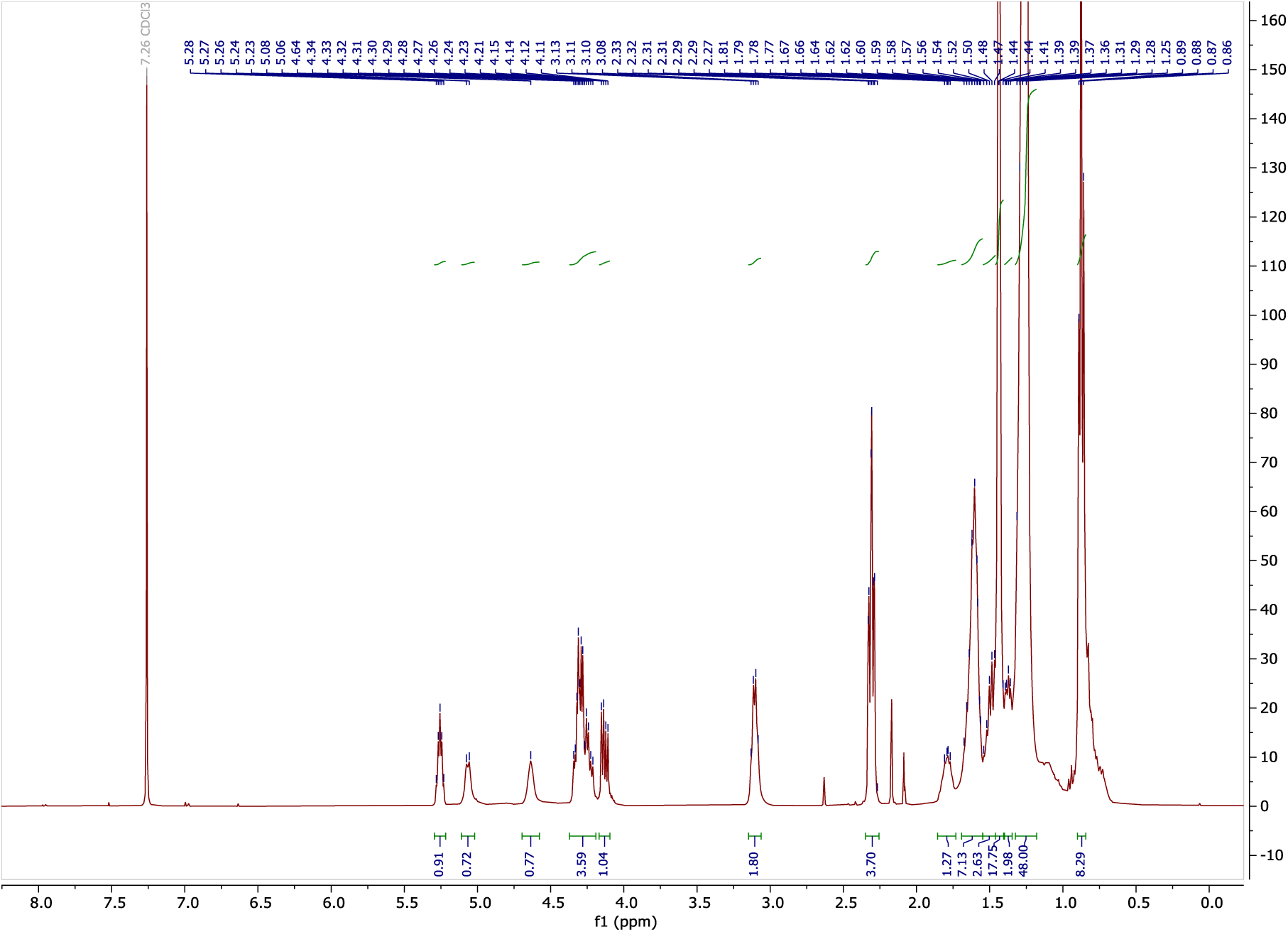

#### ^13^C-NMR of 4

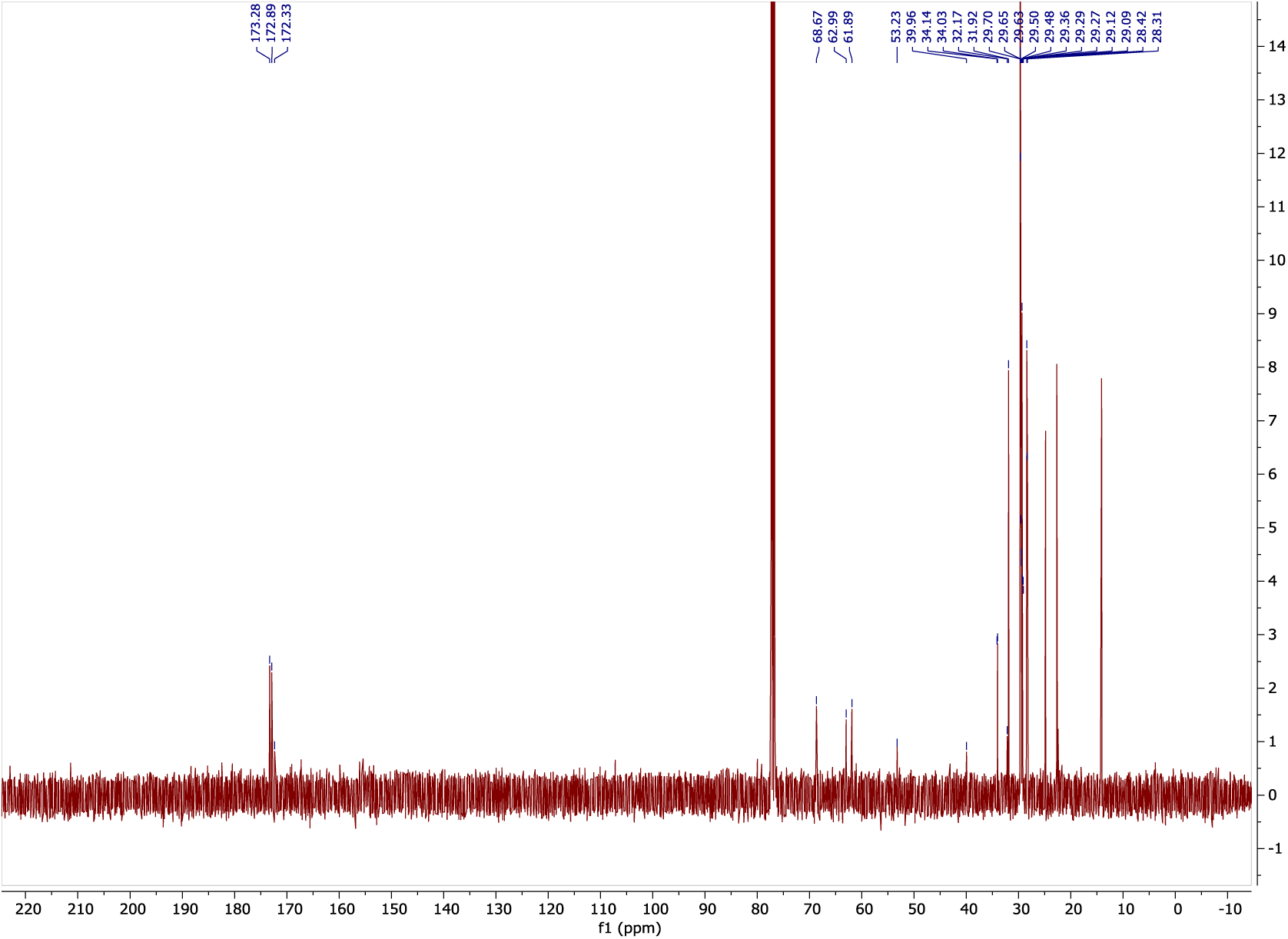

#### COSY of 4

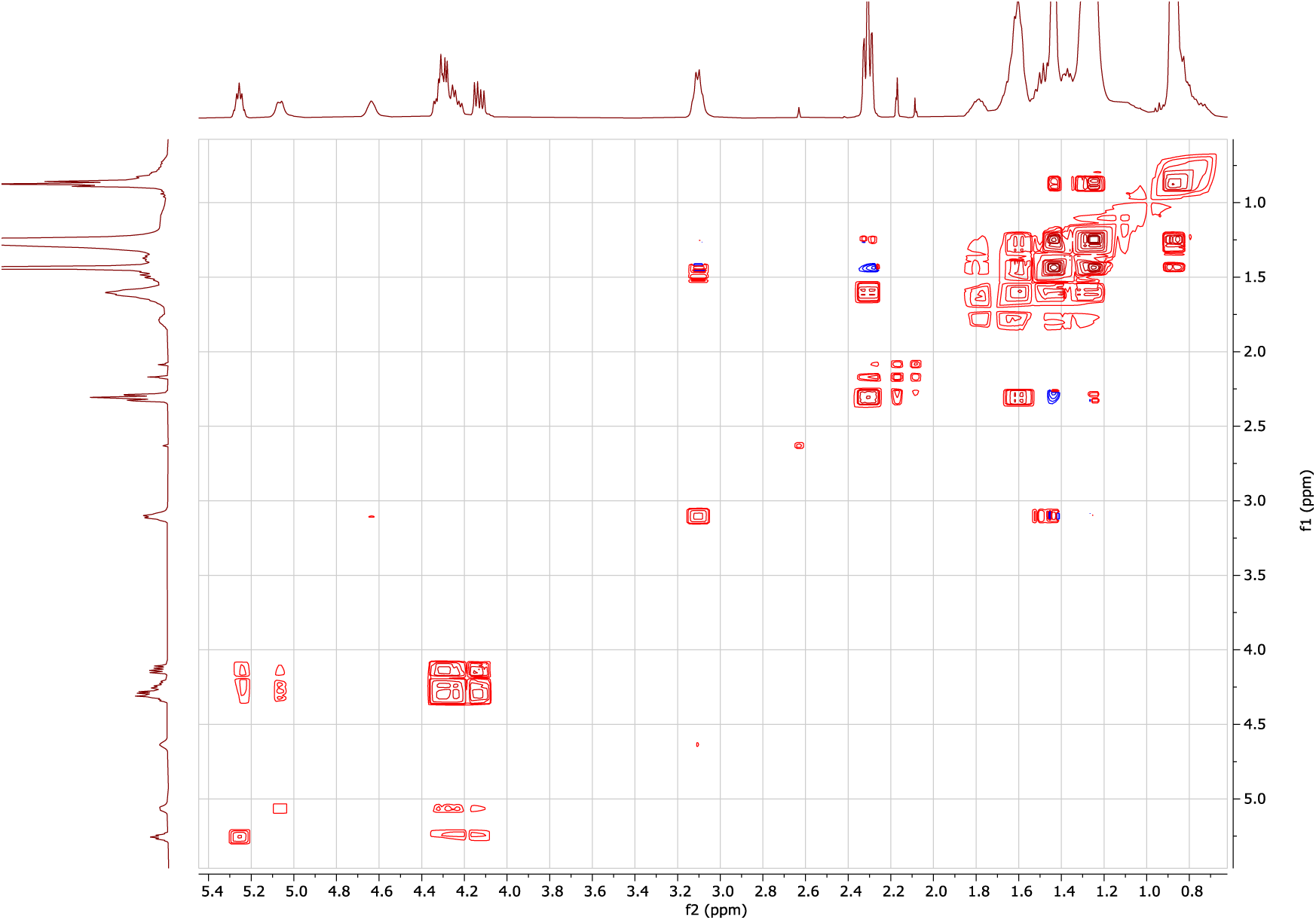

#### gHSQC of 4

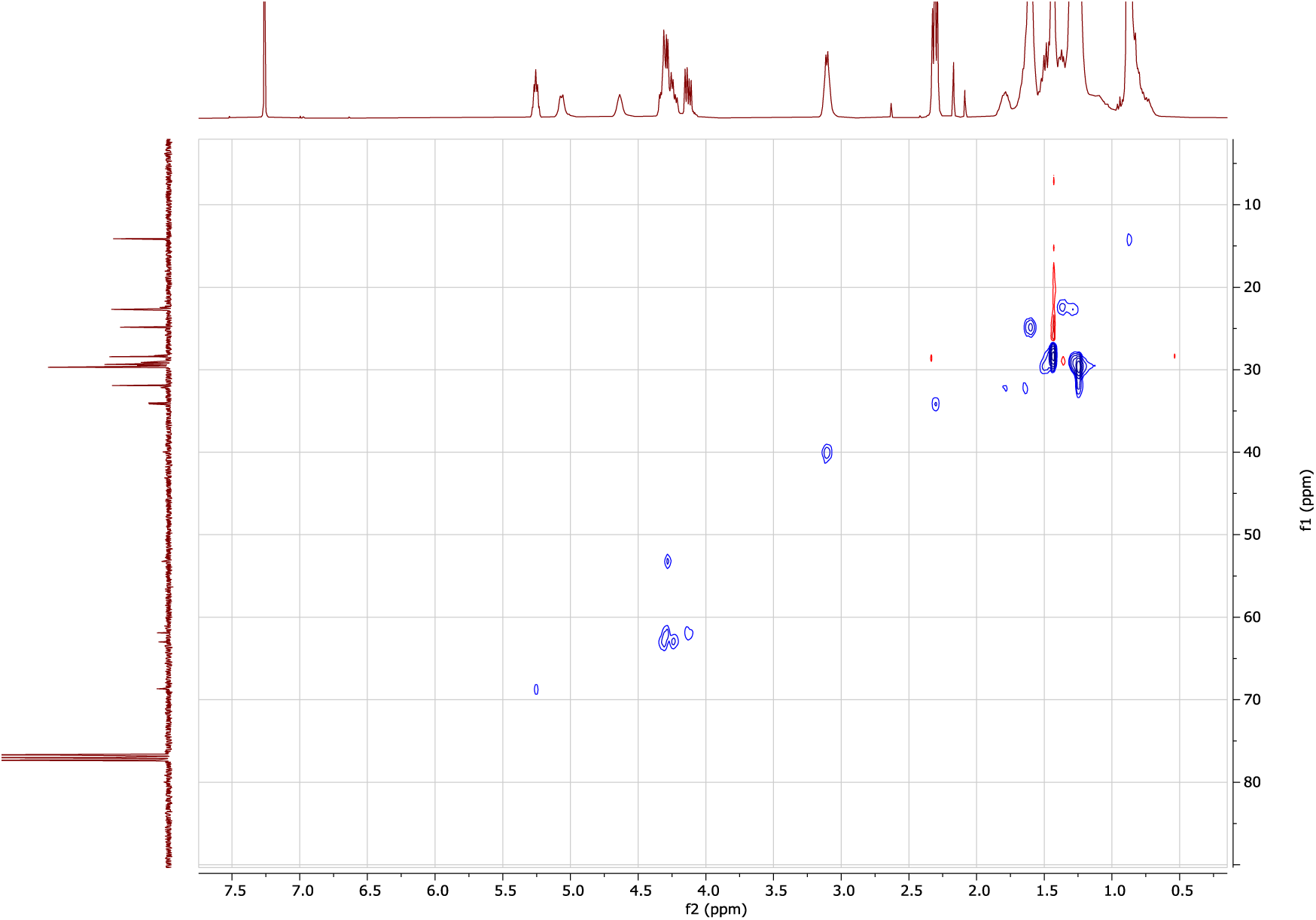

#### gHMBC of 4

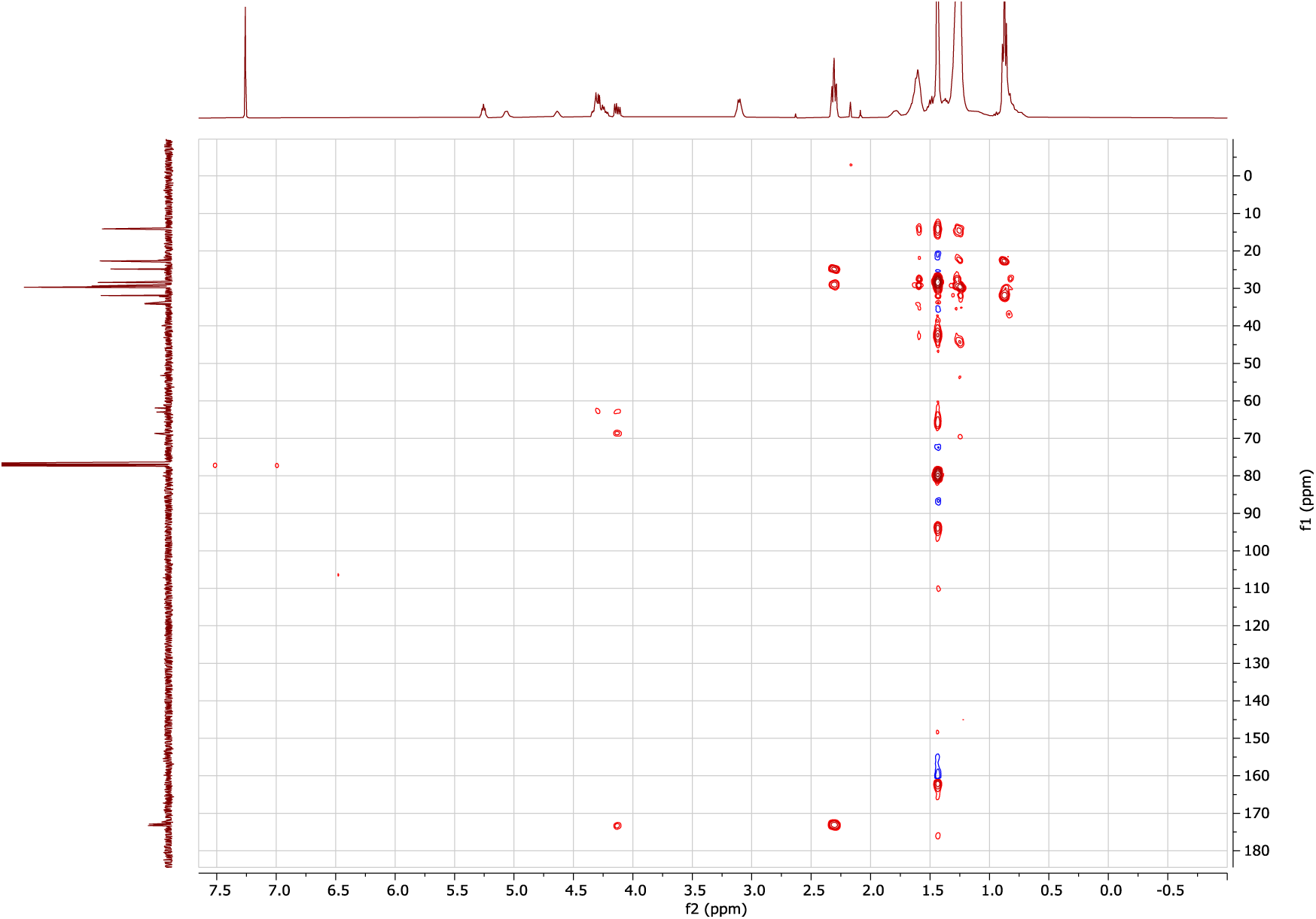

#### ESI-HRMS of 4

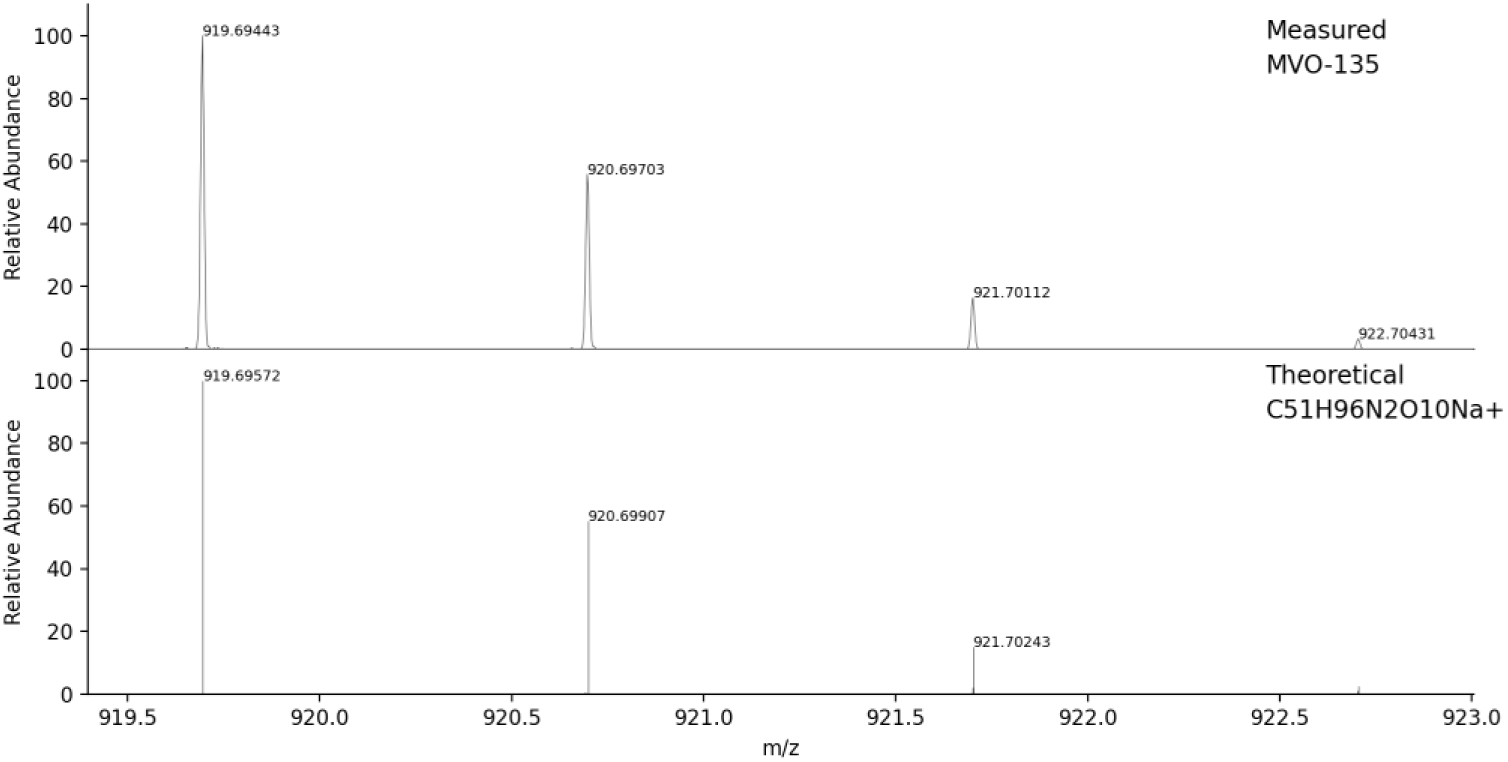

#### ^1^H-NMR of 5

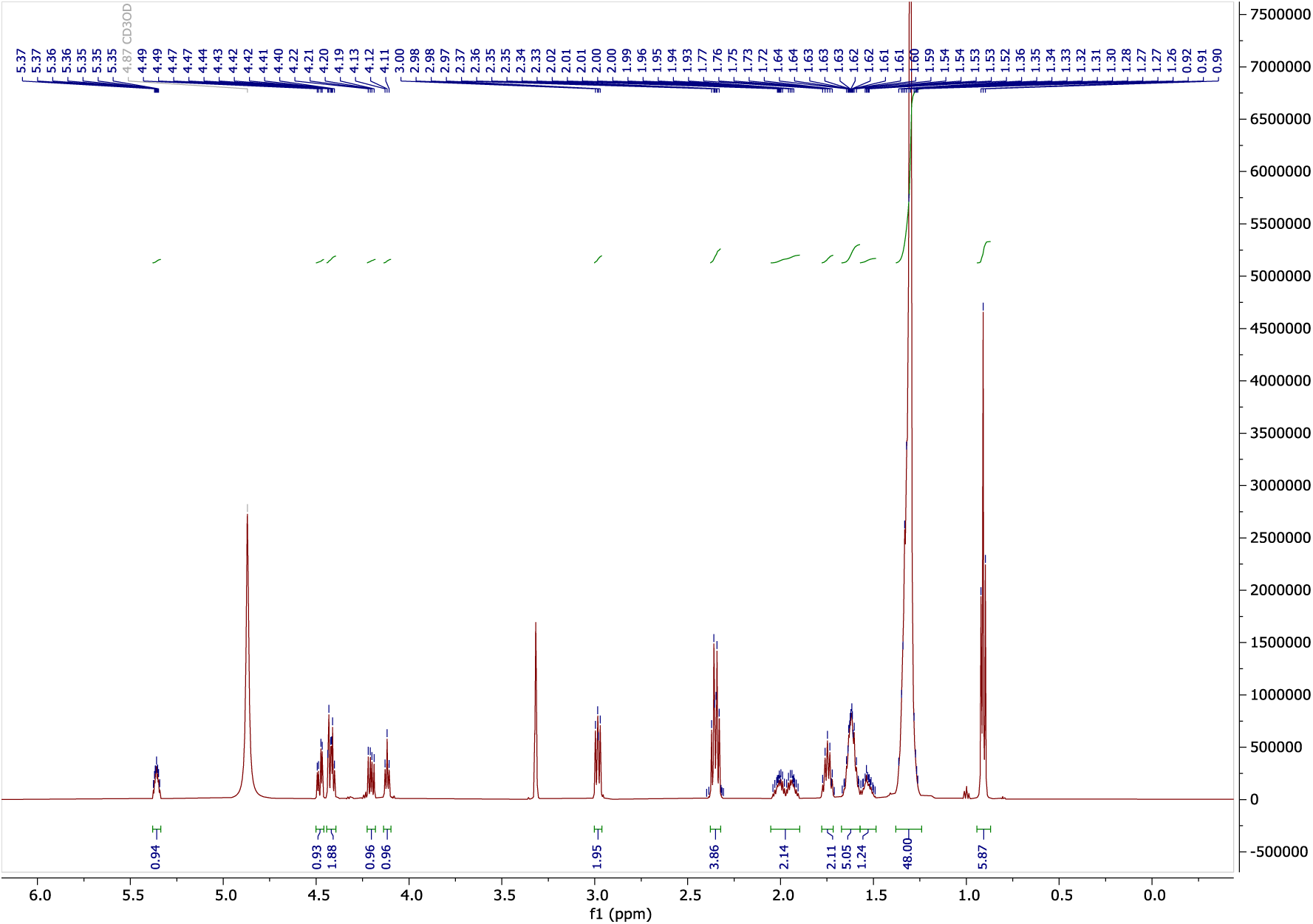

#### ^13^C-NMR of 5

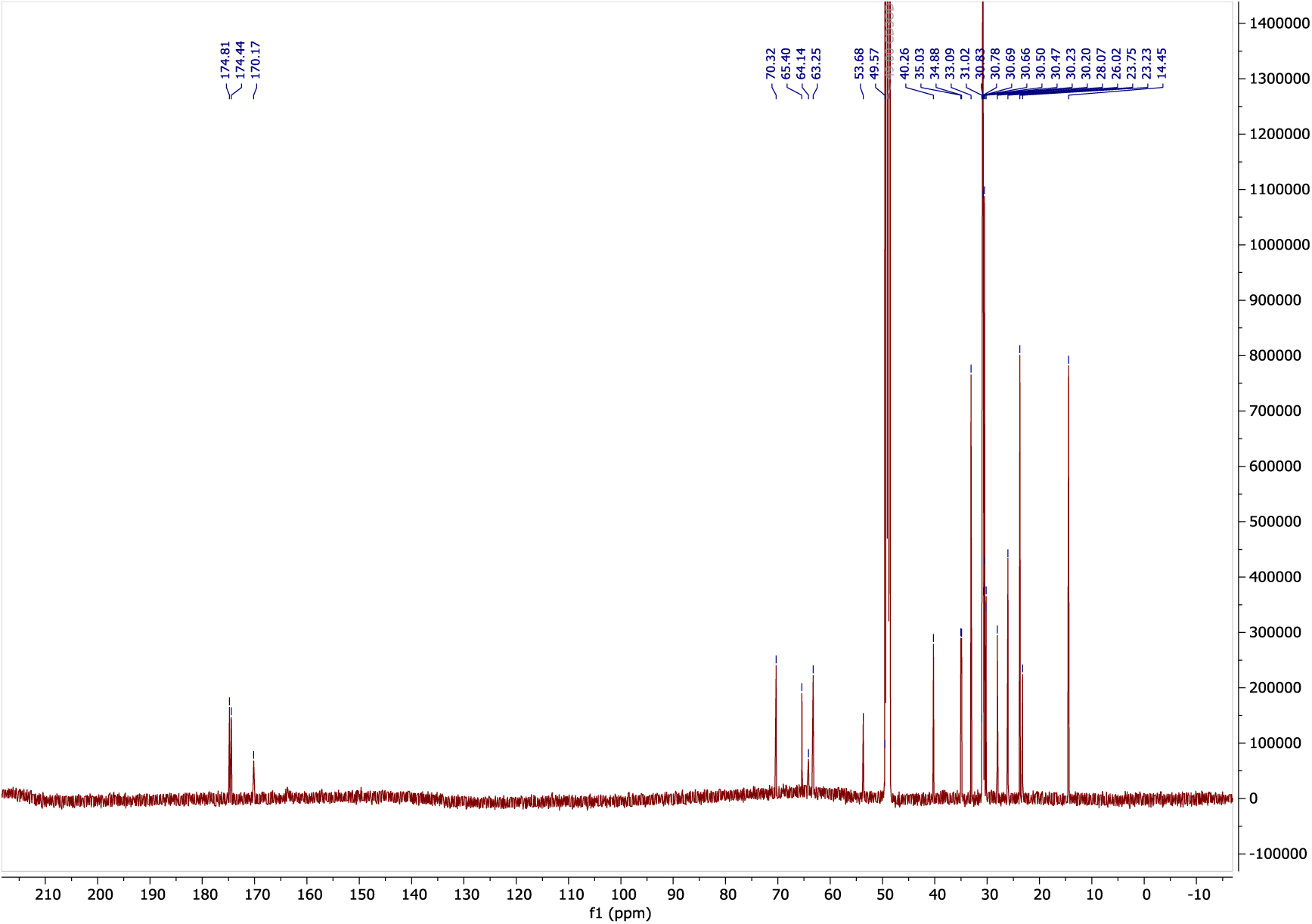

#### COSY of 5

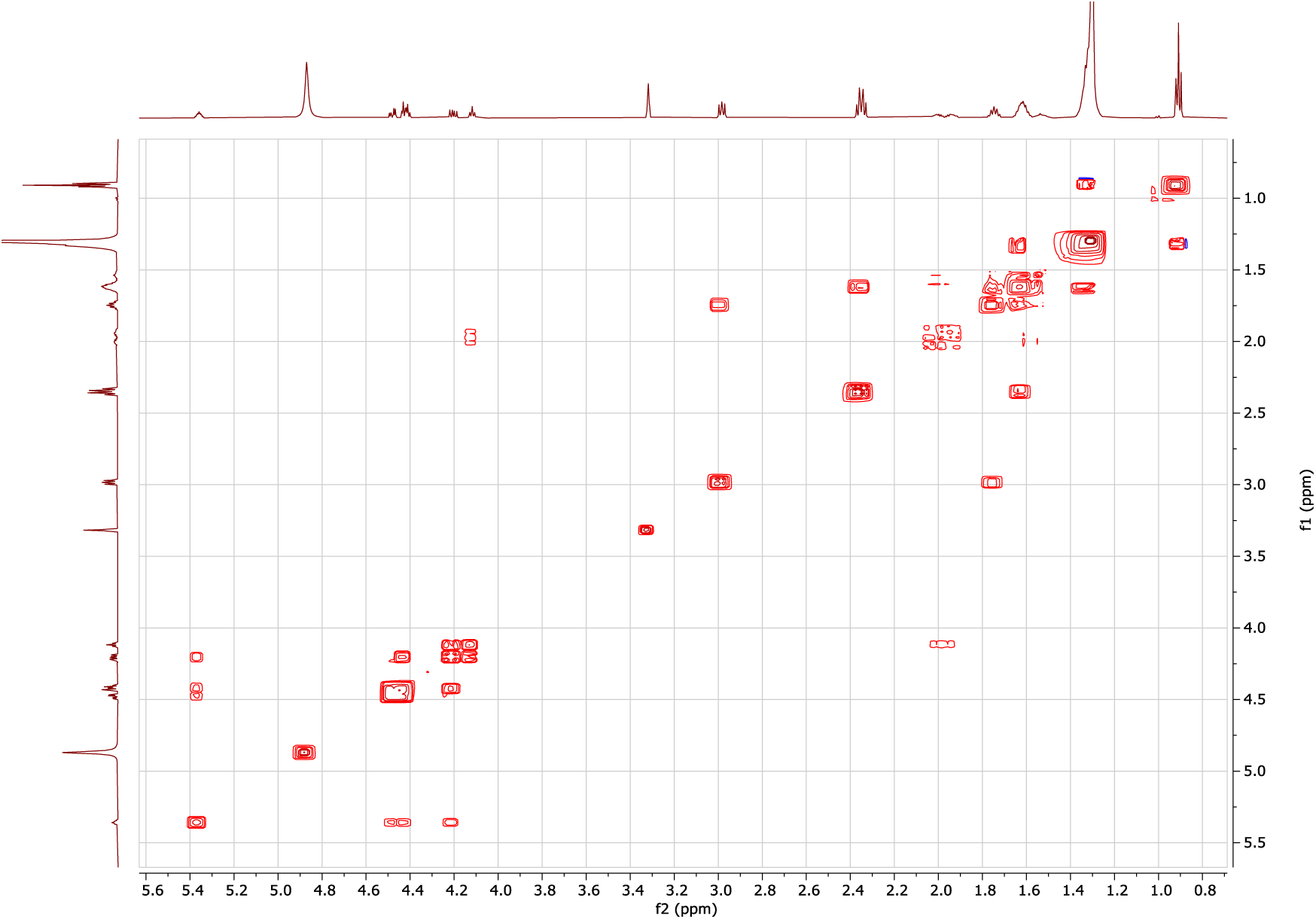

#### HSQC of 5

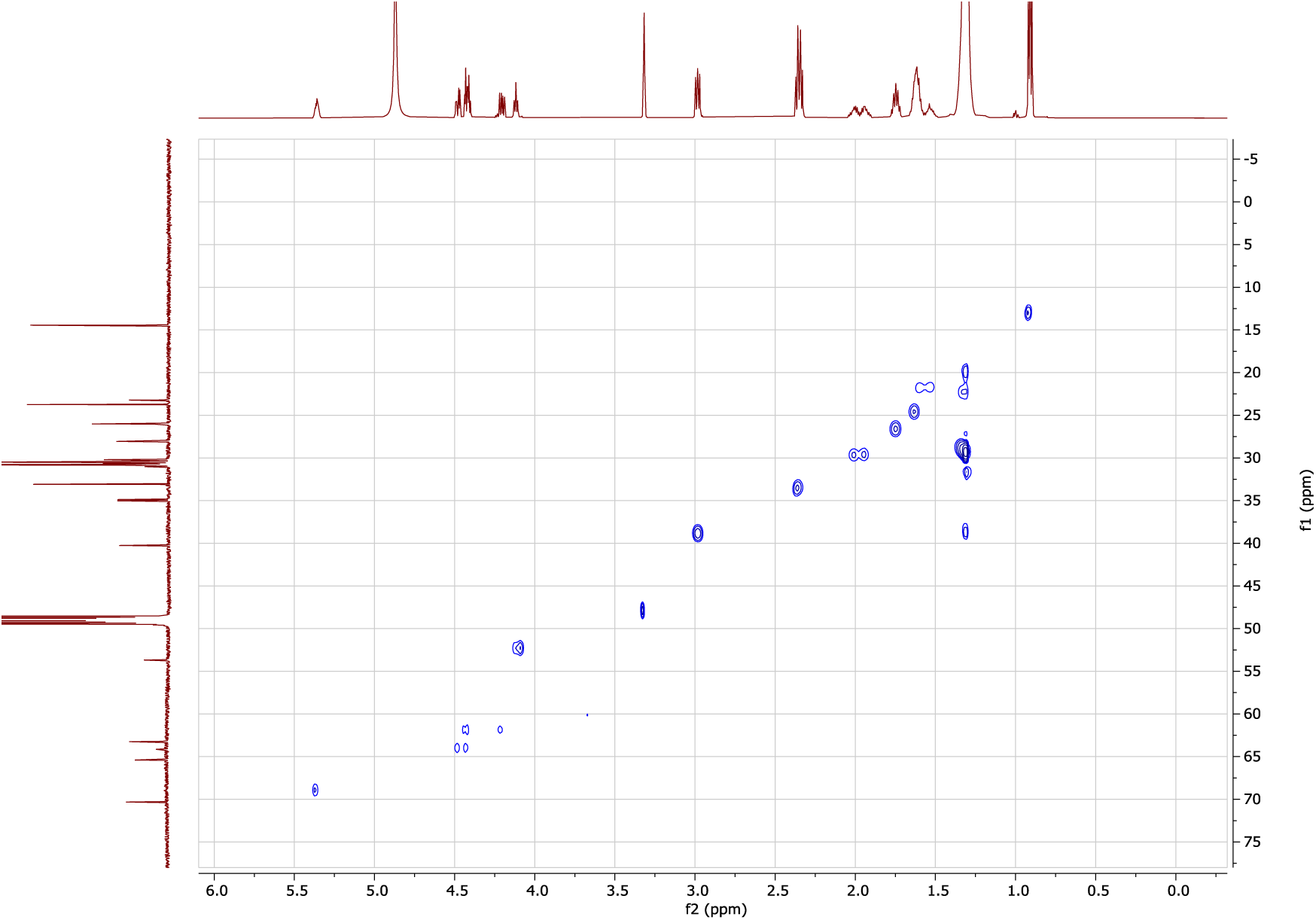

#### HMBC of 5

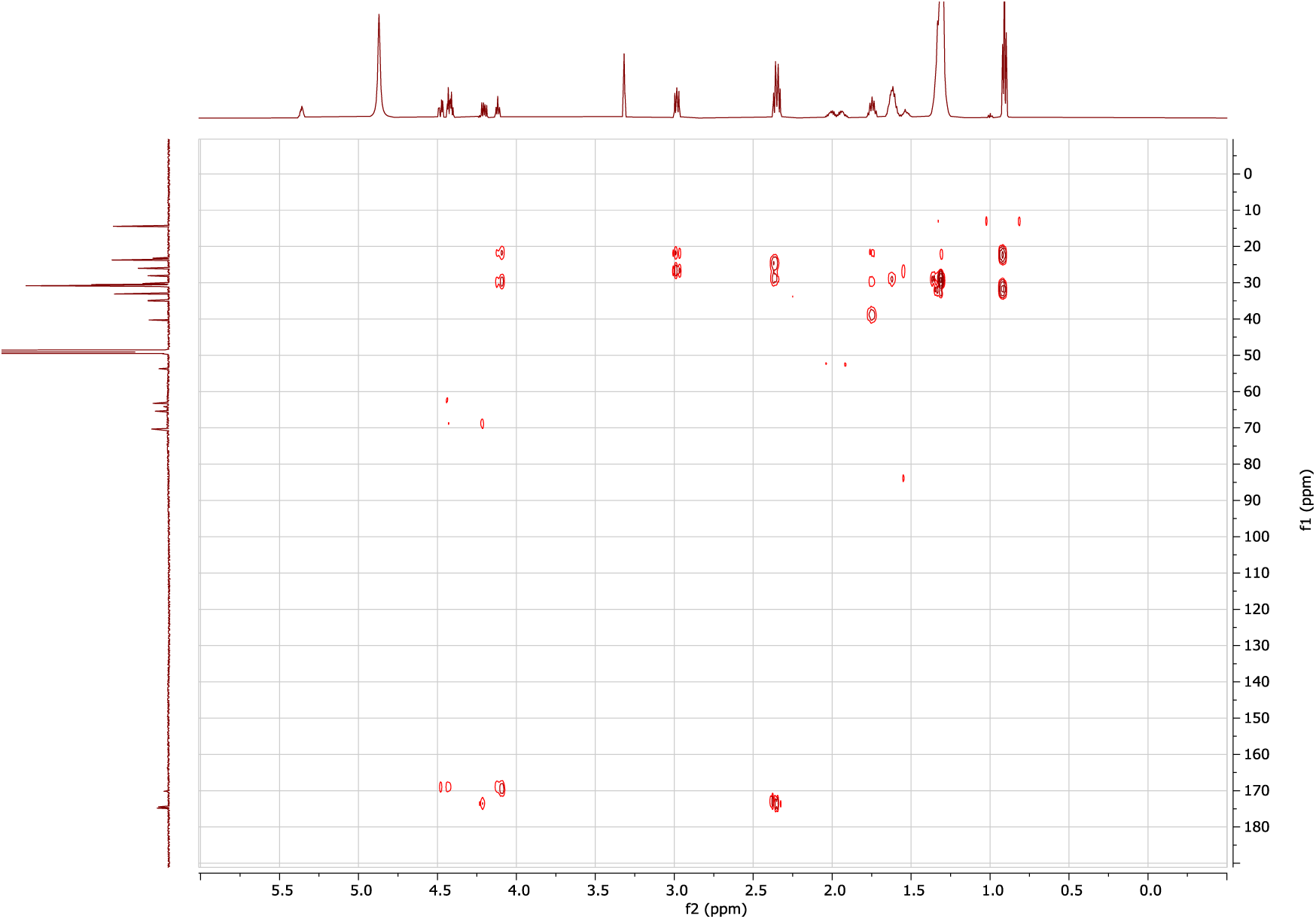

#### ESI-HRMS of 5

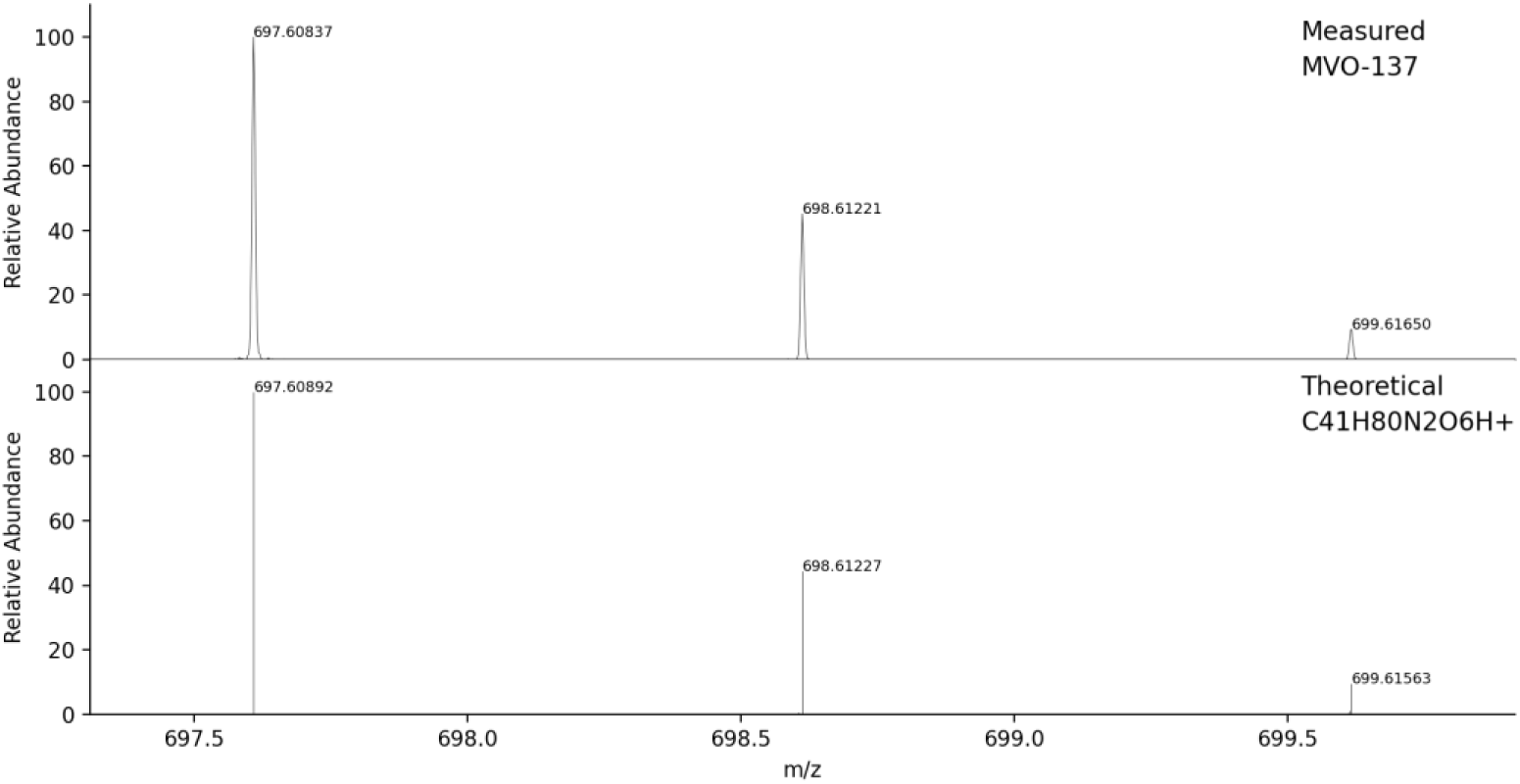

### Phylogeny of mycobacterial lysyldiacylglycerol production

Initial amino acid sequences for the generation of the phylogenetic tree of *lysX* and other aminoacyl transferases were obtained using Foldseek (*55*) of the Alphafold structure of MprF from *S. aureus* (Uniprot: Q2G2M2) and matching to the AFDB-SwissPROT database, yielding 152 sequences. An e-value for alignment threshold < 1 x 10^-12^ reduced the number of matching sequences to 41. After adding 10 sequences in bacteria of interest identified using Blastp, a final 51 sequences were aligned using Clustal Omega (*56*). The evolutionary analyses were conducted by the Maximum Likelihood Method in MEGA12 (*57*) using up to 6 parallel computing threads to identify the tree with the highest log likelihood (−56 974.80), and visualized using FigTree (available at: http://tree.bio.ed.ac.uk/software/figtree).

### *M. marinum lysX* mutant and complement

Disruption of the *lysX/*MMAR_2447 gene in *M. marinum* was confirmed through isolation and sequencing of a strain with a transposon insertion in MMAR_2447, 10.6% of the way through the *LysX* open reading frame, from a sequenced arrayed transposon library (kindly provided by C. Cosma and L. Ramakrishnan, University of Cambridge UK). Both the wildtype strain and transposon mutant were then transformed with *msp12:cerulean,* a plasmid constitutively expressing the fluorescent protein cerulean under the *msp12* promoter. For complementation, the entire *M. marinum lysX* open reading frame was cloned behind the *hsp60* promoter and inserted into the *msp12:cerulean* plasmid.

### Zebrafish husbandry and *M. marinum* infections

Zebrafish at two days post-fertilization were infected via the caudal vein as described (*58*) with 150–200 fluorescent bacteria per fish. All experiments had a minimum of 30 individuals per group on at least three different days to provide biologically and statistically robust data. Infection burden was measured at one day post-infection and four days post-infection through measurement of fluorescence. Fluorescence is enumerated by calculating the number of pixels above background using a constant threshold. Results were analyzed using GraphPad Prism, using Welch’s ANOVA followed by a Dunnett’s T3 multiple comparison of each group to the WT strain. Representative images of each group were chosen from the median values. Zebrafish husbandry and experimental procedures were performed in accordance and compliance with policies approved by the Duke University Institutional Animal Care and Use Committee (protocol A049-23-03).

---

**Fig. S1.**
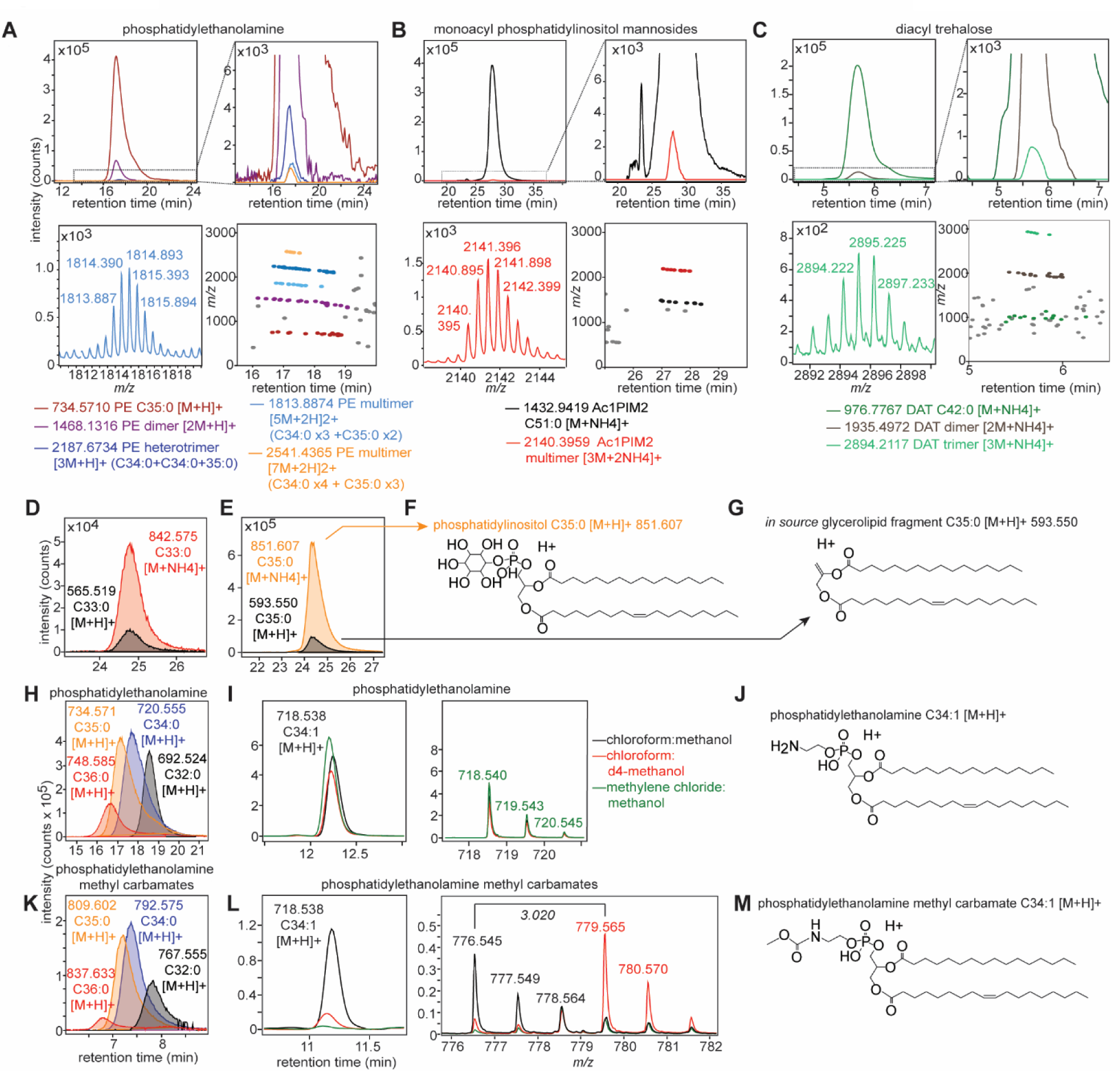
Identification of *in source* multimers, fragments and solvent-induced modification of known Mtb lipids. (**A**) phosphatidyl ethanolamine (PE), (**B**) monoacyl phosphatidylinositol dimannoside (Ac1PIM2) and (**C**) diacyl trehalose (DAT) multimer chromatograms were censored manually in the credentialling pipeline (Fig. 1F). The inset for each panel expands low-intensity multimer peaks, *top right*; representative mass spectra of multimer adducts, *bottom left*; and mapping the identified multimers onto retention time space, *bottom right*. (**D** to **G**) Representative (**D** and **E**) chromatograms and (**F**) structure of phosphatidylinositol (PI) acylforms with overlapping peaks that were consistent with (**G**) glycerolipid fragments (*black*). (**H**) Representative chromatograms of acylform distributions of PE in *M. marinum.* (**I**) Chromatograms, *left*, mass spectra, *right*, and (**J**) a representative structure show consistent detection of PE with modified solvents including deuterated methanol and methylene chloride for extraction. (**K**) Representative chromatograms of acylform distributions of PE methyl carbamates in *M. marinum*. (**L**) Chromatograms, *left*, mass spectra, *right*, and (**M**) a representative structure show loss of PE methyl carbamates with methylene chloride extraction and d4-methanol integration demonstrated by a mass shift of 3.020.

**Fig. S2.**
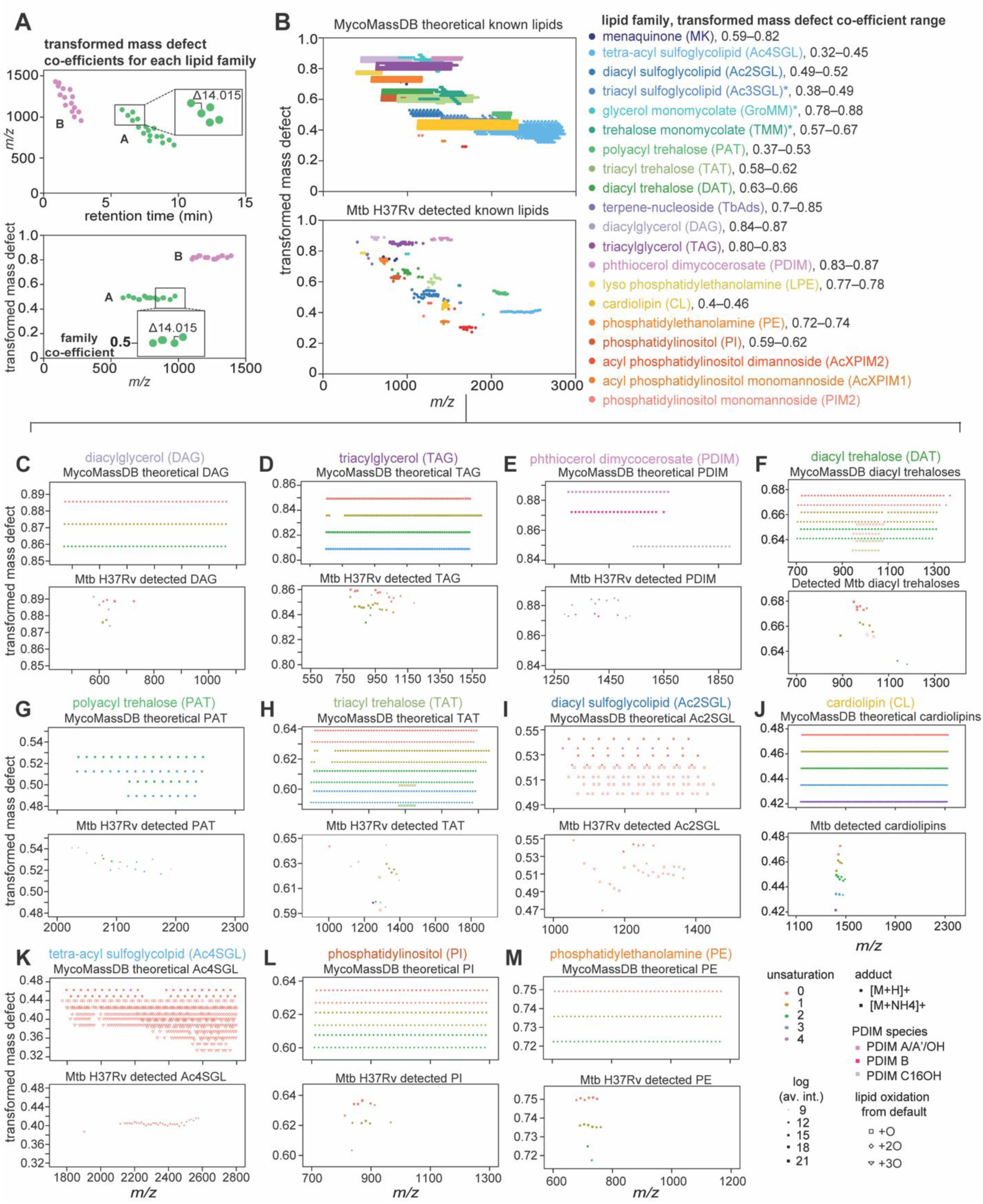
A methylene transformed mass defect of known *M. tuberculosis* lipids. (**A**) Schematic of example theoretical lipid families represented as *m/z* vs retention time, *top*, and as transformed or Kendrick mass defect vs *m/z*, *bottom*, to identify co-efficients for each lipid family. (**B**) Theoretical known lipids from MycoMassDB were transformed into a scale where related acylforms aligned along a horizontal axis, *top*. The detectable lipids from the credentialed Mtb H37Rv lipidome as in Figure 1I were also transformed showing narrower acylform distributions, *bottom*. The range of transformed mass defects of lipid families from MycoMassDB is appended to the legend. For each lipid family with alkylforms where a lead compound was confirmed by collisional MS a focused plot as in Fig. S2*A* shows the distribution of theoretical and detectable alkylforms, where differences in unsaturation, oxygenation, and ionization adduct are observed as horizontal displacements in (**C**) diacylglycerol, (**D**) triacylglycerol, (**E**) phthiocerol dimycocerosate, (**F**) diacyl trehalose, (**G**) polyacyltrehalose, (**H**) triacyl trehalose, (**I**) diacyl sulfoglycolipid, (**J**) cardiolipin, (**K**) tetra-acyl sulfoglycolipid, (**L**) phosphatidyl inositol, and (**M**) phosphatidyl ethanolamine. Point size was scaled to the log of average intensity across the matched quadruplicate cultures of Mtb H37Rv analyzed. Lipids with asterisks mass matched to MycoMassDB and were not studied by CID-MS.

**Fig. S3.**
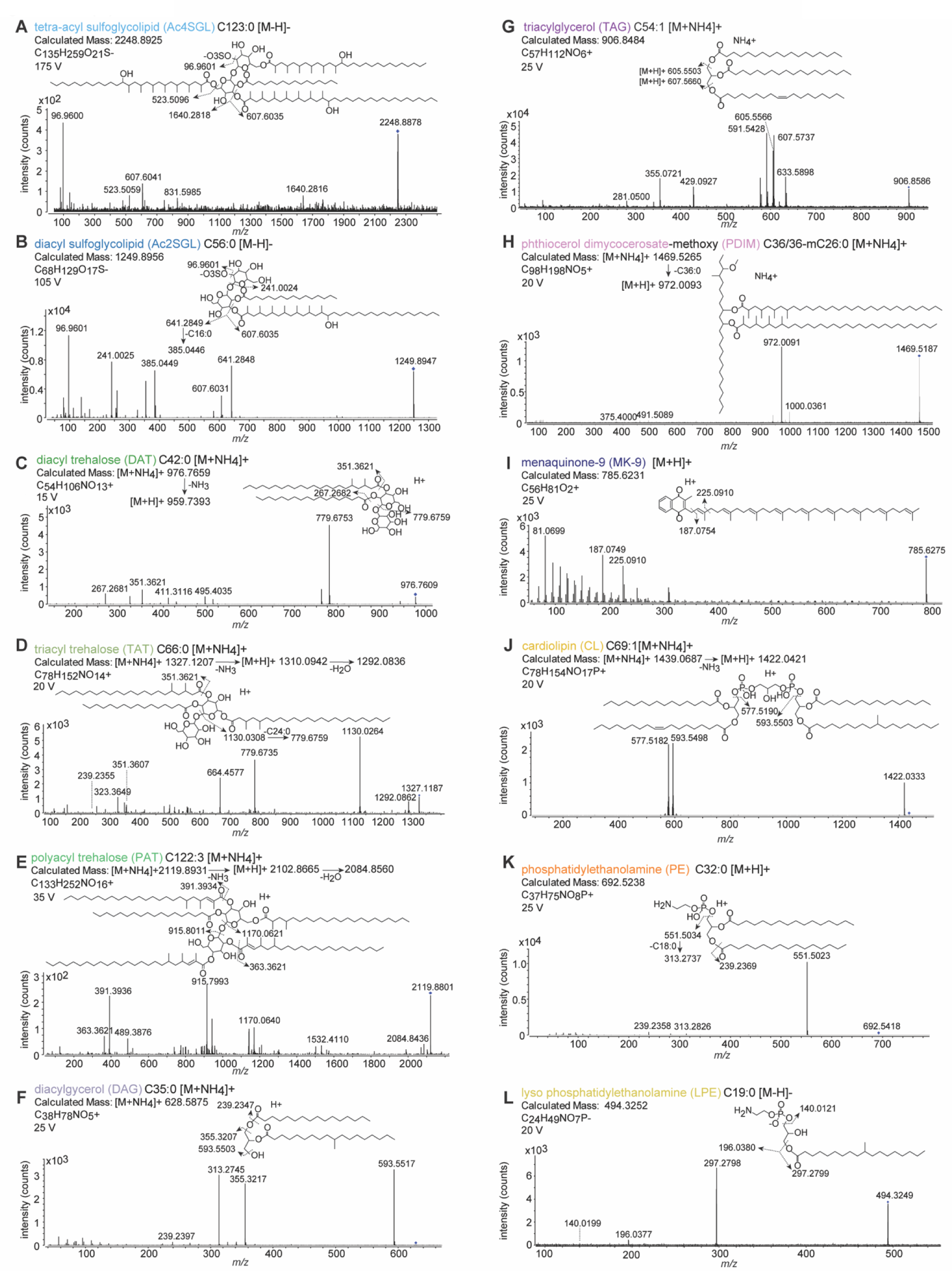
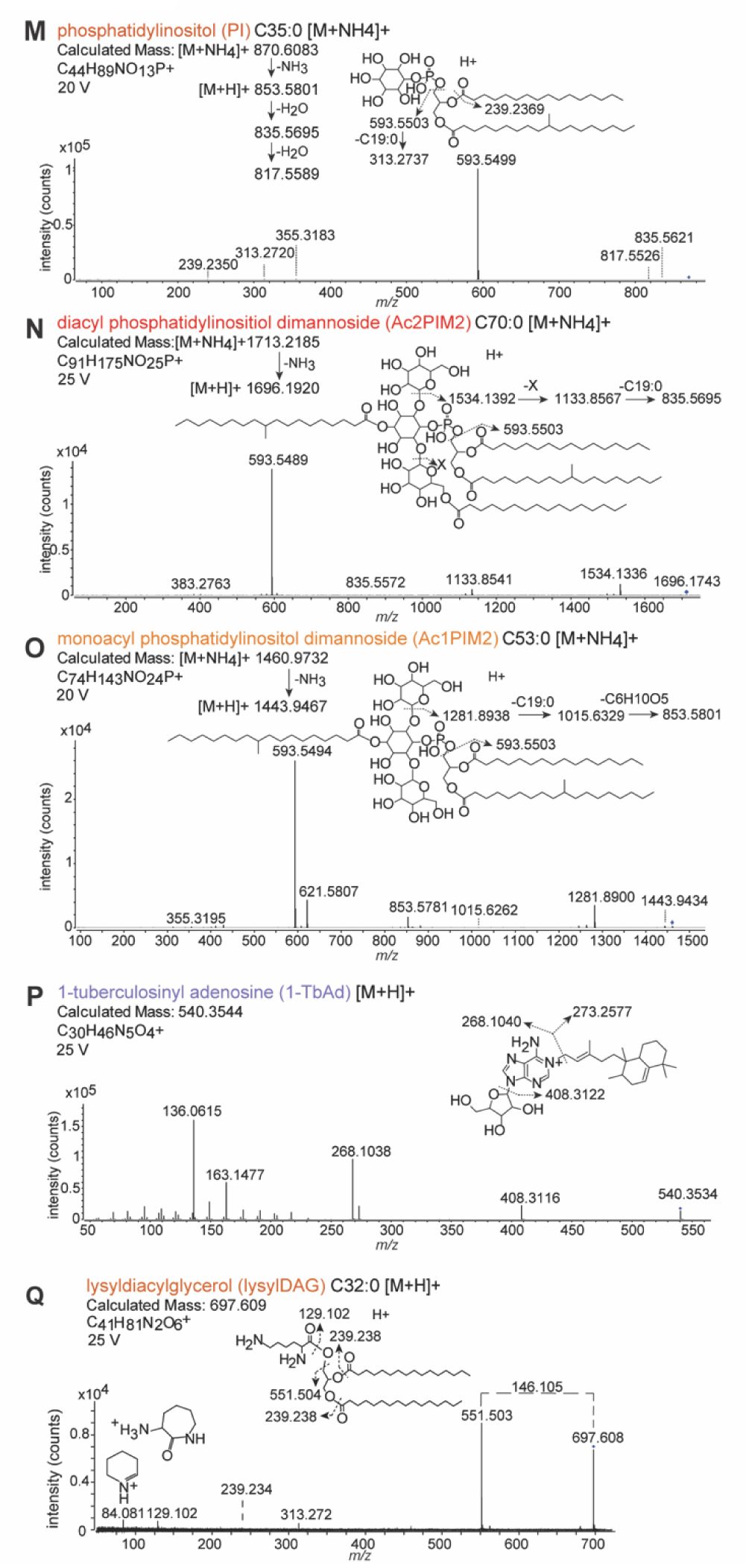
Collisional MS spectra of *M. tuberculosis* lipids. Collisional MS was performed on lead compounds from the ‘lipidome of knowns’ in either the negative or positive modes with a voltage optimized to each compound. A representative chemical structure and overlay of fragments detected is shown for (**A**) tetra-acyl sulfoglycolipid, (**B**) diacyl sulfoglycolipid, (**C**) diacyl trehalose, (**D**) triacyl trehalose, (**E**) polyacyl trehalose, (**F**) diacylglycerol, (**G**) triacylglycerol, (**H**) phthiocerol dimycocerosate, (**I**) menaquinone, (**J**) cardiolipin, (**K**) phosphatidyl ethanolamine, (**L**) lyso phosphatidylethanolamine, (**M**) phosphatidyl inositol, (**N**) diacyl phosphatidylinositol dimannoside, (**O**) monoacyl phosphatidylinositol dimannoside, and (**P**) tuberculosinyl adenosine. A representative chemical structure and overlay of fragments detected for previously unknown (**Q**) lysyldiacylglycerol, identified in these studies from the ‘lipidome of unknowns’.

**Fig. S4.**
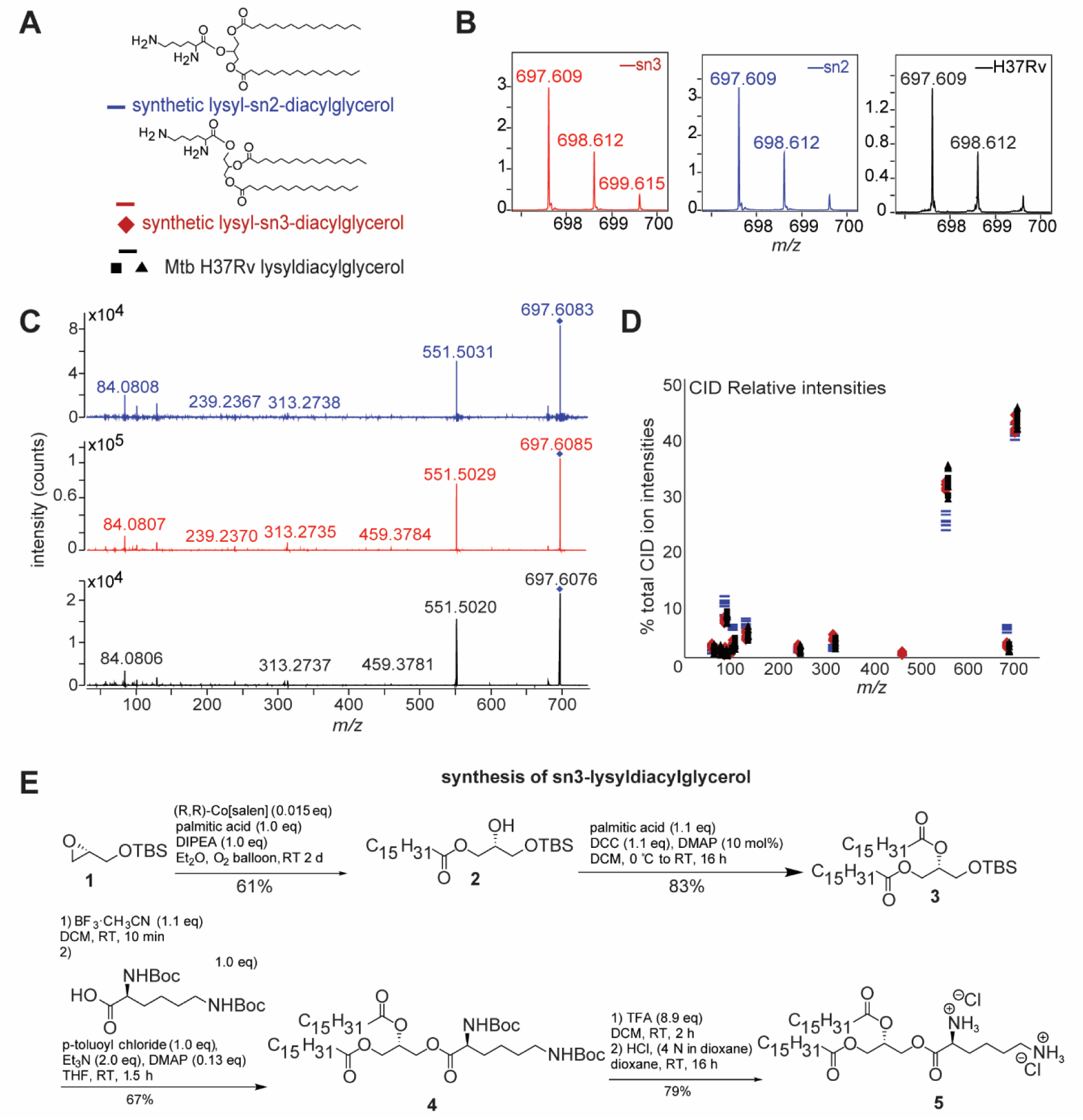
Comparison of synthesized and natural lysyldiacylglycerol. (**A**) Chemical structures of synthesized lysydiacylglycerol with the lysine head group at the *sn2* (*blue*) or *sn3* (*red*) position serve as a legend for panels C to F below. (**B**) Mass spectra of synthesized and natural lysyldiacylglycerol show identical detected masses and isotope distributions and (**C**) identical CID-MS fragments between synthesized isomers and the natural product (< 10 ppm). However, (**D**) the proportion of intensities of fragments of the Mtb H37Rv lysyldiacylglycerol (*black*) matches the synthetic *sn3* isomer (*red*) and not the *sn2* isomer (*blue*) further confirming the natural product structure. (**E**) A simplified chemical synthesis for *sn3* dipalmityl lysyldiacylglycerol is shown, with representative structures of intermediates as SI Methods.

**Fig. S5.**
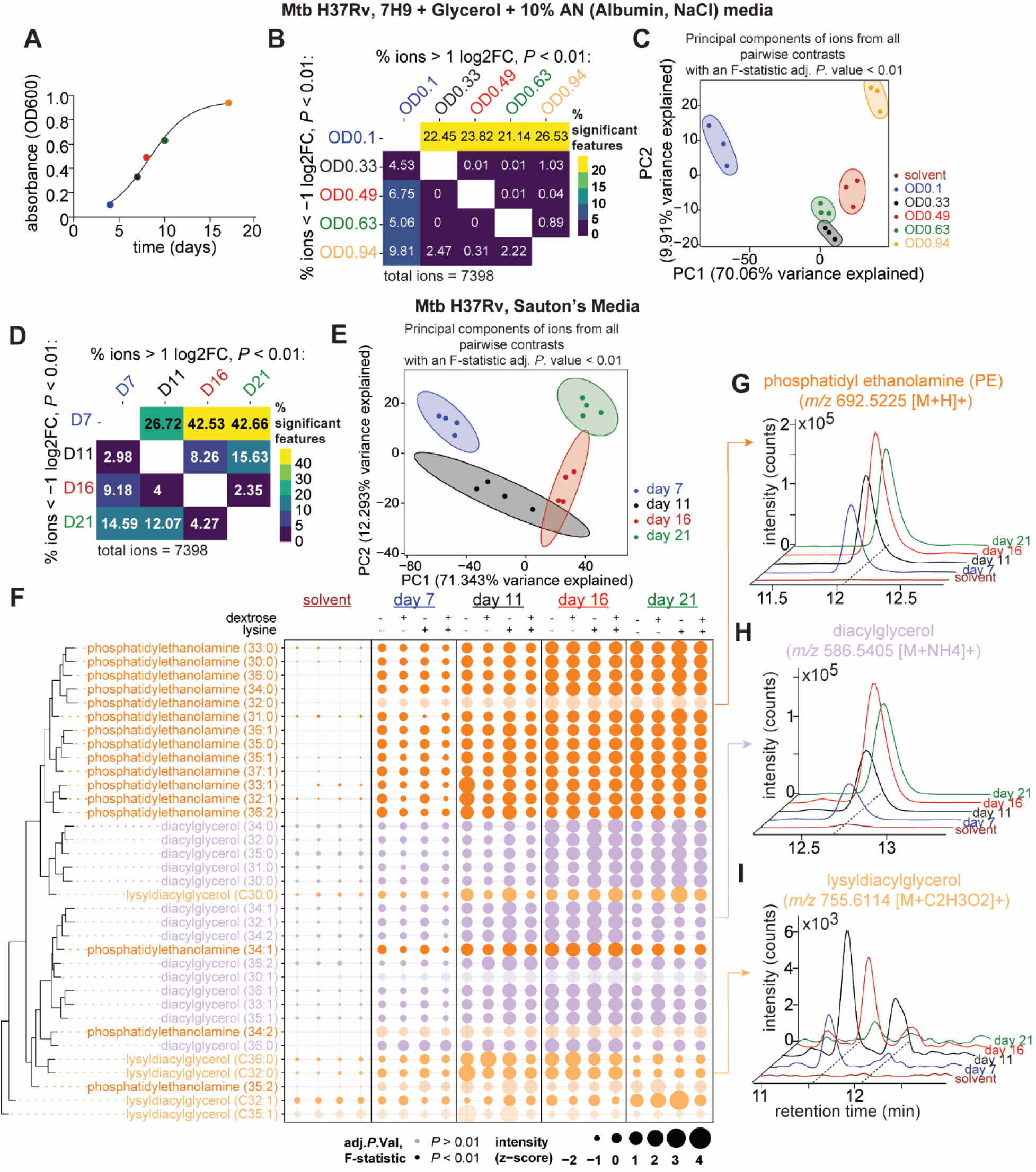
Identification of lysyldiacylglycerol in 7H9 and Sauton’s media. Mtb H37Rv was grown in (**A** to **C**) matched triplicate cultures in 7H9 media with 10% Albumin and NaCl or (**D** to **H**) in detergent-free Sauton’s media with glycerol as a sole carbon source in parallel quadruplicate cultures supplemented with either dextrose, lysine or both. (**A**) A matched 0.05% Tween-80 culture in dextrose-free 7H9 was used to determine the optical density at 600 nm at various timepoints as a measure of growth phase. (**B** and **D**) A lipidomic contrast of all pairwise comparisons identified the number of significant lipids with > two-fold change and a Benjamini-Hochberg adjusted *P* value < 0.01 contrasting each time point sampled. These non-solvent contrasts informed the F-statistic used to determine a significance threshold for variability in Figure 2L and Figure S5F respectively. (**C** and **E**) The variance between timepoints was explained using principal components analysis of lipids meeting the significance threshold for variation, *P* value of the F-statistic < 0.01. (**F**) The z-score intensities plotted as bubbles showed the distribution of known acylforms of glycerolipids across growth timepoints in dextrose-free Sauton’s media. A significant *P* value of the F-statistic < 0.01, identified variable lipids across growth phase as darkened circles, with representative single ion chromatograms of (**G**) phosphatidyl ethanolamine, (**H**) diacylglycerol, and (**I**) lysyldiacylglycerol.

**Fig. S6.**
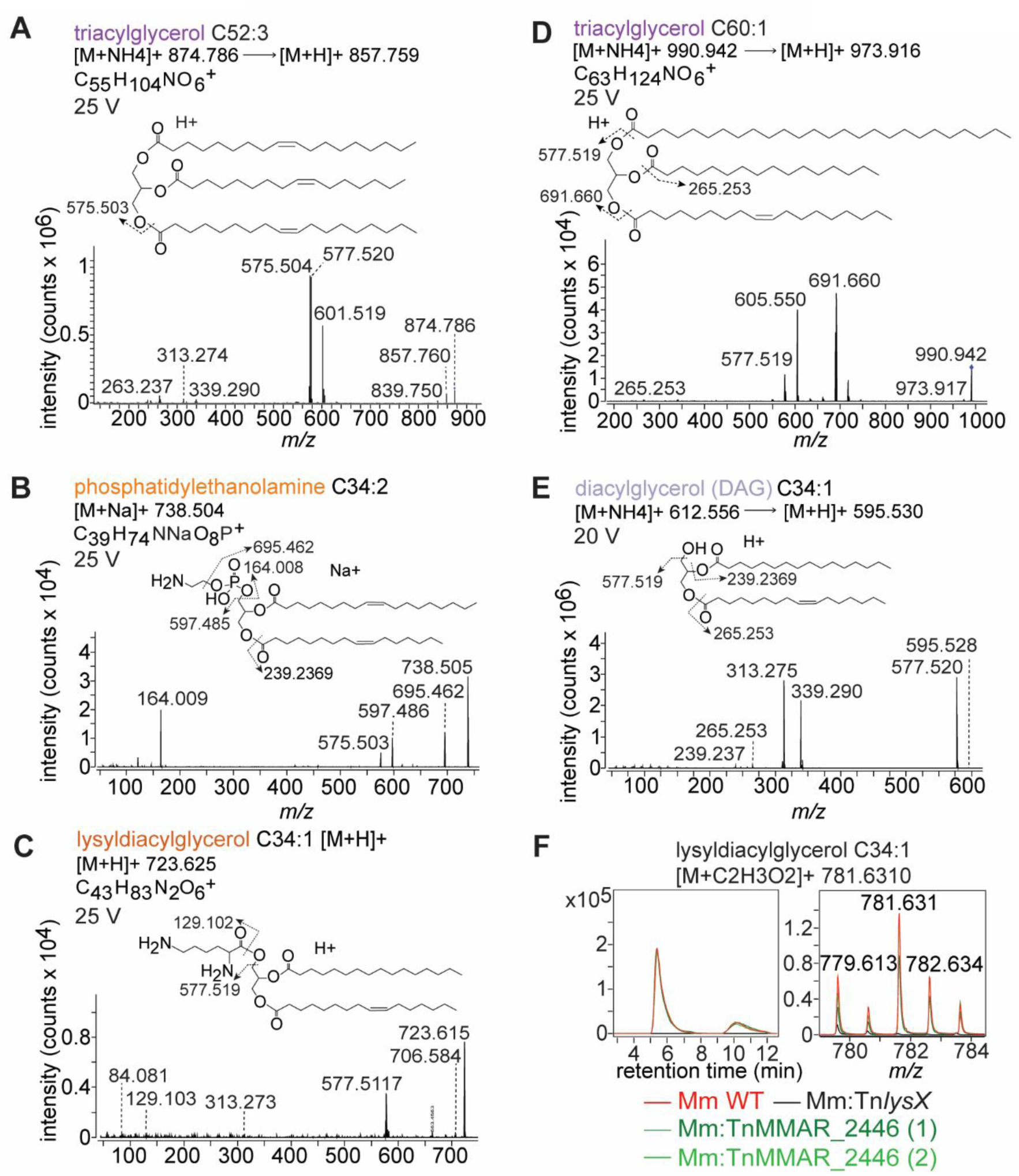
Collisional MS of *M. marinum* lipids. Collisional MS spectrum and the interpreted fragments overlaid on a representative molecular structure for (**A** and **D**) triacylglycerol, (**B**) phosphatidylethanolamine, (**C**) lysyldiacylglycerol, and (**E**) diacylglycerol in the positive modes with collisional energy indicated for each species identified in the *M. marinum lysX*-dependent lipidome. (**F**) Disruption of the *lysX* downstream gene Rv1639c/MMAR_2246, did not affect levels of lysylDAG represented as chromatograms in the normal phase (*left*) or mass spectrum (*right*), representative of biological quadruplicate cultures. Two independent TnMMAR_2446 mutants were tested.

**Fig. S7.**
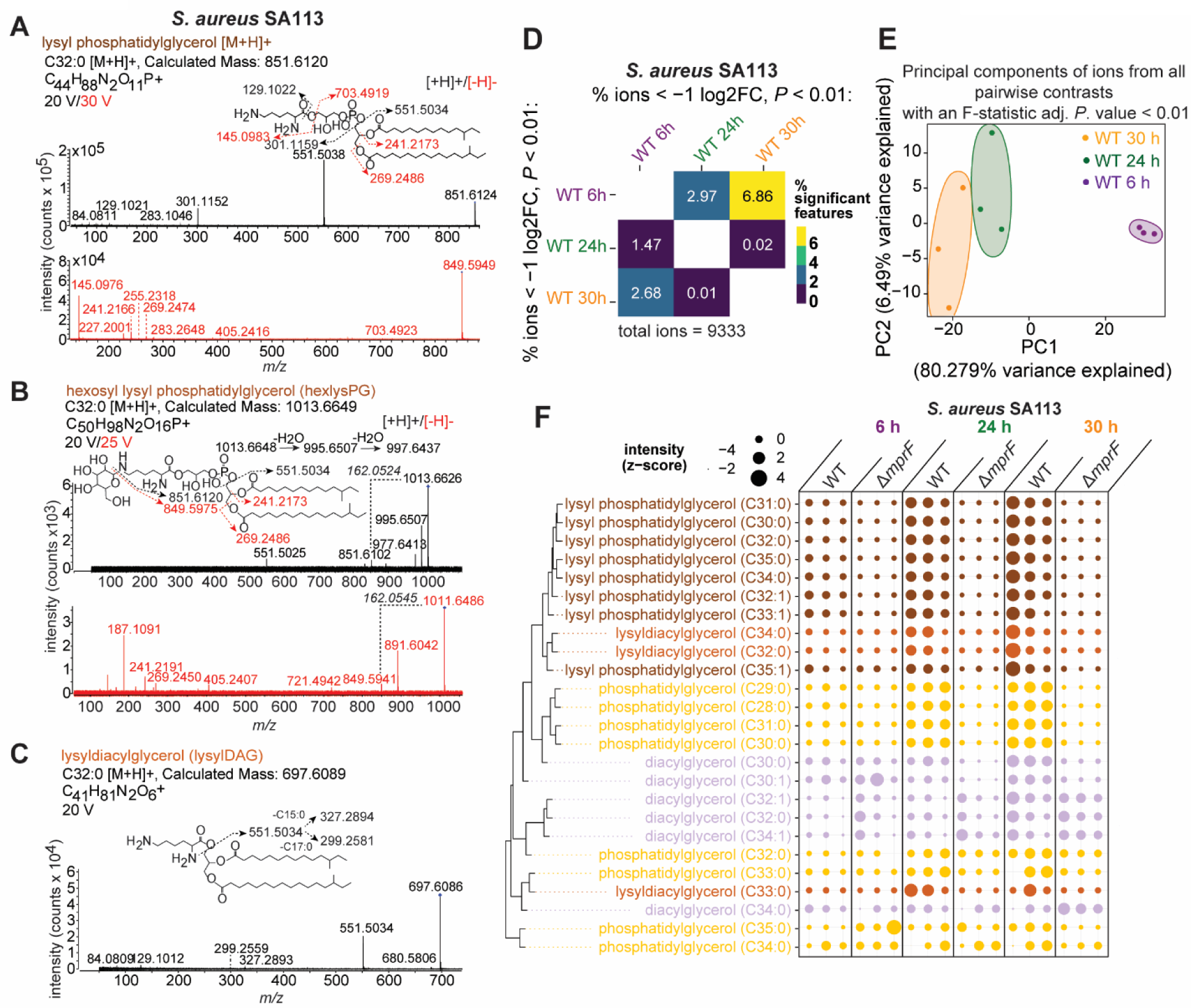
Collisional MS spectra for *mprF*-dependent lipids. Collisional MS spectrum and the interpreted fragments overlaid on a representative molecular structure in the positive (*black*) and negative modes (*red*) for the C32:0 acylforms of (**A**) lysyl phosphatidylglycerol, (**B**) hexosyl lysyl phosphatidylglycerol, and (**C**) lysyldiacylglycerol in *S. aureus* SA113 lipid extracts. (**D**) A tileplot of all pairwise contrasts between SA 113 WT strains grown at 3 timepoints show the percentage of significant lipids enriched in each phase. (**E**) The variance between timepoints was explained using principal components analysis of lipids meeting the significance threshold for variation, F-statistic adj. *P*. value < 0.01. (**F**) The z-score intensity of lysine and neutral glycerolipids in *S. aureus* SA113 shows *mprF*-dependence of both 8 acylforms of lysylPG and 3 lysylDAG across 3 timepoints in biological triplicate, in contrast to 6 diacylglycerol (DAG).

**Data S1.** Replication R-Markdown R markdown (lines 1 - 2145) and 6 supporting csv files, 10 phenotype objects, 12 *xcms* objects, and 1 table, required with formatted datasets S2-S13 to generate all R-based analysis and figures, including the feature credentialling pipeline. A second R markdown includes 4 custom functions to interface with R packages *xcms* and *limms*, and a new function *mzrtMeta* to align mass spectrometry experiments sourced by the replication markdown (available at: https://github.com/jamayfie/mzrtMatch). Users must modify path statements to their own data drive with downloaded data files below. Additionally all R packages, dependencies, and custom functions must be sourced and loaded.

**Data S2.** MycoMassDB Mycobacterial database of validated lipids and metabolites, including newly discovered lysine lipoamino acids, formatted for automated matching.

**Data S3.** MycoLOBSTAH Mycobacterial database of theoretical lipids in the positive mode propagated from MycoMassDB using LOBSTAHs R-package.

**Data S4.** Credentialed map of known mycobacterial lipids and lysyldiacylglycerols Map of Mtb H37Rv qualified features in two groups: known lipids with identities confirmed by collisional MS, and unknown features ranked by quality criteria.

**Data S5.** *lysX*/*mprF* domain ontology *M. tuberculosis* genes with homology to firmicute *mprF*, including their subdomains and Interpro classifications.

**Data S6.** Lipids knocked down with lysX silencing in Mtb H37Rv Lipids significantly enriched in the Mtb H37Rv *lysX* CRISPRi knockdown uninduced (-ATc) condition compared to the induced strain (+ATc).

**Data S7.** Mtb H37Rv growth phase glycerolipids, 7H9 Acylform intensities of diacylglycerol, lysyldiacylglycerol and phosphatidylethanolamine detected across growth phases in Mtb H37Rv grown in 7H9 media.

**Data S8.** Mtb H37Rv growth phase glycerolipids, Sauton’s Acylform intensities of diacylglycerol, lysyldiacylglycerol and phosphatidylethanolamine detected across growth phases in Mtb H37Rv grown in Sauton’s media.

**Data S9.** Distribution of lysylDAG and other glycerolipids across mycobacteria Aligned dataset of acylform intensities of diacylglycerol, lysyldiacylglycerol and phosphatidylethanolamine across 15 mycobacterial strains and species.

**Data S10.** Lipids enriched in the Mm *lysX* mutant Lipids meeting significance criteria as enriched in the Mm *lysX* mutant from a compound contrast against Mm WT and complement.

**Data S11.** Lipids enriched in Mm WT and *lysX* complement Lipids meeting significance criteria as enriched in the Mm WT and *lysX* complement from a compound contrast against the Mm *lysX* mutant.

**Data S12.** *mprF*-dependent lipids in *S. aureus* Lipids meeting significance criteria as enriched in *S. aureus* WT compared to the Δ*mprF* mutant

**Data S13.** *S. aureus* SA113 growth phase glycerolipids Acylform intensities of diacylglycerol, lysyldiacylglycerol and lysyl phosphatidylglycerol detected across growth phases in *S. aureus* SA 113 and the Δ*mprF* mutant

## References

1. D. G. Russell, C. E. Barry, J. L. Flynn, Tuberculosis: What We Don’t Know Can, and Does, Hurt Us. Science (2010). 328, 852–856 (2010) doi:10.1126/science.1184784.

2. C. L. Dulberger, E. J. Rubin, C. C. Boutte, The mycobacterial cell envelope — a moving target. Nat Rev Microbiol 18, 47–59 (2020) doi:10.1038/s41579-019-0273-7.

3. E. Layre, L. Sweet, S. Hong, C. A. Madigan, D. Desjardins, D. C. Young, T. Y. Cheng, J. W. Annand, K. Kim, I. C. Shamputa, M. J. McConnell, C. A. Debono, S. M. Behar, A. J. Minnaard, M. Murray, C. E. Barry, I. Matsunaga, D. B. Moody, A comparative lipidomics platform for chemotaxonomic analysis of mycobacterium tuberculosis. Chem Biol 18, 1537–1549 (2011) doi:10.1016/j.chembiol.2011.10.013.

4. C. R. Ruhl, B. L. Pasko, H. S. Khan, L. M. Kindt, C. E. Stamm, L. H. Franco, C. C. Hsia, M. Zhou, C. R. Davis, T. Qin, L. Gautron, M. D. Burton, G. L. Mejia, D. K. Naik, G. Dussor, T. J. Price, M. U. Shiloh, Mycobacterium tuberculosis Sulfolipid-1 Activates Nociceptive Neurons and Induces Cough. Cell 181, 293–305.e11 (2020) doi:10.1016/j.cell.2020.02.026.

5. A. M. Block, S. B. Namugenyi, N. P. Palani, A. M. Brokaw, L. Zhang, K. B. Beckman, A. D. Tischler, Mycobacterium tuberculosis Requires the Outer Membrane Lipid Phthiocerol Dimycocerosate for Starvation-Induced Antibiotic Tolerance. mSystems 8, e0069922 (2023) doi:10.1128/msystems.00699-22.

6. E. Mittal, G. V. R. K. Prasad, S. Upadhyay, J. Sadadiwala, A. J. Olive, G. Yang, C. M. Sassetti, J. A. Philips, Mycobacterium tuberculosis virulence lipid PDIM inhibits autophagy in mice. Nat Microbiol 9, 2970–2984 (2024) doi:10.1038/s41564-024-01797-5.

7. C. J. Cambier, K. K. Takaki, R. P. Larson, R. E. Hernandez, D. M. Tobin, K. B. Urdahl, C. L. Cosma, L. Ramakrishnan, Mycobacteria manipulate macrophage recruitment through coordinated use of membrane lipids. Nature 505, 218–222 (2014) doi:10.1038/nature12799.

8. C. A. Madigan, T. Y. Cheng, E. Layre, D. C. Young, M. J. McConnell, C. A. Debono, J. P. Murry, J. R. Wei, C. E. Barry, G. M. Rodriguez, I. Matsunaga, E. J. Rubin, D. B. Moody, Lipidomic discovery of deoxysiderophores reveals a revised mycobactin biosynthesis pathway in Mycobacterium tuberculosis. Proc Natl Acad Sci U S A 109, 1257–1262 (2012) doi:10.1073/pnas.1109958109.

9. E. Ishikawa, T. Ishikawa, Y. S. Morita, K. Toyonaga, H. Yamada, O. Takeuchi, T. Kinoshita, S. Akira, Y. Yoshikai, S. Yamasaki, Direct recognition of the mycobacterial glycolipid, trehalose dimycolate, by C-type lectin Mincle. J Exp Med 206, 2879 (2009) doi:10.1084/JEM.20091750.

10. J. Buter, T. Y. Cheng, M. Ghanem, A. E. Grootemaat, S. Raman, X. Feng, A. R. Plantijn, T. Ennis, J. Wang, R. N. Cotton, E. Layre, A. K. Ramnarine, J. A. Mayfield, D. C. Young, A. Jezek Martinot, N. Siddiqi, S. Wakabayashi, H. Botella, R. Calderon, M. Murray, S. Ehrt, B. B. Snider, M. B. Reed, E. Oldfield, S. Tan, E. J. Rubin, M. A. Behr, N. N. van der Wel, A. J. Minnaard, D. B. Moody, Mycobacterium tuberculosis releases an antacid that remodels phagosomes. Nat Chem Biol 15, 889–899 (2019) doi:10.1038/s41589-019-0336-0.

11. B. N. Koleske, W. R. Jacobs, W. R. Bishai, The Mycobacterium tuberculosis genome at 25 years: lessons and lingering questions. J Clin Invest 133 (2023) doi:10.1172/JCI173156.

12. E. Layre, H. J. Lee, D. C. Young, A. J. Martinot, J. Buter, A. J. Minnaard, J. W. Annand, S. M. Fortune, B. B. Snider, I. Matsunaga, E. J. Rubin, T. Alber, D. B. Moody, Molecular profiling of Mycobacterium tuberculosis identifies tuberculosinyl nucleoside products of the virulence-associated enzyme Rv3378c. Proc Natl Acad Sci U S A 111, 2978–2983 (2014) doi:10.1073/pnas.1315883111.

13. J. D. Mougous, R. H. Senaratne, C. J. Petzold, M. Jain, D. H. Lee, M. W. Schelle, M. D. Leavell, J. S. Cox, J. A. Leary, L. W. Riley, C. R. Bertozzi, A sulfated metabolite produced by stf3 negatively regulates the virulence of Mycobacterium tuberculosis. Proc Natl Acad Sci U S A 14, 4258–4263 (2006) doi:10.1073/pnas.0510861103.

14. M. J. Sartain, D. L. Dick, C. D. Rithner, D. C. Crick, J. T. Belisle, Lipidomic analyses of Mycobacterium tuberculosis based on accurate mass measurements and the novel “Mtb LipidDB.” J Lipid Res 52, 861–872 (2011) doi:10.1194/jlr.M010363.

15. H. C. Leier, J. B. Weinstein, J. E. Kyle, J.-Y. Lee, L. M. Bramer, K. G. Stratton, D. Kempthorne, A. R. Navratil, E. G. Tafesse, T. Hornemann, W. B. Messer, E. A. Dennis, T. O. Metz, E. Barklis, F. G. Tafesse, A global lipid map defines a network essential for Zika virus replication. Nat Commun 11, 3652 (2020) doi:10.1038/s41467-020-17433-9.

16. J. A. Mayfield, S. Raman, A. K. Ramnarine, V. K. Mishra, A. D. Huang, S. Dudoit, J. Buter, T.-Y. Cheng, D. C. Young, Y. M. Nair, I. G. Ouellet, B. T. Griebel, S. Ma, D. R. Sherman, L. Mallet, K. Y. Rhee, A. J. Minnaard, D. Branch Moody, Mycobacteria that cause tuberculosis have retained ancestrally acquired genes for the biosynthesis of chemically diverse terpene nucleosides. PLoS Biol 22, e3002813 (2024) doi:10.1371/journal.pbio.3002813.

17. M. Sindelar, G. J. Patti, Chemical Discovery in the Era of Metabolomics. J Am Chem Soc 142, 9097–9105 (2020) doi:10.1021/jacs.9b13198.

18. M. Giera, A. Aisporna, W. Uritboonthai, G. Siuzdak, The hidden impact of in-source fragmentation in metabolic and chemical mass spectrometry data interpretation. Nat Metab 6, 1647–1648 (2024) doi:10.1038/s42255-024-01076-x.

19. N. Zamboni, A. Saghatelian, G. J. Patti, Defining the Metabolome: Size, Flux, and Regulation. Mol Cell 58, 699–706 (2015) doi:10.1016/j.molcel.2015.04.021.

20. N. G. Mahieu, X. Huang, Y. J. Chen, G. J. Patti, Credentialing features: A platform to benchmark and optimize untargeted metabolomic methods. Anal Chem 86, 9583–9589 (2014) doi:10.1021/ac503092d.

21. J. R. Collins, B. R. Edwards, H. F. Fredricks, B. A. S. Van Mooy, LOBSTAHS: An Adduct-Based Lipidomics Strategy for Discovery and Identification of Oxidative Stress Biomarkers. Anal Chem 88, 7154–7162 (2016) doi:10.1021/acs.analchem.6b01260.

22. C. Kuhl, R. Tautenhahn, C. Böttcher, T. R. Larson, S. Neumann, CAMERA: An integrated strategy for compound spectra extraction and annotation of liquid chromatography/mass spectrometry data sets. Anal Chem 84, 283–289 (2012) doi:10.1021/ac202450g.

23. T. A. Garrett, C. R. H. Raetz, J. D. Son, T. D. Richardson, C. Bartling, Z. Guan, Non-enzymatically derived minor lipids found in Escherichia coli lipid extracts. Biochim Biophys Acta Mol Cell Biol Lipids 1811, 827–837 (2011) doi:10.1016/j.bbalip.2011.08.012.

24. H. Zhang, D. Zhang, K. Ray, M. Zhu, Mass defect filter technique and its applications to drug metabolite identification by high-resolution mass spectrometry. Journal of Mass Spectrometry 44, 999–1016 (2009) doi:10.1002/jms.1610.

25. E. Kendrick, A Mass Scale Based on CH2 = 14.0000 for High Resolution Mass Spectrometry of Organic Compounds. Anal Chem 35, 2146–2154 (1963) doi:10.1021/ac60206a048.

26. P. Zhang, W. Chan, I. L. Ang, R. Wei, M. M. T. Lam, K. M. K. Lei, T. C. W. Poon, Revisiting Fragmentation Reactions of Protonated α-Amino Acids by High-Resolution Electrospray Ionization Tandem Mass Spectrometry with Collision-Induced Dissociation. Sci Rep 9 (2019) doi:10.1038/s41598-019-42777-8.

27. C. Sohlenkamp, O. Geiger, Bacterial membrane lipids: Diversity in structures and pathways. FEMS Microbiol Rev 40, 133–159 (2015) doi:10.1093/femsre/fuv008.

28. O. Geiger, N. González-Silva, I. M. López-Lara, C. Sohlenkamp, Amino acid-containing membrane lipids in bacteria. Prog Lipid Res 49, 46–60 (2010) doi:10.1016/j.plipres.2009.08.002.

29. P. Lerouge, M.-H. Lebas, C. Agapakis-Causse, J.-C. Prome, Isolation and structural characterization of a new non-phosphorylated lipoamino acid from Mycobacterium phlei. Chem Phys Lipids 49, 161–166 (1988) doi:10.1016/0009-3084(88)90003-5.

30. C. P. Gill, C. Phan, V. Platt, D. Worrell, T. Andl, H. Roy, The MprF homolog LysX synthesizes lysyl-diacylglycerol contributing to antibiotic resistance and virulence. Microbiol Spectr 11, e0142929–23 (2023) doi:10.1128/spectrum.01429-23.

31. C. M. Ernst, A. Peschel, Broad-spectrum antimicrobial peptide resistance by MprF-mediated aminoacylation and flipping of phospholipids. Mol Microbiol 80, 290–299 (2011) doi:10.1111/j.1365-2958.2011.07576.x.

32. A. Peschel, R. W. Jack, M. Otto, L. V. Collins, P. Staubitz, G. Nicholson, H. Kalbacher, W. F. Nieuwenhuizen, G. Jung, A. Tarkowski, Kok, P. M., Van Kessel, J. A. G., Van Strijp, Staphylococcus aureus Resistance to Human Defensins and Evasion of Neutrophil Killing via the Novel Virulence Factor MprF Is Based on Modification of Membrane Lipids with L-Lysine. J. Exp. Med 193, 1067– 1076 (2001) doi:10.1084/jem.193.9.1067.

33. P. M. Christensen, J. Martin, A. Uppuluri, L. R. Joyce, Y. Wei, Z. Guan, F. Morcos, K. L. Palmer, Lipid discovery enabled by sequence statistics and machine learning. Elife 13 (2024) doi:10.7554/eLife.94929.

34. L. R. Joyce, H. S. Manzer, J. da C Mendonça, R. Villarreal, P. E. Nagao, K. S. Doran, K. L. Palmer, Z. Guan, Identification of a novel cationic glycolipid in Streptococcus agalactiae that contributes to brain entry and meningitis. PLoS Biol 20, e3001555 (2022) doi:10.1371/journal.pbio.3001555.

35. E. Maloney, D. Stankowska, J. Zhang, M. Fol, Q. J. Cheng, S. Lun, W. R. Bishai, M. Rajagopalan, D. Chatterjee, M. V. Madiraju, The two-domain LysX protein of Mycobacterium tuberculosis is required for production of lysinylated phosphatidylglycerol and resistance to cationic antimicrobial peptides. PLoS Pathog 5 (2009) doi:10.1371/journal.ppat.1000534.

36. F. Boldrin, L. Cioetto Mazzabò, M.-A. Lanéelle, L. Rindi, G. Segafreddo, A. Lemassu, G. Etienne, M. Conflitti, M. Daffé, A. Garzino Demo, R. Manganelli, H. Marrakchi, R. Provvedi, LysX2 is a Mycobacterium tuberculosis membrane protein with an extracytoplasmic MprF-like domain. BMC Microbiol 22, 85 (2022) doi:10.1186/s12866-022-02493-2.

37. J. M. Rock, F. F. Hopkins, A. Chavez, M. Diallo, M. R. Chase, E. R. Gerrick, J. R. Pritchard, G. M. Church, E. J. Rubin, C. M. Sassetti, D. Schnappinger, S. M. Fortune, Programmable transcriptional repression in mycobacteria using an orthogonal CRISPR interference platform. Nat Microbiol 2, 16274 (2017) doi:10.1038/nmicrobiol.2016.274.

38. A. M. Smith, J. S. Harrison, C. D. Grube, A. E. F. Sheppe, N. Sahara, R. Ishii, O. Nureki, H. Roy, tRNA-dependent alanylation of diacylglycerol and phosphatidylglycerol in Corynebacterium glutamicum. Mol Microbiol 98, 681–693 (2015) doi:10.1111/mmi.13150.

39. R. Vargas, M. J. Luna, L. Freschi, M. Marin, R. Froom, K. C. Murphy, E. A. Campbell, T. R. Ioerger, C. M. Sassetti, M. R. Farhat, Phase variation as a major mechanism of adaptation in Mycobacterium tuberculosis complex. Proc Natl Acad Sci U S A 120, e2301394120 (2023) doi:10.1073/pnas.2301394120.

40. H. Okuyama, T. Kankura, S. Nojima, Positional distribution of fatty acids in phospholipids from Mycobacteria. J Biochem 61, 732–7 (1967) doi:10.1093/oxfordjournals.jbchem.a128607.

41. M. Prithviraj, T. Kado, J. A. Mayfield, D. C. Young, A. D. Huang, D. Motooka, S. Nakamura, M. S. Siegrist, D. B. Moody, Y. S. Morita, Tuberculostearic Acid Controls Mycobacterial Membrane Compartmentalization. mBio 14, e0339622 (2023) doi:10.1128/mbio.03396-22.

42. E. Maloney, S. Lun, D. Stankowska, H. Guo, M. Rajagoapalan, W. R. Bishai, M. V Madiraju, Alterations in phospholipid catabolism in Mycobacterium tuberculosis lysX mutant. Front Microbiol 2, 19 (2011) doi:10.3389/fmicb.2011.00019.

43. D. M. Tobin, L. Ramakrishnan, Comparative pathogenesis of Mycobacterium marinum and Mycobacterium tuberculosis. Cell Microbiol 10, 1027–1039 (2008) doi:10.1111/j.1462-5822.2008.01133.x.

44. I. Mallick, P. Santucci, I. Poncin, V. Point, L. Kremer, J.-F. Cavalier, S. Canaan, Intrabacterial lipid inclusions in mycobacteria: unexpected key players in survival and pathogenesis? FEMS Microbiol Rev 45 (2021) doi:10.1093/femsre/fuab029.

45. M. S. Linz, A. Mattappallil, D. Finkel, D. Parker, Clinical Impact of Staphylococcus aureus Skin and Soft Tissue Infections. Antibiotics 12, 557 (2023) doi:10.3390/antibiotics12030557.

46. G. C. Monnot, M. Wegrecki, T. Y. Cheng, Y. L. Chen, B. N. Sallee, R. Chakravarthy, I. M. Karantza, S. Y. Tin, A. E. Khaleel, I. Monga, L. N. Uwakwe, A. Tillman, B. Cheng, S. Youssef, S. W. Ng, A. Shahine, J. A. Garcia-Vilas, A. C. Uhlemann, L. A. Bordone, A. Han, C. H. Rohde, G. Ogg, D. B. Moody, J. Rossjohn, A. de Jong, Staphylococcal phosphatidylglycerol antigens activate human T cells via CD1a. Nat Immunol 24, 110–122 (2023) doi:10.1038/s41590-022-01375-z.

47. G. J. Patti, O. Yanes, G. Siuzdak, Innovation: Metabolomics: the apogee of the omics trilogy. Nat Rev Mol Cell Biol 13, 263–9 (2012) doi:10.1038/nrm3314.

48. E. Stancliffe, G. J. Patti, PeakDetective: A Semisupervised Deep Learning-Based Approach for Peak Curation in Untargeted Metabolomics. Anal Chem 95, 9397–9403 (2023) doi:10.1021/acs.analchem.3c00764.

49. M. Blum, A. Andreeva, L. C. Florentino, S. R. Chuguransky, T. Grego, E. Hobbs, B. L. Pinto, A. Orr, T. Paysan-Lafosse, I. Ponamareva, G. A. Salazar, N. Bordin, P. Bork, A. Bridge, L. Colwell, J. Gough, D. H. Haft, I. Letunic, F. Llinares-López, A. Marchler-Bauer, L. Meng-Papaxanthos, H. Mi, D. A. Natale, C. A. Orengo, A. P. Pandurangan, D. Piovesan, C. Rivoire, C. J. A. Sigrist, N. Thanki, F. Thibaud-Nissen, P. D. Thomas, S. C. E. Tosatto, C. H. Wu, A. Bateman, InterPro: the protein sequence classification resource in 2025. Nucleic Acids Res 53, D444–D456 (2025) doi:10.1093/nar/gkae1082.

50. C. Vilchèze, J. Copeland, T. L. Keiser, T. Weisbrod, J. Washington, P. Jain, A. Malek, B. Weinrick, W. R. Jacobs, Rational design of biosafety level 2-approved, multidrug-resistant strains of Mycobacterium tuberculosis through nutrient auxotrophy. mBio 9 (2018) doi:10.1128/mBio.00938-18.

51. T. Parish, D. M. Roberts Editors, “Mycobacteria Protocols Third Edition Methods in Molecular Biology 1285;” http://www.springer.com/series/7651.

52. D. Kessner, M. Chambers, R. Burke, D. Agus, P. Mallick, ProteoWizard: Open source software for rapid proteomics tools development. Bioinformatics 24, 2534–2536 (2008) doi:10.1093/bioinformatics/btn323.

53. C. A. Smith, E. J. Want, G. O’Maille, R. Abagyan, G. Siuzdak, XCMS: Processing mass spectrometry data for metabolite profiling using nonlinear peak alignment, matching, and identification. Anal Chem 78, 779–787 (2006) doi:10.1021/ac051437y.

54. B. Bosch, M. A. DeJesus, N. C. Poulton, W. Zhang, C. A. Engelhart, A. Zaveri, S. Lavalette, N. Ruecker, C. Trujillo, J. B. Wallach, S. Li, S. Ehrt, B. T. Chait, D. Schnappinger, J. M. Rock, Genome-wide gene expression tuning reveals diverse vulnerabilities of M. tuberculosis. Cell 184, 4579–4592.e24 (2021) doi:10.1016/j.cell.2021.06.033.

55. M. van Kempen, S. S. Kim, C. Tumescheit, M. Mirdita, J. Lee, C. L. M. Gilchrist, J. Söding, M. Steinegger, Fast and accurate protein structure search with Foldseek. Nat Biotechnol 42, 243–246 (2024) doi:10.1038/s41587-023-01773-0.

56. F. Sievers, A. Wilm, D. Dineen, T. J. Gibson, K. Karplus, W. Li, R. Lopez, H. McWilliam, M. Remmert, J. Söding, J. D. Thompson, D. G. Higgins, Fast, scalable generation of high-quality protein multiple sequence alignments using Clustal Omega. Mol Syst Biol 7 (2011) doi:10.1038/msb.2011.75.

57. S. Kumar, G. Stecher, M. Suleski, M. Sanderford, S. Sharma, K. Tamura, MEGA12: Molecular Evolutionary Genetic Analysis version 12 for adaptive and green computing. Mol Biol Evol 41 (2024) doi:10.1093/molbev/msae263.

58. C. L. Cosma, L. E. Swaim, H. Volkman, L. Ramakrishnan, J. M. Davis, “Zebrafish and frog models of Mycobacterium marinum infection.” in Current Protocols in Microbiology (2006)vol. 3, pp. 10B.2.1–10B.2.33 doi:10.1002/0471729256.mc10b02s3.

